# Computationally-guided exchange of substrate selectivity motifs in a modular polyketide synthase acyltransferase

**DOI:** 10.1101/2020.04.23.058214

**Authors:** Edward Kalkreuter, Kyle S Bingham, Aaron M Keeler, Andrew N Lowell, Jennifer J. Schmidt, David H Sherman, Gavin J Williams

## Abstract

Acyltransferases (ATs) of modular polyketide synthases catalyze the installation of malonyl-CoA extenders into polyketide scaffolds. Subsequently, AT domains have been targeted extensively to site-selectively introduce various extenders into polyketides. Yet, a complete inventory of AT residues responsible for substrate selection has not been established, critically limiting the efficiency and scope of AT engineering. Here, molecular dynamics simulations were used to prioritize ~50 mutations in the active site of EryAT6 from erythromycin biosynthesis. Following detailed *in vitro* studies, 13 mutations across 10 residues were identified to significantly impact extender unit selectivity, including nine residues that were previously unassociated with AT specificity. Unique insights gained from the MD studies and the novel EryAT6 mutations led to identification of two previously unexplored structural motifs within the AT active site. Remarkably, exchanging both motifs in EryAT6 with those from ATs with unusual extender specificities provided chimeric PKS modules with expanded and inverted substrate specificity. Our enhanced understanding of AT substrate selectivity and application of this motif-swapping strategy is expected to advance our ability to engineer PKSs towards designer polyketides.

## INTRODUCTION

Type I polyketide synthases (PKSs) are giant multi-modular enzymes responsible for the biosynthesis of the scaffolds of many clinically relevant polyketides through the controlled stepwise assembly of a-substituted malonyl-CoA extender units.^1^ For canonical PKSs such as the prototypical 6-deoxyerythronolide B synthase (DEBS) from erythromycin biosynthesis, acyltransferase (AT) domains within each module are responsible for the selection of each extender unit and subsequent loading onto the cognate acyl carrier protein (ACP) (**Fig. 1a**-**b**). The ketosynthase (KS) domain from each module can then catalyze the decarboxylative condensation between the ACP-linked extender unit and the extended chain from the prior module.^2, 3, 4^ The most common extender units are malonyl-CoA (mCoA) and methylmalonyl-CoA (mmCoA, **1**), while ethylmalonyl-CoA (**2a**) and a few others are less frequently incorporated.^5^ A large portion of the polyketide structure derives from extender units, therefore contributing significantly to their potent biological activity and pharmacological properties.^6^ Accordingly, there is much interest in manipulating the extender unit selectivity of PKS modules to achieve the site-selective modification of polyketides. To this end, AT-swapping, motif-based chimeragenesis, and point mutagenesis have been explored to manipulate extender unit selection by PKSs.^6, 7, 8, 9, 10^ However, AT-swapped PKSs typically produce the desired polyketide in poor yields or are completely inactive, likely due to disruption of critical protein interactions and substrate channeling.^6^ Motif-swapping aims to exchange short sequences of residues that confer substrate selectivity. However, only a few conserved motifs have been identified and characterized, most notably the YASH motif (**Supplementary Figure S1**).^11^ Usually, replacement of the YASH motif produces only a modest change in extender unit specificity, indicating that motif-swapped hybrids are disrupted in some way and/or that the current motifs do not capture all of the specificity conferring residues.^10, 12^ For example, the active sites of AT2 and AT3 from the epothilone PKS differ by only nine residues, but the corresponding substrate selectivity is mCoA in AT2 and is relaxed towards mCoA/mmCoA in AT3. The YASH motif alone is not responsible for this difference, even though it accounts for two of the nine amino differences between the two active sites.^8, 13^ In contrast to exchanging residues between ATs, point mutations in or near the YASH motif of ATs of the DEBS (EryAT6) or pikromycin (PikAT5/PikAT6) pathways enable incorporation of non-natural extender units and significant changes to substrate selectivity without the deleterious effects of domain/module swapping.^14, 15, 16, 17, 18^ Yet, not all of these first-generation mutations are transferable between ATs in different PKSs.^18^ To be maximally efficient and broadly applicable, motif-swapping and point mutagenesis would benefit from a comprehensive inventory of AT residues responsible for substrate selection. Moreover, although the chemical diversity of naturally occurring extender units is somewhat limited, ATs that accept rare ones provide a potential template for engineering other PKSs to accept non-native or by extension, non-natural extender units.

**Fig. 1.**
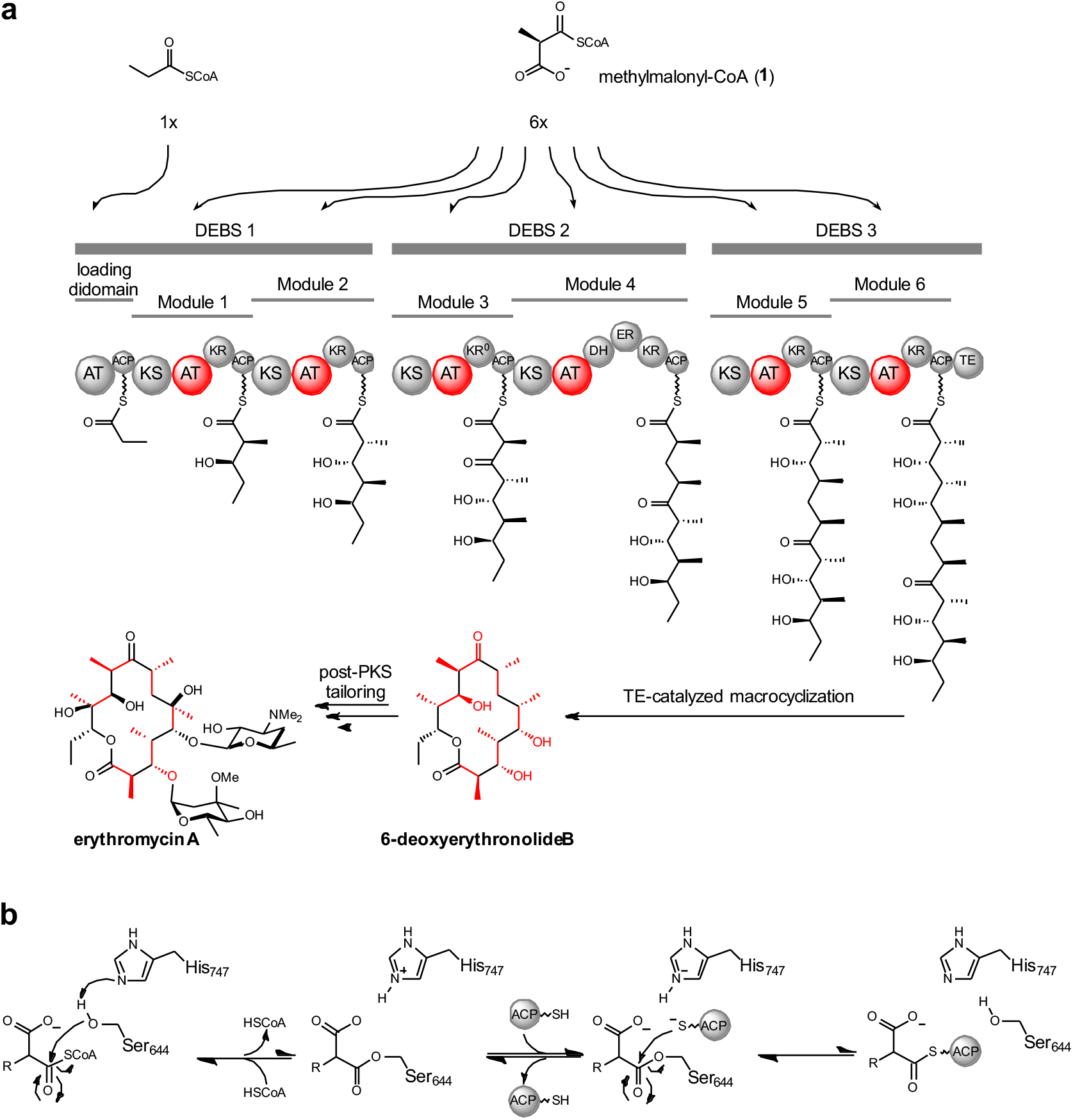
Organization of type I PKSs and mechanism of extender unit installation. (**a**) Domain and module organization of the prototypical type I PKS, DEBS. Each acyltransferase (AT) that selects an extender unit is shown in red, as is the contribution of each extender unit to the final polyketide structure. ACP, acyl carrier protein; KS, ketosynthase; KR, ketoreductase; DH, dehydratase; ER, enoylreductase; TE, thioesterase. (**b**) Abbreviated mechanism accounting for the installation of an extender unit onto the cognate ACP by EryAT6.

Here, a methodology guided by molecular dynamics (MD) simulations was employed to identify and mutate specificity-conferring residues within the AT domain, using EryAT6 as a model system. Collectively, these data informed additional predictions and enabled further exploration of a recently-identified ketoreductase (KR) domain bottleneck.^18^ Consequently, nine residues previously unassociated with AT substrate selectivity were identified. Insights from MD studies and analysis of EryAT6 mutations culminated in the prioritization of two previously unexplored structural motifs within the AT active site. Substitution of both motifs in EryAT6 with those from unrelated ATs with unusual extender unit specificities provided chimeric PKS modules with expanded and inverted substrate scope. These improvements in our understanding of AT substrate selectivity is expected to advance our ability to engineer PKSs towards an eventual goal of designer polyketides.

## RESULTS

### Molecular dynamics simulations of wild-type and first generation engineered EryAT6 variants

To obtain an enhanced understanding of how EryAT6 selects its extender unit for transfer to the ACP, MD simulations of the wild-type EryAT6 were carried out and compared to those containing previously identified substrate selectivity shifting mutations, V742A, Y744R, and L673H.^14, 16, 19^ The first two of these mutations shifts selectivity of Ery6TE towards propargylmalonyl-CoA while the third mutation shifts selectivity towards mCoA. A small panel of EryAT6 homology models was first constructed that included different combinations of the N- and C-terminal interdomain linkers with the core AT structure (**Supplementary Fig. S2**). These models were subjected to 10 ns MD simulations, and the simulations were analyzed for stability of secondary structure and maintenance of a reasonable distance between the catalytic dyad of Ser644 and His747, which are located on opposite sides of the hinge-like active site, within the large and small subunits of the AT, respectively (**Fig. 2a**). This distance was used to approximate the width of the active site and likely determines if the enzyme is in a catalytically competent state (**Fig. 2b**). The final model, with a 109-residue N-terminal KS-AT linker and a 43-residue C-terminal AT-KR linker flanking the 278-residue core AT (**Fig. 2a**), is in good agreement with an earlier study on AT boundaries (**Supplementary Table S1**).^12^ This wild-type EryAT6 model was solvated and simulated over 30 ns until structural convergence (**Supplementary Fig. S3a**). This structure was then used as the basis for all subsequent simulations, including the introduction of the individual mutations V742A, Y744R, and L673H, and the double mutation V742A/Y744R.^14, 16^ For each simulation trajectory, structural convergence (**Supplementary Fig. S3b-e**), the ‘dyad distance’ between the catalytic residues Ser644 and His747 (**Fig. 2b**), structural rearrangements (individual snapshots, **Fig. 2c** and **3**, **Supplementary Fig. S4**), and flexibility (root-mean-square fluctuation, RMSF, **Fig. 2d-f**, **Supplementary Fig. S5**) were inspected.

**Fig. 2.**
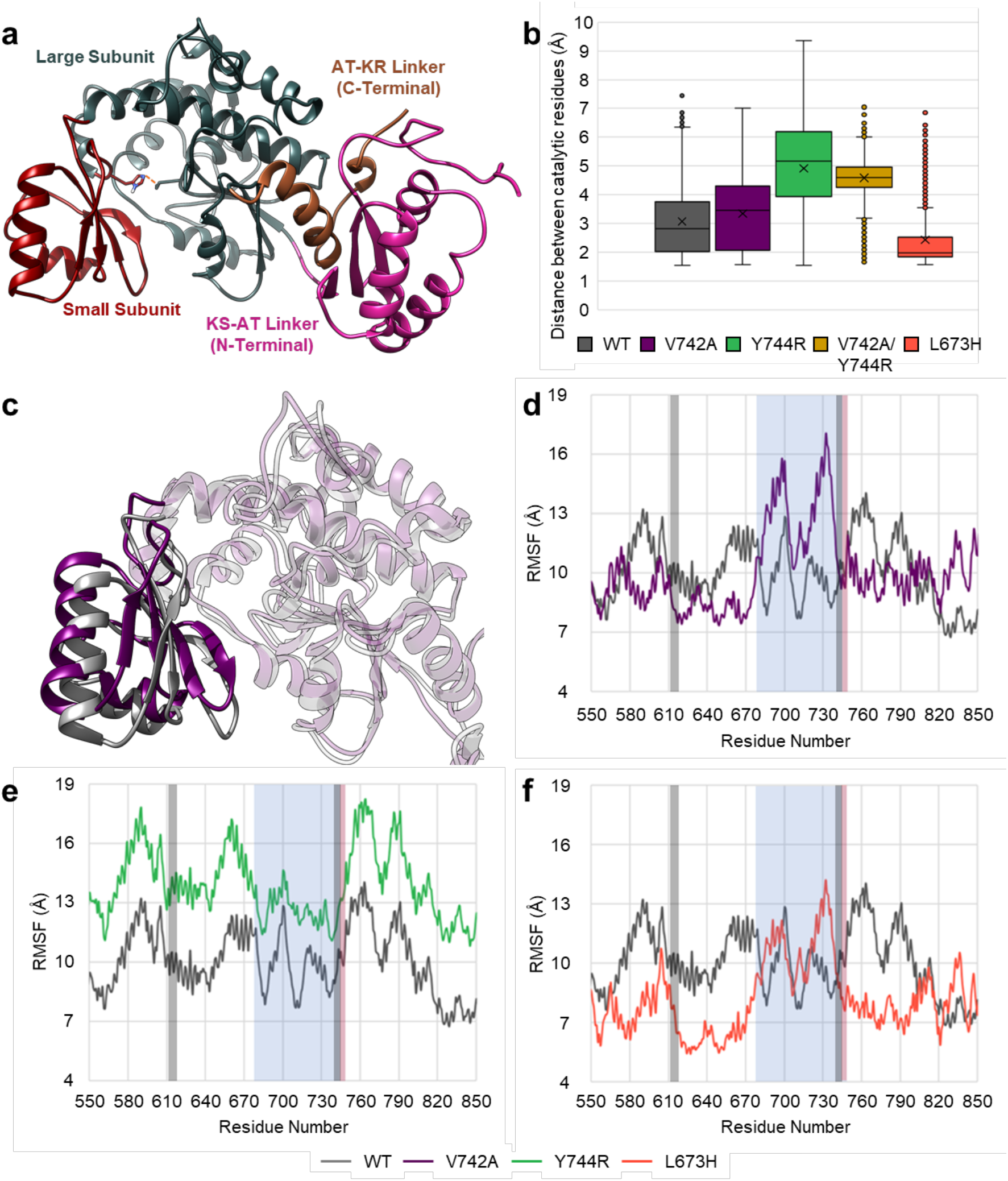
Molecular dynamic simulations of wild-type and variant EryAT6. (**a**) Final homology model of EryAT6 containing both linkers. The catalytic dyad is shown as sticks with the distance between indicated with a dashed orange line. (**b**) Box plot indicating the range of distances between the catalytic dyad over time in each MD simulation. (**c**) Overlay of a snapshot from each of the EryAT6 wild-type and V742A simulations highlighting the distortion of the small subunit. (**d-f**) Root-mean-square fluctuation (RMSF) of the 60 ns wild-type (grey) and the V742A (**d**, purple), Y744R (**e**, green), and L673H (**f**, orange) MD simulations. Higher values correspond to increased movement of a residue over the time frame. The blue box represents the residues found in the small subunit of the AT. The red box highlights the location of the YASH motif, and the grey boxes describe the locations of Motif-1 and Motif-2, including the YASH motif. Linkers were excluded from the analyses.

**Fig. 3.**
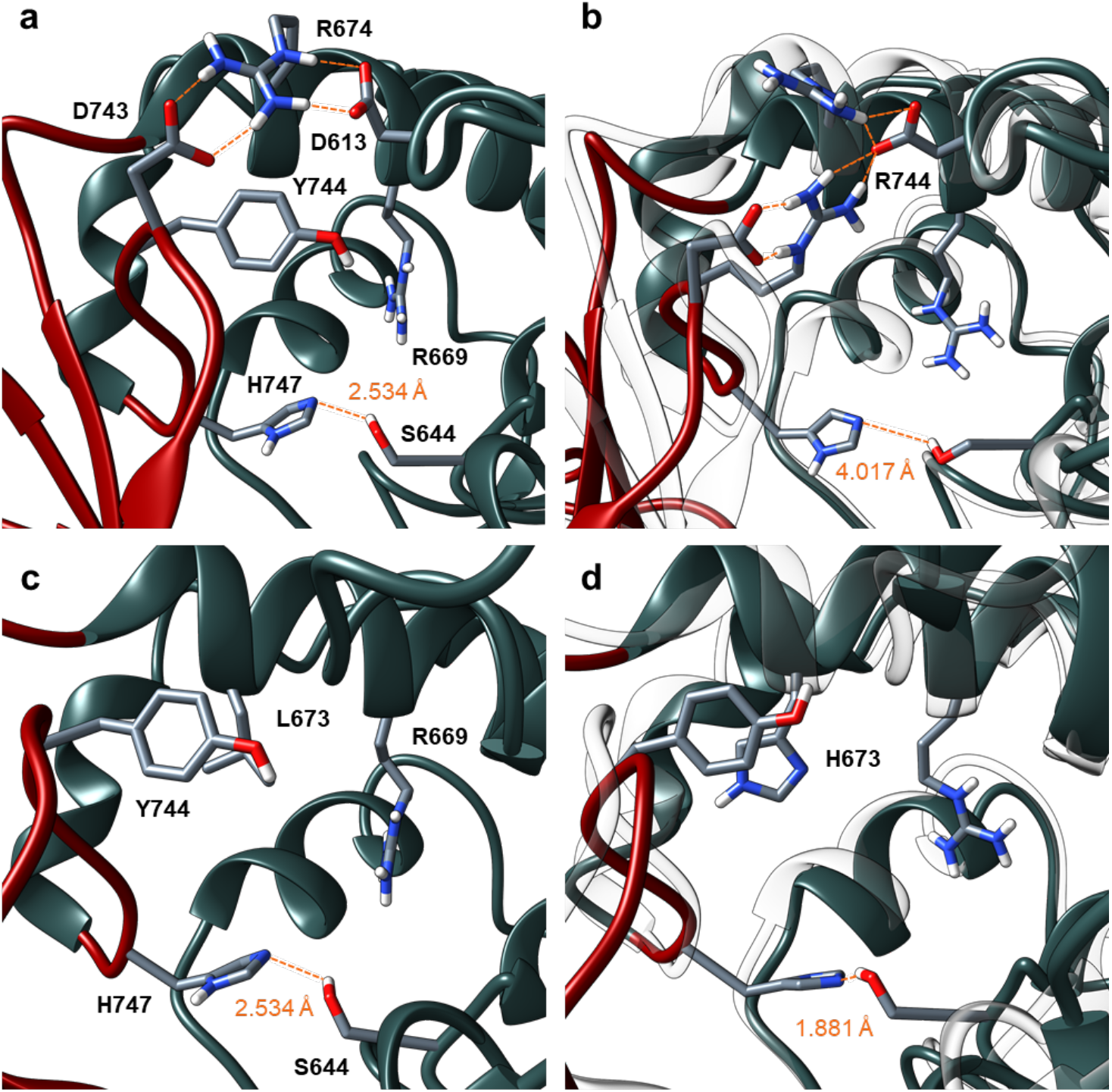
Snapshots from EryAT6 simulations showing effects of mutations on interactions between subunits. (**a**) Wild-type EryAT6 showing a critical salt bridge network involving Asp613, Arg674, and Asp743. (**b**) The Y744R mutation impacts the salt bridge, resulting in a larger inter-subunit distance. The wild-type model is overlaid as transparent ribbons. (**c**) Wild-type EryAT6 showing few interactions between Tyr744 and the large subunit. (**d**) Introduction of the L673H mutation results in a narrower active site via π-stacking between Tyr744 and His673. The wild-type model is overlaid as transparent ribbons.

Significant differences were observed when the distance between the catalytic dyad was measured over the entirety of the trajectories (**Fig. 2b**). For the wild-type enzyme, the distance between the two residues was <4 Å for >75% of the simulation (average of 3.07 Å). This result contrasted with the two mutations that inverted selectivity towards the larger propargyl substrate, Y744R and V742A/Y744R,^14^ as the dyad distance was >4 Å for ~75% of their simulations (average of 4.91 Å and 4.67 Å, respectively). The double mutant, which has poor *in vitro* activity, appeared unstable during its simulation and could not sustain dyad distances <3 Å across the active site. For V742A, which showed a modest shift towards propargyl,^14, 16^ the average dyad distance of the active site only increased 0.28 Å compared to wild-type. Finally, for the mmCoA-selective L673H mutant, the simulation depicted an active site that could not access the width required to accept the larger extenders (average distance of 2.43 Å).

Directly related to the variable dyad distances, the full-trajectory RMSF measurements highlight how the AT acts like a hinge, where the two subunits move closer or further apart, and where a single mutation may disrupt an entire subunit (**Fig. 2c**). In the V742A mutant, residues in the large subunit, including the YASH motif, remain mostly unaffected compared to the wild-type enzyme, but there are increases in movement across the small subunit, resulting in a deformed small subunit, likely due to the increased flexibility of alanine relative to valine in the active site loop (**Fig. 2cd**). Notably, the dramatic RMSF shift seen for the small subunit appears to exclude the YASH motif, indicating that instead, residues in the small subunit determine substrate selectivity. This conclusion is supported by previous examples of swapping the YASH motif.^8, 13, 20^

For the Y744R mutant, the entire enzyme undergoes more movement than the wild-type enzyme during simulations, while the small subunit retains its general shape (**Fig. 2e**). Unlike with the V742A mutant, the YASH motif itself sees a significant increase in flexibility relative to wildtype. This is likely due to both the disruption of an inter-subunit salt bridge between Asp743 and Arg674 (**Fig. 3a**) and subsequent formation of a new salt bridge with Arg744 on the same subunit (**Fig. 3b**), as well as the introduction of a bulky, positively charged residue into the active site YASH motif via Y744R.

In contrast, the L673H mutant replaces a hydrophobic leucine with a π-π stacking interaction between Tyr744 of the YASH motif and His673 of the large subunit (**Fig. 3c-d**). This mutation draws the two subunits closer together and reduces flexibility, resulting in a smaller active site and a more rigid structure overall (**Fig. 2f**). As with the two other mutations, L673H modulates the flexibility of the hinge-like architecture of the AT and the interactions between the two subunits.

Inclusion of substrate during the simulations was expected to result in a conformational change of the EryAT6 structure given the structural flexibility observed in the substrate-free simulations. The converged wild-type model was simulated with either mmCoA (**1**) or propargylmalonyl-CoA (**2b**) docked in the active site for 10 ns each to determine the residues likely involved in substrate binding and recognition (**Supplementary Fig. S6**). Consistent with previous structural studies,^16, 21, 22^ few residues appear to interact directly with the C2-sidechain of the CoA-linked substrate.

Outside of the YASH motif, however, the orientation of the **2b** propargyl side chain runs alongside the flexible small subunit loop (**Supplementary Fig. S6-S7**), emphasizing the important role of residues along this loop indicated by the RMSF plots of the V742A and Y744R mutants (**Fig. 2d-e**).

Interestingly, these simulations indicated that His643, located in the conserved GHSxG motif containing the catalytic Ser644, along with Gln560, may be responsible for positioning of the thioester near Ser644 (**Supplementary Fig. S6c**). This contrasts with previous suggestions that His643 is hydrogen bonded with the conserved Asn709 or stabilizes the malonyl-enzyme intermediate via water.^23^ To provide additional insight into the malonyl-enzyme intermediate, a simulation of wild-type EryAT6 with propargylmalonyl-Ser644 was carried out (**Supplementary Fig. S7**), revealing the disruption of the salt bridge network between Asp743 of the small subunit and Arg674 and Asp613 of the large subunit. In contrast to the malonyl-enzyme intermediate, this network is retained in ~50% of the MD trajectory frames for substrate-free EryAT6 variants, except for Y744R, wherein it is mostly disrupted by Arg744 as described (**Supplementary Fig. S8**). With the smaller native substrate, this network may help to preclude water from entering the active site and hydrolyzing the substrate.

Together, these insights suggest that a large portion of the AT active site contributes to extender unit specificity and activity by impacting (1) subunit interactions, (2) critical salt-bridges, and (3) catalytic dyad distance. Moreover, it sheds light on networks of interacting residues spanning both subunits of the AT.

### Probing the extender unit selectivity of wild-type Ery6TE via a multi-substrate competition assay

Next, the *in vitro* extender unit selectivity of wild-type Ery6TE was determined using a pool of competing equimolar extender units (300 μM each of **1**, **2a**-**d**, **Fig. 4a** and **Supplementary Fig. S9**) and a previously-described synthetic thiophenol-pentaketide substrate (**3**) based on the pikromycin PKS pathway.^14, 18, 24^ An NADPH recycling system was included to support KR activity and production of the 10-deoxymethynolide (10-dML) series **4** and **5a**-**d** upon extender unit installation and cyclization. The corresponding non-reduced series **6a**-**d** are possible if the KR does not process the nascent β-carbonyl, as reported previously.^18^ Subsequently, the percentage distribution of the products derived from each extender unit was determined by high-resolution LC-MS analysis. As anticipated, the major product in the wild-type Ery6TE competition assay was derived from the native extender **1** (**4**, 58%), followed by those derived from the propargyl-**2b** (27%), isopropyl-**2c** (11%), ethyl-**2a** (3%), and butyl-**2d** (1%) extenders (**Table 1** and **Supplementary Tables S2**-**S3**). Although wild-type Ery6TE displays significant *in vitro* extender unit promiscuity, some extenders are poorly utilized (e.g. **2b**, **2d**). Nevertheless, this baseline promiscuity serves as a good starting point to manipulate the extender unit specificity by protein engineering.

**Fig. 4.**
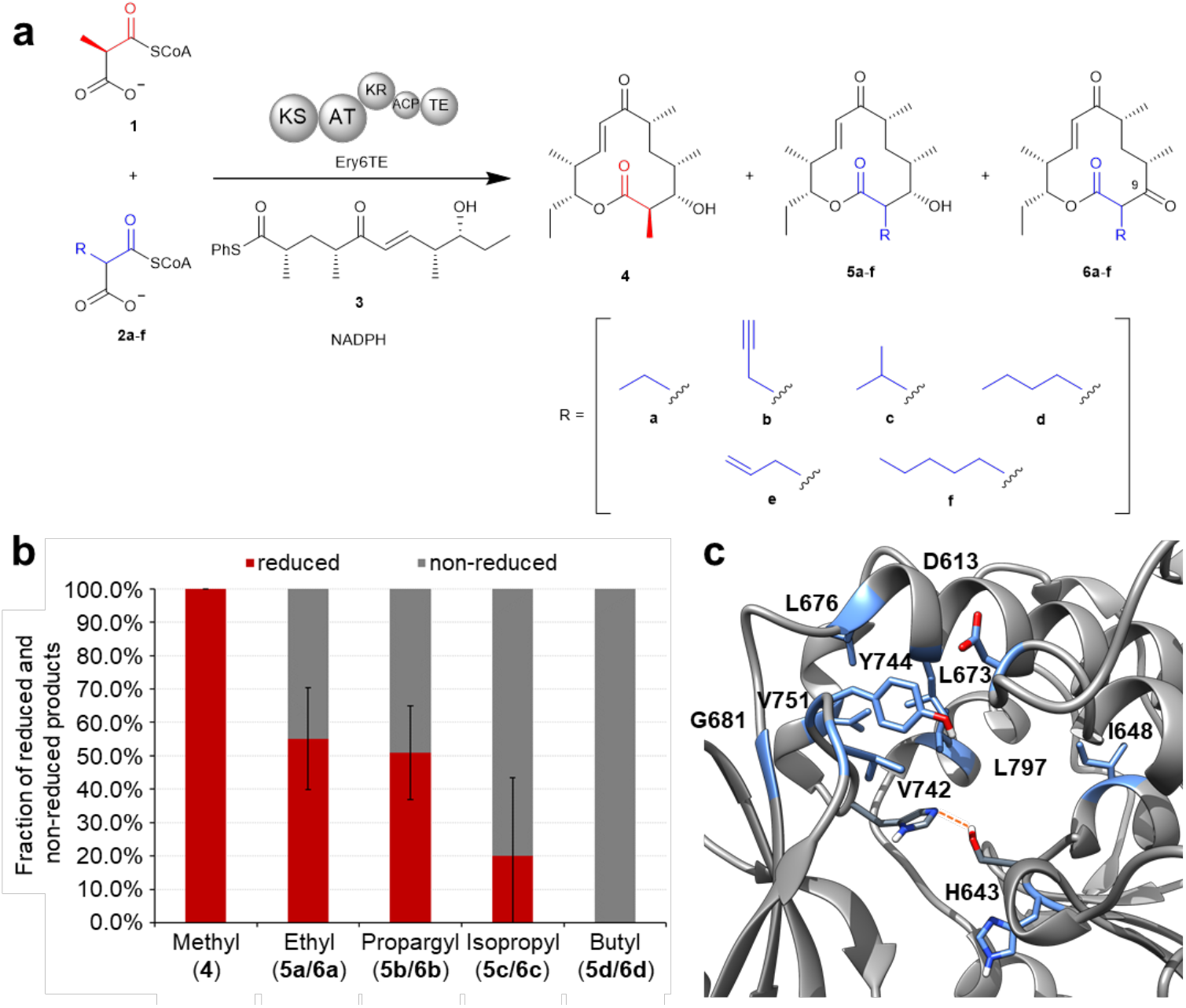
Ery6TE multi-substrate competition assay. (**a**) The Ery6TE module is incubated with the synthetic pentaketide chain mimic **3** and an equimolar mixture of the native extender **1** and mixtures of **2a**-**f** *in vitro*. Products **4** and **5a**-**f** are produced upon AT-catalyzed extender unit installation, KS-extension, KR-reduction, and cyclization by the TE. In contrast, **6a**-**f** are produced when the KR does not process the C-9 keto. NADPH was produced *in situ* with an NADPH regeneration system (not shown). (**b**) The fraction of each reduced (**4**, **5a**-**d**) and non-reduced (**6a**-**d**) product (total = 100) catalyzed by wild-type Ery6TE as determined by HR LC-MS analysis. Error bars are the standard deviation (*n* = 3 biological replicates) of the average fraction of the non-reduced (**6a**-**6d**) product. (**c**) The active site of wild-type EryAT6 showing key residues (in blue) that were targeted for individual mutations and successfully affected substrate selectivity.

**Table 1.** Product distributions catalyzed by wild-type and variant Ery6TE in the presence of competing equimolar 300 μM extender units.^*a*^

| Ery6TE Variant | **4<sup>b</sup>** | **5a + 6a<sup>b</sup>** | **5b + 6b<sup>b</sup>** | **5c + 6c<sup>b</sup>** | **5d + 6d<sup>b</sup>** | Relative Activity<sup>c</sup> |
| --- | --- | --- | --- | --- | --- | --- |
| WT | 57.7 ± 2.2% | 2.9 ± 0.2% | 27.1 ± 0.9% | 11.4 ± 0.4% | 1.0 ± 0.0% | 100.0 |
| D613E | 56.4 ± 4.6% | 2.5 ± 0.4% | 25.9 ± 3.1% | 13.4 ± 1.1% | 1.9 ± 0.0% | 101.9 ± 13.3 |
| H643F | 95.0 ± 2.9% | 1.1 ± 0.0% | 1.9 ± 0.2% | 1.7 ± 0.2% | 0.3 ± 0.0% | 153.5 ± 4.3 |
| H643S | 52.9 ± 2.7% | 4.5 ± 0.1% | 25.5 ± 2.3% | 13.1 ± 1.5% | 4.1 ± 0.1% | 57.6 ± 7.0 |
| I648G | 96.7 ± 2.1% | 0.4 ± 0.1% | 0.8 ± 0.1% | 1.7 ± 0.2% | 0.4 ± 0.0% | 180.8 ± 12.8 |
| L673G | 65.7 ± 2.2% | 2.5 ± 0.4% | 22.9 ± 1.6% | 6.5 ± 0.9% | 2.4 ± 0.2% | 152.6 ± 17.6 |
| L676V | 73.4 ± 1.6% | 3.1 ± 0.1% | 7.8 ± 0.9% | 11.7 ± 1.0% | 4.1 ± 0.5% | 135.1 ± 23.9 |
| G681A | 59.5 ± 2.0% | 2.7 ± 0.4% | 18.6 ± 0.8% | 17.8 ± 1.2% | 1.5 ± 0.0% | 124.6 ± 22.6 |
| V742A | 56.0 ± 2.4% | 4.3 ± 0.2% | 35.6 ± 2.4% | 2.4 ± 0.2% | 1.7 ± 0.2% | 76.8 ± 1.8 |
| Y744R | 17.7 ± 1.2% | 2.5 ± 0.1% | 57.0 ± 3.2% | 13.0 ± 1.9% | 9.7 ± 0.2% | 46.2 ± 4.1 |
| V751A | 54.3 ± 2.8% | 2.7 ± 0.3% | 29.7 ± 2.3% | 10.7 ± 0.2% | 2.5 ± 0.2% | 76.5 ± 3.7 |
| L797E | 98.4 ± 1.3% | 0.5 ± 0.1% | 0.5 ± 0.0% | 0.3 ± 0.0% | 0.3 ± 0.0% | 259.6 ± 47.0 |
| Ery6(Cin1)TE | 16.3 ± 0.2% | 0.6 ± 0.1% | 55.8 ± 5.9% | 6.9 ± 0.5% | 20.3 ± 2.7% | 35.9 ± 5.7 |
| Ery6(Tha13)TE | 11.2 ± 0.2% | 0.9 ± 0.0% | 18.3 ± 0.8% | 23.2 ± 3.0% | 46.5 ± 2.9% | 30.1 ± 1.4 |

<sup>a</sup> Module lysates were incubated with 500 μM each of **1**, **2a**, **2b**, **2c**, and **2d** simultaneously. Product distributions of mutants that could not be distinguished from wild-type Ery6TE are omitted for clarity.
<sup>b</sup> Percent of each indicated product out of all products (**4**, **5a**, **6a**, **5b**, **6b**, **5c**, **6c**, **5d**, and **6d**). Error bars are the standard deviation ( $n = 3$ biological replicates) of the mean. See Figure 3a for structures of products.
<sup>c</sup> Total activity of each system relative to wild-type Ery6TE is based on total amount of products and normalized to amount of enzyme (**4**, **5a**, **6a**, **5b**, **6b**, **5c**, **6c**, **5d**, and **6d**).

Notably, while wild-type Ery6TE appears to discriminate against the largest extender (butyl) and the extenders most similar to the native extender (ethyl and malonyl, data not shown), the fraction of the non-reduced product correlated to the size of the extender unit C-2 side-chain (**Fig. 4b**). For example, 100% of the product is reduced when the natural substrate **1** is utilized, while 100% of the product is non-reduced when derived from the butyl substrate **2d** (**Fig. 4b**), in a similar fashion to KR5 in an engineered pikromycin module.^18^ While the KR likely plays some role in determining whether a given extender unit is converted into a final cyclized product by Ery6TE, clearly the specificity restrictions of the AT needs to be overcome first in order to provide the KR the opportunity to carry out both native and non-native C-9 reductions.

### Extensive mutagenesis of the Ery6AT active site

To test the hypothesis from the MD simulations that a large portion of residues in the AT active site contributes to extender unit selectivity, mutations at residues outside the YASH and other established motifs were designed by inspection of the EryAT6 model and simulations (**Fig. 4c** and **Supplementary Table S4**). Briefly, mutations were designed to influence extender unit selectivity based on predicted impact on subunit interactions and flexibility, critical salt-bridges, and active site size. In total, 32 mutants were constructed, spanning 26 residues. Included for comparison were three previously reported mutations: V742A, Y744G, and Y744R.^61, 143^ Val742 and Tyr744 are located near and in the conserved YASH motif, respectively (**Fig. 4c**). Each mutant was probed using the multi-substrate competition assay as described for the wild-type enzyme (**Table 1** and **Fig. 4c**).

For the first-generation mutants, the largest change in product distribution for V742A was the ethyl- (**5a**+**6a**) and butyl-macrolactones (**5d**+**6d**), with a 1.5- and 2.0-fold improvement compared to the wild-type, respectively. Nevertheless, similar to wild-type Ery6TE, the methyl-derived product (**4**) was the major product for V742A. Introduction of Y744R led to a 2- and 10-fold improved production of the propargyl- (**5b**+**6b**) and butyl-macrolactones (**5d**+**6d**), respectively, with the propargyl-macrolactones (**5b**+**6b**) being the major products. The product distributions of 21 mutants could not be distinguished from that of the wild-type (**Supplementary Table S4**). However, 8 designed mutants (H643F, H643S, I648G, L673G, L676V, G681A, V751A, L797E) led to changes in product distributions, indicating that their substrate selectivity’s were altered (**Table 1**). Although the **1**-derived macrolactone **4** was the major product for each of these single mutants, some supported up to ~4-fold increase in the proportion of products derived from nonnative extenders. For example, the butyl products (**5d**+**6d**) were increased by 2.4-, 4.1-, 2.5-, and 1.9-fold with L673G, L676V, V751A, and D613E, respectively, compared to wild-type. Interestingly, substitution at His643 (**Fig. 4c**), which we propose to position the thioester of the malonyl-CoA substrate for catalysis, with serine or phenylalanine providing opposite effects on the product distributions. H643S improved the fraction of the ethyl (**5a**+**6a**) and butyl (**5d**+**6d**) products 1.6- and 4.1-fold, respectively, compared to the wild-type, while 95% of the products of the H643F mutant were from the native substrate **1**, and this mutant was more stringent than the wild-type (**Table 1**).

Two other **1**-specific mutations were identified. I648G and L797E both resulted in >95% methyl product **4** (**Table 1**). I648G is located outside the active site within the large subunit (**Fig. 4c**) and is proposed to increase the flexibility of the conserved GHSxG loop and thus shorten the distance between the catalytic dyad. In contrast, the introduction of charge directly behind the hydrophobic Leu673 via L797E likely restricts space in the rear of the active site.

To summarize, mutations at five previously untargeted residues (H643S, L673G, L676V, G681A, V751A) impact selectivity for larger extenders, and mutations at three additional novel residues (H643F, I648G, L797E) increase specificity for the native substrate **1**. While only one of these residues (His643) likely directly interacts with the substrate, the overall findings highlight that residues in both subunits of the AT play a role in dictating specificity, albeit making minor contributions individually. As seen with the RMSF analysis (**Fig. 2d-f**), single mutations can cause significant structural changes throughout the active site. Accordingly, because most of these mutations implicate molecular networks outside the established YASH motif, a combination of rationally-designed substitutions at multiple contiguous sites could realize more extreme selectivity shifts.

### Introduction of active site motifs to invert extender unit selectivity of Ery6TE

The large conformational changes observed by MD simulations, coupled with the established impact of mutations throughout the entire AT active site, suggest that targeting structural motifs could be more effective than single mutations by compensating for otherwise deleterious effects and maximizing synergy between subtle amino acid variations. Most notably, a loop in the small subunit spanning Thr739 to His747 (including the YASH motif) in Ery6TE, defined here as the small subunit motif (SSM; **Fig. 5a**,**d**), is the region that contains the most direct interactions with the substrate (**Supplementary Fig. S6-7**). Additionally, MD simulations implicated mutations within this loop (especially V742A and Y744R), to impact substrate specificity (**Fig. 2**). Mutations at the nearby Gly681, Val751, Leu673, and Leu676, suggest that these residues likely interact with this loop.

**Fig. 5.**
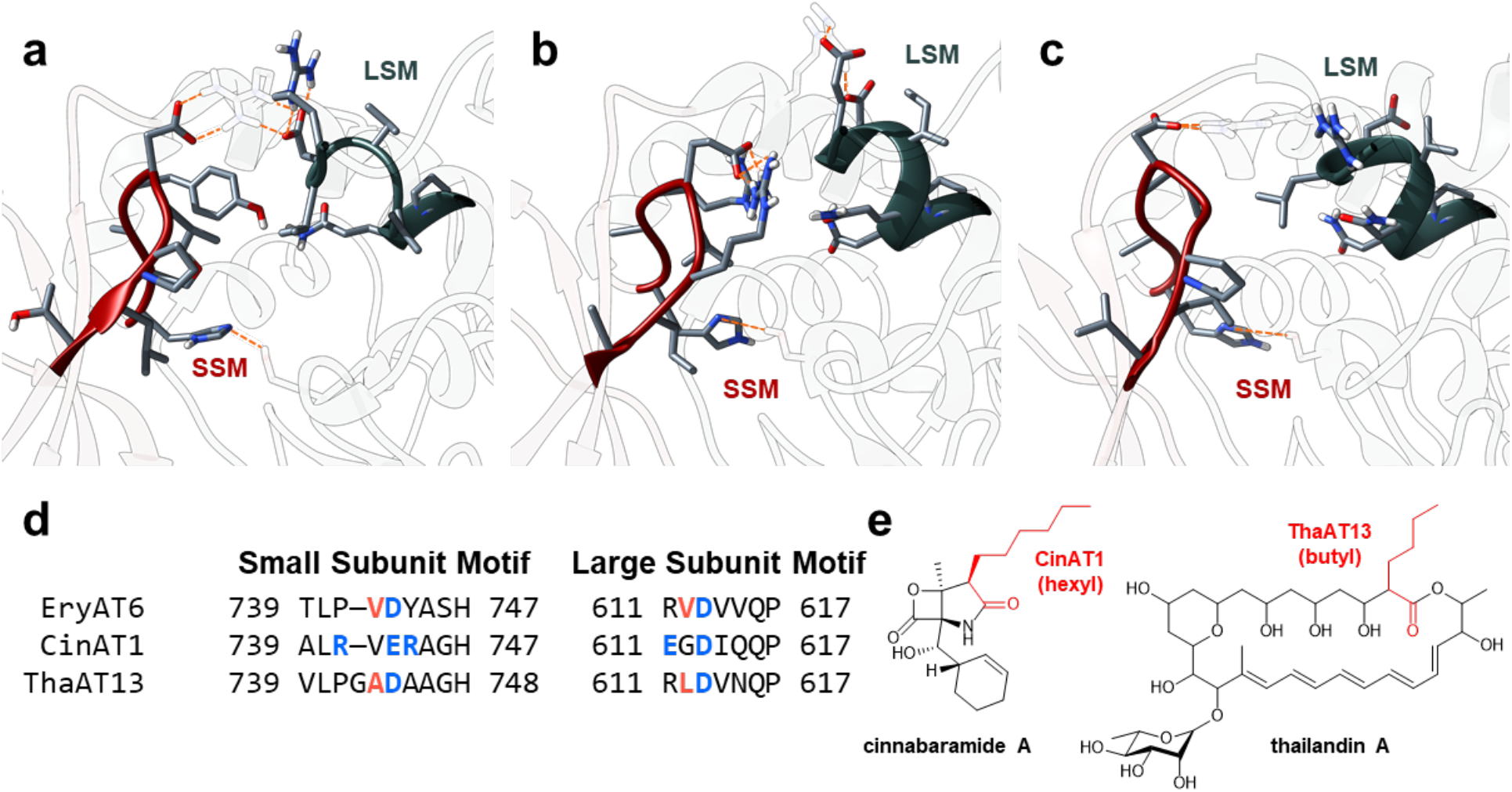
EryAT6 active site and chimeragenesis. The active sites of wild-type (**a**), Cin1 (**b**), and Tha13 (**c**) EryAT6 homology models based on the molecular dynamics-generated wild-type model. The two structural motifs are represented by sticks and highlight the interactions between the large subunit (blue grey) and small subunit (red). The catalytic dyad Ser644 and His747 are shown for reference. (**d**) Amino acid alignment of EryAT6, CinAT1, and ThaAT13, showing the identity of residues in motif-1 and motif-2. Indicated residues are expected to interact with the opposite motif through hydrophobic (orange) and charge-mediated (blue) interactions. (**e**) Structures of the natural products biosynthesized by the cinnabaramide and thailandin PKS. Portions of the structures dictated by CinAT1 and ThaAT13 are highlighted in red.

While the entirety of the SSM is not conserved by substrate specificity (**Supplementary Fig. S10**), it spans the full length of the active site, from the active site opening through to the back of the pocket, and therefore if targeted for motif swapping would be expected to retain structural integrity while being able to better accommodate the CoA-linked substrate not only once bound, but also during its approach into the pocket. In addition, a second motif dubbed the large subunit motif (LSM), makes up the portion of the helix directly above the catalytic Ser644 in the large subunit, spanning from Arg611 to Pro617 in EryAT6, and contains the well-conserved mmCoA-specific VDVVQP residues (**Fig. 5a**,**d**, **Supplementary Fig. S10b**).^17, 25^ Residues within this motif have been previously implicated in substrate selectivity, with the Q616H mutation increasing promiscuity in EryAT6 and a Phe residue shown to π-stack with a phenylmalonyl-CoA substrate in the splenocin PKS.^17, 25^ Based on the enzyme-substrate simulations (**Supplementary Fig. S6-S7**), the LSM does not have any obvious interactions with the C2-sidechain of most substrates. However, in several of the wild-type and mutant MD trajectories, with and without malonyl-CoAs, residues from the SSM and the LSM interact directly or through Arg674 to form a “closed” conformation of the hinge-like AT (**Fig. 3a** and **Supplementary Fig. S4, S6-S7**, **S11**). In support of these predicted interactions, residue differences in one motif often correspond to complementary changes in the other motif in many ATs of varying substrate specificities (**Supplementary Fig. S10a**). Therefore, the motifs are implicated as potential paired exchangeable structural unit that could be targeted to manipulate extender unit selection.

To test this hypothesis, the SSM and the LSM in EryAT6 were replaced with the corresponding residues from two unrelated ATs with different extender unit specificities (**Fig. 5b**-**e**), the hexylmalonyl-CoA-selective AT1 from the cinnabaramide PKS in *Streptomyces cinnabarigriseus* (CinAT1) and the butylmalonyl-CoA-selective AT13 from the thailandin PKS in *Actinokineospora bangkokensis* (ThaAT13),^26, 27^ producing two chimeras, Ery6(Cin1)TE and Ery6(Tha13)TE, respectively. Each chimera now differs from the wild-type Ery6TE at nine and seven residues, respectively.

Each parent AT has one or more distinctive features in the two motifs, which are predicted to play major roles in their respective substrate selectivities. CinAT1 represents one of the rare natural ATs containing the equivalent Y744R mutation within the YASH motif, but it also introduces a second arginine into the SSM (**Fig. 5b**,**d**). Despite suggestions that an arginine at position 744 prevents efficient use of extenders much larger than propargyl,^17^ CinAT1 uses larger extenders natively.^26^ The two flexible and positively charged arginines are positioned to be compensated by one or both of two nearby glutamates, one in each motif (**Fig. 5b,d**). These charge-mediated interactions may replace the largely hydrophobic interactions between the two motifs in EryAT6 (Val742 and Val612, **Fig. 5a**,**d**) and ThaAT13 (Ala743 and Leu612, **Fig. 5c**-**d**). The additional methylene group of Glu743 in the CinAT1 SSM also provides the side chain length required to interact with both arginines and hold them out of the active site channel. This arrangement contrasts with that of Arg744 in the Y744R single mutant in EryAT6, where it is instead held in place between the two subunits (**Fig. 3b**) and likely contributes to additional flexibility of the small subunit loop. ThaAT13 has a similar LSM to EryAT6, but in its SSM, five of the ten residues are either alanine or glycine, resulting in a significantly smaller and more flexible loop than that found in EryAT6. In addition to AAGH replacing the YASH motif, Val742 in EryAT6 has been replaced with a glycine and an alanine in ThaAT13, providing an extra residue and increased flexibility and space within the active site (**Supplementary Fig. S10-S11**). Unlike in the unstable V742A/Y744R mutant, the Tha13 motifs cooperatively compensate for the smaller residue (Ala743) via the extra length of ThaAT13’s Leu612 relative to EryAT6’s Val612, maintaining the stabilizing hydrophobic interaction that is replaced by charge-mediated interactions in CinAT1 (**Supplementary Fig. S11**).

Based on this logic, the extender unit selectivity of the two motif-swapped Ery6TE chimeras was probed with the multi-substrate competition assay, revealing a large shift in preference towards larger extender units and mirroring the selectivity expected from the parent wild-type ATs (**Fig. 6**, **Table 1**, and **Supplementary Table S3**). For example, the propargyl (**5b**+**6b**) and butyl (**5d**+**6d**) products increased 2- and 20-fold, respectively, in the Ery6(Cin1)TE-catalyzed reaction, compared to wild-type Ery6TE. Concomitantly, the fraction of the methyl (**4**), ethyl (**5a**+**6a**), and isopropyl (**5c**+**6c**) products decreased 3.5-, 4.8-, and 1.7-fold, respectively, indicating a shift in substrate selectivity towards the propargyl- and butyl-extender units. Similarly, the Ery6(Tha13)TE chimera also provided a shift in selectivity towards the larger extender units, but the overall change is more pronounced. For instance, the fraction of the isopropyl (**5c**+**6c**) and butyl (**5d**+**6d**) products increased 2- and 47-fold, respectively, compared to wild-type Ery6TE, while that of the methyl (**4**), ethyl (**5a**+**6a**), and propargyl (**5b**+**6b**) products decreased 5-, 3.2-, and 1.5-fold, respectively, compared to wild-type Ery6TE. Subsequently, the product distribution of the Ery6(Tha13)TE chimera mirrors what is expected from the use of the motif from the butyl-**2d** utilizing ThaAT13. Interestingly, the optimal extender units for the Ery6(Cin1)TE chimera or CinAT1 are not yet known, propargyl (**5b**+**6b**) production by Ery6(Cin1)TE is indistinguishable from that of Y744R (**Fig. 6**), while the fraction of the butyl product (**5d**+**6d**) supported by this chimera is more than double that of the single mutant Y744R. For comparison, the methyl-specific I648G mutant inspired by the MD simulations was included (**Fig. 6**) to emphasize the orthogonal AT substrate selectivities expected of the designed AT mutants. Crucially, in addition to the expected dramatic shifts in extender unit selectivities displayed by the motif-swapped chimeras, the relative activities compared to the wild-type Ery6TE of both chimeras are robust (>30%, **Table 1**), often higher than individual or double mutants (e.g., V742A/Y744R).^14^

**Fig. 6.**
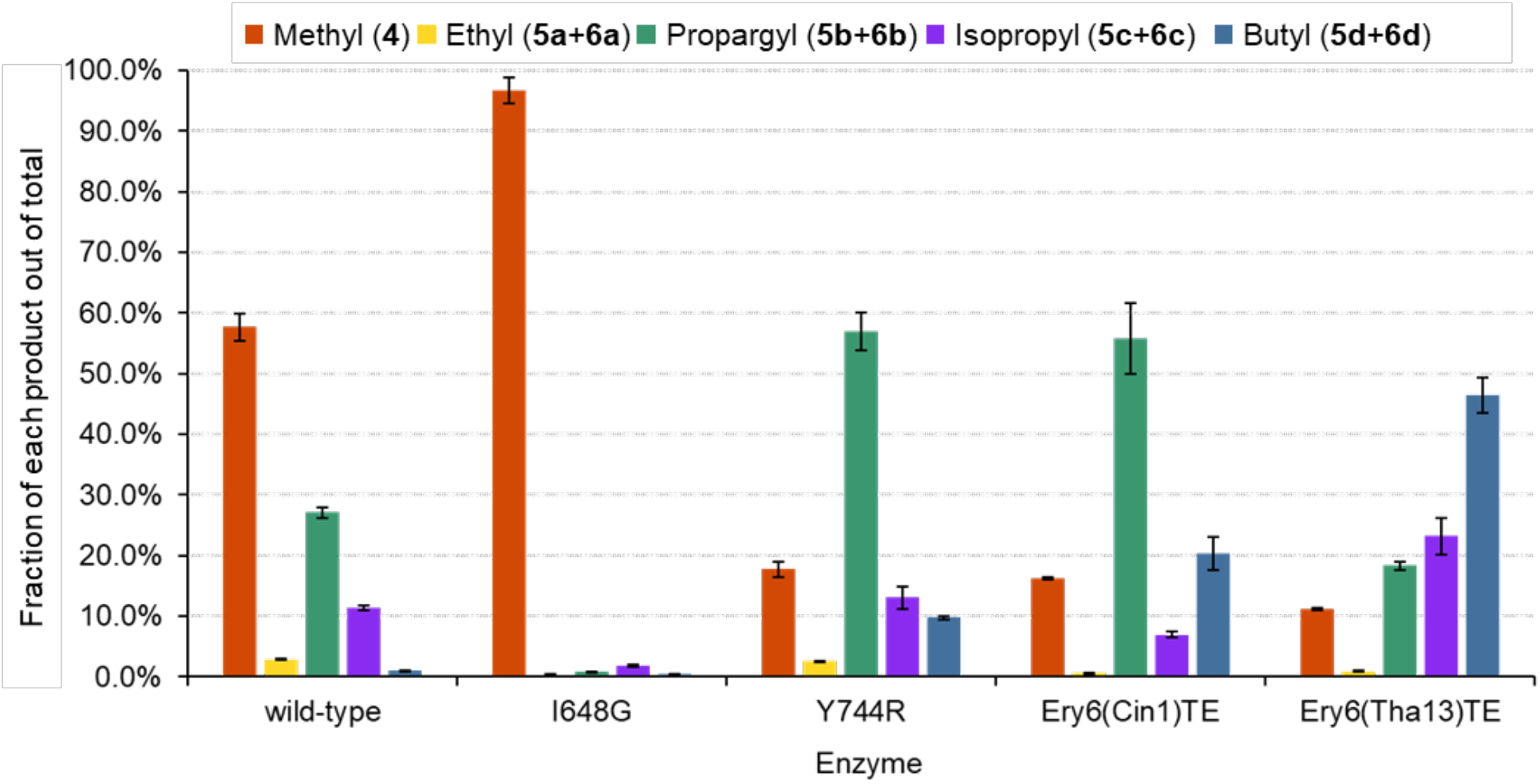
Product distributions catalyzed by motif-swapped Ery6TE variants with five competing substrates. Motif-swapped and selected individual mutant module lysates were incubated with 500 μM each of **1**, **2a**, **2b**, **2c**, and **2d**, simultaneously. Products were analyzed by HR-LCMS, and the product distributions are shown. Error bars are the standard deviation (*n* = 3 biological replicates) of the mean.

Next, to better mimic expected *in vivo* conditions in an engineered strain,^28^ the product distribution catalyzed by the wild-type Ery6TE, I648G, Y744R, and the two motif-swapped chimeras, was determined using high concentrations (1.5 mM) of the native Ery6TE substrate (**1**) in competition with just one other extender unit (either **2b**, **2d**, **2e**, or **2f**, **Fig. 4a**). Notably, under these conditions, the wild-type Ery6TE is remarkably stringent. Only ≤ 10% of the products are derived from **2b**, **2d**, **2e**, or **2f**, respectively, in each reaction (**Fig. 7** and **Supplementary Table S5**), indicating that the **1**-derived macrolactones are the major products. As expected, Y744R inverted the AT selectivity towards the propargyl products (**5b**+**6b**), but the single mutation could not invert the substrate preference from methyl to any of the other three extender units tested. The I648G mutation proved even more stringent for the native extender **1** under these conditions, with only 0.3% incorporation of **2b** and no incorporation of any of the remaining extenders. In contrast, >70% of the products catalyzed by the Ery6(Tha13)TE and Ery6(Cin1)TE chimeras were the propargyl (**5b**+**6b**), butyl (**5d**+**6d**), and pentyl (**5f**+**6f**) macrolactones in each competition assay,indicating a complete inversion of substrate selectivity toward each corresponding extender unit over the native methyl substrate **1**. Notably, for Ery6(Tha13)TE, the pentyl keto analogue (**6f**) comprised 93.5% of the total products when **2f** was in direct competition with the native extender unit **1**, a ~300-fold increase in substrate selectivity as compared to wild-type Ery6TE. As expected from analysis of the distribution of reduced and non-reduced products with **2a**-**2d** and our prior studies with the pikromycin PKS, most of the allyl products were not reduced while none of the pentyl products had been processed by the KR (**Supplementary Table S5**).^18^

**Fig. 7:**
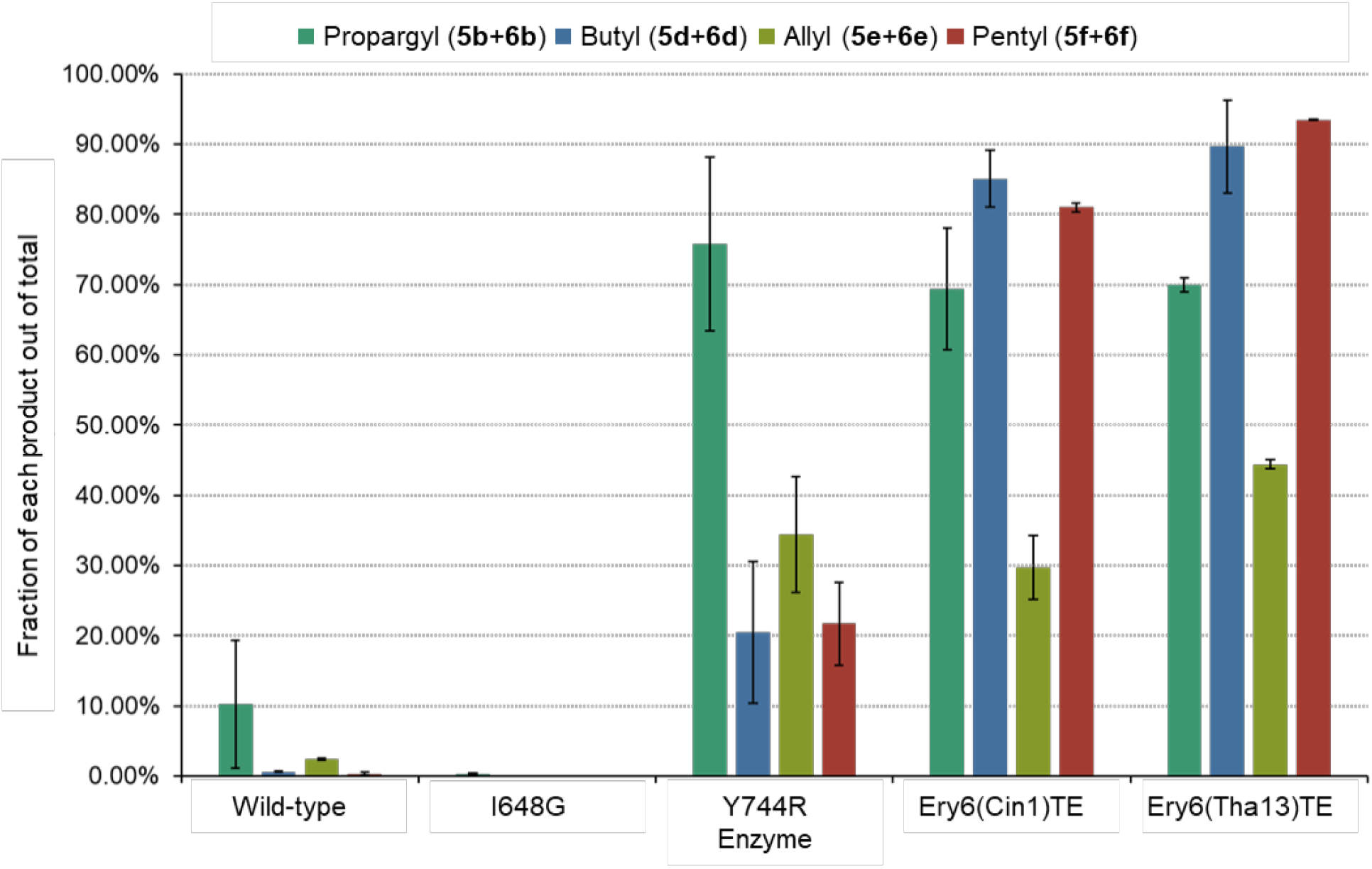
Product distributions catalyzed by motif-swapped Ery6TE variants with two competing substrates. Ery6TE extension unit competition assay with 1.5 mM **1** and 1.5 mM each of either **2b**, **2d**, **2e**, or **2f**. Average percentage of each indicated product out of the total products in each reaction as determined by HR-LCMS analysis. Error bars are the standard deviation (*n = 3* biological replicates) of the mean.

## DISCUSSION

Engineering PKSs to accept non-native extender units is a necessary step towards diversifying bioactive polyketide core structures. To date, there have been several advances towards introduction of non-natural moieties using native AT promiscuity,^29, 30^ *trans*-ATs,^28, 31, 32, 33^ full AT swaps,^12^ and AT active site mutagenesis.^14, 15, 16, 17, 18, 25^ Despite these advances, our lack of insight into the molecular and structural basis for substrate selectivity has meant targeting the same small number of AT residues without exploring the remainder of the ~430-residue domain or even the entire active site. Engineering selectivity between substrates often differing in only a single carbon unit without a defined binding pocket is a unique protein engineering challenge. Here, this was tackled through modeling the AT’s internal dynamics and exploration of non-traditional residues either individually or as part of structural motifs.

Guided by MD simulations, sequence analysis, and previous mutagenesis data, a total of 13 mutations (spanning 10 residues) out of less than 50 that were screened *in vitro*, resulted in significant (1.5- to >10-fold) increases in selectivity towards a provided extender. These selectivity-shifting mutations were identified throughout EryAT6’s active site, particularly in the areas proximal to the YASH loop (SSM) or the VDVVQP motif (LSM). Notably, the single mutations did not always simply increase overall substrate promiscuity but shifted selectivity towards specific extenders (e.g., L676V increased butyl incorporation at the expense of propargyl).

Notably, and consistent with previous studies, the MD simulations suggested that the previously tested combination of V742A/Y744R is unstable. This might reflect the approach used to develop these first-generation mutations, whereby mutations were identified by screening small libraries of single mutants which were then arbitrarily combined. This does not guarantee any additivity or synergy between mutations. Gratifyingly however, the substrate specificity of the AT was dramatically shifted via larger structural motif swaps that targeted the residues highlighted by MD simulations and single site mutagenesis. Harnessing the inter- and intra-motif interactions avoids the issues provoked by full domain swaps by leaving >95% of the AT unchanged.^6^ As such, both chimeras, Ery6(Tha13)TE and Ery6(Cin1)TE, retained relatively high overall activities while shifting substrate selectivity almost completely towards larger substrates, as anticipated according to the presumed native preference of each parent AT (e.g., Ery6(Tha13)TE realized a >300-fold change in pentyl (**1g**) incorporation compared to wild-type Ery6TE).

In combination with the *in vitro* assay results, the MD simulations of EryAT6 afford the most complete picture of a PKS AT domain to date. The simulations provide a description of the dynamics of the entire domain, with and without an extender unit present, and the effect(s) of mutations at various residues throughout the active site. In addition to identifying novel mutations that shift substrate specificity, new roles for other residues, such as the thioester-positioning His643 in the conserved GHSxG motif, have been proposed. Several additional insights were enabled via this comprehensive systematic approach. Examination of the mass spectra of the cyclized products produced by Ery6TE revealed the expected ions corresponding to non-natural incorporation (i.e. **4b**-**e**) but also ions corresponding to the unreduced 10-dML products (i.e. **5b**-**g**, **Fig. 4b**). This lack of KR tolerance towards the C2 extender position, only briefly reported before,^18^ will play a major role in future PKS engineering, especially with larger substrates, as the butyl and pentyl products were not reduced at all. Thus, overcoming the stringent specificity of EryAT6 has revealed a new bottleneck that needs to be overcome to capture the full catalytic capabilities of the PKS. An additional challenge that arises is the evolution of ATs capable of recognizing more structurally-diverse extenders. Neither of the inflexible and bulky extenders, thiophene- or phenylmalonyl-CoA were accepted by any Ery6(TE) variant (data not shown), but the bottleneck is unknown. As shown here, a PKS module can utilize substrates without being processed by the KR, so the failure to use the thiophene and phenyl extenders likely lies within the KS domain of Ery6. While new focus needs to be placed on engineering promiscuity of the KR and KS domains, it has been shown here that AT substrate selectivity can be modified using individual and grouped mutations throughout the AT active site, and it is anticipated that the number of implicated residues and incorporated substrates will grow as more are explored. Finally, this combined *in silico* and mutagenesis approach can be readily expanded to include other AT and substrate combinations and is expected to advance future engineering of the AT domain.

## METHODS

### General information

Materials and reagents were purchased from Sigma Aldrich unless otherwise noted. Isopropyl β-D-thiogalactoside (IPTG) was purchased from Calbiochem. The *E. coli* K207-3 strain was provided by Dr. Adrian Keatinge-Clay at the University of Texas at Austin.^34^ Construction of the pET28-Ery6TE plasmid was described previously.^14^ All module sequences are listed in **Supplementary Table S6**. Primers were purchased from Integrated DNA Technologies. All *holo* proteins were expressed in *E. coli* K207-3 cells containing *sfp*, and all *apo* proteins were expressed in *E. coli* BL21 (DE3) cells. All oligo sequences are listed in **Supplementary Table S7**.

### Homology models and molecular dynamics

Wild-type homology models for EryAT6 (with different boundaries) were created using the I-TASSER online server.^35, 36, 37^ I-TASSER was also used to create homology models of EryAT6 with motif swaps. Mutations were introduced into structurally-converged wild-type models with Discovery Studio 4.1 from Accelrys Software, Inc. Molecular graphics and analyses of MD trajectories and PDB snapshots were performed with VMD 1.9.2 and UCSF Chimera 1.10.1.^38, 39, 40^ Further analysis was performed with CPPTRAJ.^41^ Images were rendered with POV-Ray.^42^

Covalent malonyl intermediates were created using non-natural amino acids in place of the catalytic serine. The PDB files were created in ChemDraw 3D as dipeptides with the structure *N-* acetvi-X-*N*’-methylamide, where X is the non-natural serine. These dipeptides were parameterized in a similar fashion to that of the free substrates using the PyRED server and Gaussian 09.^43, 44^

Using the AMBER14 software package, individual models’ charges were neutralized with sodium ions in Xleap.^45^ All models were then solvated with a 15 Å buffer of TIP3P water, also within Xleap. The enzymes and substrates were parameterized with ff12SB and GAFF force fields from the AMBER14 software package. Prior to production MD simulations, solvated systems were treated with four heating and seven minimization steps. Steps 2, 3, 5, and 11 heated the system to 300 K over times of 20-100 ps each. The first nine steps held the protein fixed, with the restraint constant being lowered each step. Steps 10 and 11 used no restraints. Minimization steps were completed when the change in the root mean square was below 0.01 kcal mol^−1^ Å^−1^ for the first two minimization steps and below 0.001 kcal mol^−1^ Å^−1^ for the remaining minimizations. Production simulations lasted between 10 ns and 60 ns for each model. Step times were 2 fs. The non-bonded interaction cut-off was imposed at 9.0 Å.

### Expression and purification of wild-type and mutant Ery6TE

Cells were grown in 300 mL of LB media with 50 μg mL^−1^ kanamycin at 37 °C and 250 rpm until OD_600_ reached ~0.6, and then the culture was induced with 1 mM IPTG and allowed to express at 18 °C for 20 h. Proteins were purified on a nickel column with a Bio-Rad BioLogic LP FPLC (Wash buffer: 50 mM sodium phosphate, pH 7.2, 300 mM NaCl, 20 mM imidazole, and EDTA-free protease inhibitor cocktail (Roche); Elution buffer: 50 mM sodium phosphate, pH 7.2, 300 mM NaCl, 200 mM imidazole, and EDTA-free protease inhibitor cocktail), concentrated with a 10 kDa MWCO filter (EMD Millipore), and stored as single-use aliquots in module storage buffer (100 mM sodium phosphate, pH 7.4, 1 mM EDTA, 1 mM tris(2-carboxyethyl)phosphine (TCEP), 20% v/v glycerol, 0.1 μL Benzonase (NEB), and EDTA-free protease inhibitor cocktail) at −80 °C.

### MatB reactions and acyl-CoA preparation

Wild-type MatB and the mutant MatB T207G/M306I were purified and 8 mM malonyl-CoA (**1**, **2a**-**f**, **Supplementary Fig. S9**) stocks were set up as previously disclosed^46^ and described in the **Supplemental Methods**.

### Ery6TE lysate preparation

Modules (wild-type and mutant Ery6 and Ery6TE) were expressed overnight at 18 °C in 300 mL cultures in LB media with the appropriate antibiotics. Protein production was induced with 1 mM IPTG at OD_600_ of ~0.6. After overnight expression, the culture was centrifuged at 4,700 rpm for 20 min, and the supernatant was discarded. The cells were resuspended in 1 mL module storage buffer and sonicated using 51% amplitude, 10 s on, 20 s off for 10 min. After sonication, the lysed cells were centrifuged at 18,000 rpm for 1 h. The lysates were aliquoted and stored at −80 °C. Protein expression was verified by SDS-PAGE. Protein quantification was carried out using the Bradford Protein Assay Kit from Bio-Rad.

### Ery6TE pentaketide assay

The pentaketide assay was set up with a total volume of 80 μL in 100 mM sodium phosphate, pH 7.0, and 2 mM MgCl2. The reaction conditions included 1 mM TP-pentaketide (**3**), 1.75 mM of each competing malonyl-CoA from MatB reactions (**1**, **2a**-**f**), an NADPH regeneration system (5 mM glucose-6-phosphate, 500 μM NADP+, and 0.008 U/mL glucose-6-phosphate dehydrogenase), and lysate containing the module. Module concentrations were calculated by Bradford assay and SDS-PAGE gel analysis, and results were normalized based on protein content for each reaction. Negative controls were run with apo modules lacking the phosphopantetheine modifications on the ACP domains (using lysates without Sfp). Reactions proceeded at room temperature for 16 h and were quenched with an equal volume of −20 °C methanol. After quenching, all reactions were centrifuged at 10,000 *g* three times for 3 h total, and the supernatant was filtered through a nylon 0.2 μm filter. Analysis was carried out on a high-resolution mass spectrometer (ThermoFisher Scientific Exactive Plus MS, a benchtop full-scan Orbitrap™ mass spectrometer) using Heated Electrospray Ionization (HESI). The sample was analyzed via LC-MS injection into the mass spectrometer at a flow rate of 225 μL min^−1^. The mobile phase B was acetonitrile with 0.1% formic acid and mobile phase A was water with 0.1% formic acid (see **Supplementary Table S8** for gradient and scan parameters). The mass spectrometer was operated in positive ion mode. The LC column was a Thermo Hypersil Gold 50 x 2.1 mm, 1.9 μm particle size. This assay produces 10-dML and narbonolide products that can be detected as their [M+H]+, [M+H-H2O]^+^, and [M+Na]^+^ ions and keto-10-dML products that can be detected as their [M+H]^+^ and [M+Na]^+^ ions. Extracted ions for each listed ion were summed for comparison purposes. Every assay was repeated three distinct times, unless otherwise stated. For retention times, calculated masses, observed masses, representative extracted ion counts, and representative chromatograms, see **Supplementary Tables S2**, **S3**, **S5**, and **Supplementary Figures S12-S13**.

## DATA AVAILABILITY

The data that support the *in vitro* findings (e.g., enzyme assays) of this study are available within the paper or its supplementary information. The data that support the computational findings (e.g., MD simulations) are available from the corresponding author upon reasonable request.

## ACKNOWLEDGEMENTS

This study was supported in part by National Institutes of Health grants GM104258 (G.J.W.) and GM118101 (D.H.S.), and the Hans W. Vahlteich Professorship (D.H.S.). All high-resolution LC-MS measurements were made in the Molecular Education, Technology, and Research Innovation Center (METRIC) at NC State University. We thank Dr. Yaroslava G. Yingling and Dr. Hoshin Kim (NC State University) for use of their computational resources and guidance for MD simulations.

## AUTHOR CONTRIBUTIONS

E.K. and G.J.W. conceived of the project and designed the experiments. E.K. ran the MD simulations and performed the analysis. A.N.L., J.J.S. and D.H.S. synthesized and provided the TP-pentaketide substrate **6**. E.K., K.S.B., and A.M.K. ran all other experiments. E.K. and G.J.W. wrote the manuscript.

## REFERENCES

1. Weissman, K. J. Introduction to polyketide biosynthesis. In: Methods in Enzymology (ed Hopwood DA). Academic Press (2009).

2. Chen, A. Y., Schnarr, N. A., Kim, C. Y., Cane, D. E. & Khosla, C. Extender unit and acyl carrier protein specificity of ketosynthase domains of the 6-deoxyerythronolide B synthase. J. Am. Chem. Soc. 128, 3067–3074 (2006).

3. Khosla, C. Structures and mechanisms of polyketide synthases. J. Org. Chem. 74, 6416–6420 (2009).

4. Khosla, C., Tang, Y., Chen, A. Y., Schnarr, N. A. & Cane, D. E. Structure and mechanism of the 6-deoxyerythronolide B synthase. Annu. Rev. Biochem. 76, 195–221 (2007).

5. Chan, Y. A., Podevels, A. M., Kevany, B. M. & Thomas, M. G. Biosynthesis of polyketide synthase extender units. Nat. Prod. Rep. 26, 90–114 (2009).

6. Kalkreuter, E. & Williams, G. J. Engineering enzymatic assembly lines for the production of new antimicrobials. Curr. Opin. Microbiol. 45, 140–148 (2018).

7. Hans, M., Hornung, A., Dziarnowski, A., Cane, D. E. & Khosla, C. Mechanistic analysis of acyl transferase domain exchange in polyketide synthase modules. J. Am. Chem. Soc. 125, 5366–5374 (2003).

8. Del Vecchio, F., et al. Active-site residue, domain and module swaps in modular polyketide synthases. J. Ind. Microbiol. Biotechnol. 30, 489–494 (2003).

9. Chemler, J. A., et al. Evolution of efficient modular polyketide synthases by homologous recombination. J. Am. Chem. Soc. 137, 10603–10609 (2015).

10. Shen, J. J., et al. Substrate specificity of acyltransferase domains for efficient transfer of acyl groups. Front. Microbiol. 9, 1840 (2018).

11. Khayatt, B. I., Overmars, L., Siezen, R. J. & Francke, C. Classification of the adenylation and acyl-transferase activity of NRPS and PKS systems using ensembles of substrate specific hidden Markov models. PLoS One 8, e62136 (2013).

12. Yuzawa, S., et al. Comprehensive in vitro analysis of acyltransferase domain exchanges in modular polyketide synthases and its application for short-chain ketone production. ACS Synth. Biol. 6, 139–147 (2017).

13. Petkovic, H., et al. Substrate specificity of the acyl transferase domains of EpoC from the epothilone polyketide synthase. Org. Biomol. Chem. 6, 500–506 (2008).

14. Koryakina, I., et al. Inversion of extender unit selectivity in the erythromycin polyketide synthase by acyltransferase domain engineering. ACS Chem. Biol. 12, 114–123 (2017).

15. Bravo-Rodriguez, K., et al. Substrate flexibility of a mutated acyltransferase domain and implications for polyketide biosynthesis. Chem. Biol. 22, 1425–1430 (2015).

16. Sundermann, U., et al. Enzyme-directed mutasynthesis: a combined experimental and theoretical approach to substrate recognition of a polyketide synthase. ACS Chem. Biol. 8, 443–450 (2013).

17. Li, Y., et al. Structural basis of a broadly selective acyltransferase from the polyketide synthase of splenocin. Angew. Chem. Int. Ed. 57, 5823–5827 (2018).

18. Kalkreuter, E., CroweTipton, J. M., Lowell, A. N., Sherman, D. H. & Williams, G. J. Engineering the substrate specificity of a modular polyketide synthase for installation of consecutive non-natural extender units. J. Am. Chem. Soc. 141, 1961–1969 (2019).

19. Bravo-Rodriguez, K., et al. Predicted incorporation of non-native substrates by a polyketide synthase yields bioactive natural product derivatives. ChemBioChem 15, 1991–1997 (2014).

20. Reeves, C. D., Sumati, M., Ashley, G. W., Piagentini, M., Hutchinson, C. R. & McDaniel, R. Alteration of the substrate specificity of a modular polyketide synthase acyltransferase domain through site-specific mutations. Biochemistry 40, 15464–15470 (2001).

21. Tang, Y., Kim, C. Y., Mathews, II, Cane, D. E. & Khosla, C. The 2.7-Angstrom crystal structure of a 194-kDa homodimeric fragment of the 6-deoxyerythronolide B synthase. Proc. Natl. Acad. Sci. U. S. A. 103, 11124–11129 (2006).

22. Tang, Y., Chen, A. Y., Kim, C.-Y., Cane, D. E. & Khosla, C. Structural and mechanistic analysis of protein interactions in module 3 of the 6-deoxyerythronolide B synthase. Chem. Biol. 14, 931–943 (2007).

23. Poust, S., Yoon, I., Adams, P. D., Katz, L., Petzold, C. J. & Keasling, J. D. Understanding the role of histidine in the GHSxG acyltransferase active site motif: evidence for histidine stabilization of the malonyl-enzyme intermediate. PLoS One 9, e109421 (2014).

24. Mortison, J. D., Kittendorf, J. D. & Sherman, D. H. Synthesis and biochemical analysis of complex chain-elongation intermediates for interrogation of molecular specificity in the erythromycin and pikromycin polyketide synthases. J. Am. Chem. Soc. 131, 15784–15793 (2009).

25. Vögeli, B., Geyer, K., Gerlinger, P. D., Benkstein, S., Cortina, N. S. & Erb, T. J. Combining promiscuous acyl-CoA oxidase and enoyl-CoA carboxylase/reductases for atypical polyketide extender unit biosynthesis. Cell Chem. Biol. 25, 833–839.e834 (2018).

26. Rachid, S., et al. Mining the cinnabaramide biosynthetic pathway to generate novel proteasome inhibitors. ChemBioChem 12, 922–931 (2011).

27. Greule, A., Intra, B., Flemming, S., Rommel, M. G., Panbangred, W. & Bechthold, A. The draft genome sequence of *Actinokineospora bangkokensis* 44EHW(T) reveals the biosynthetic pathway of the antifungal thailandin compounds with unusual butylmalonyl-CoA extender units. Molecules 21, (2016).

28. Walker, M. C., Thuronyi, B. W., Charkoudian, L. K., Lowry, B., Khosla, C. & Chang, M. C. Y. Expanding the fluorine chemistry of living systems using engineered polyketide synthase pathways. Science 341, 1089–1094 (2013).

29. Moller, D., et al. Flexible enzymatic activation of artificial polyketide extender units by *Streptomyces cinnamonensis* into the monensin biosynthetic pathway. Lett. Appl. Microbiol. 67, 226–234 (2018).

30. Koryakina, I., McArthur, J. B., Draelos, M. M. & Williams, G. J. Promiscuity of a modular polyketide synthase towards natural and non-natural extender units. Org. Biomol. Chem. 11, 4449–4458 (2013).

31. Thuronyi, B. W., Privalsky, T. M. & Chang, M. C. Y. Engineered fluorine metabolism and fluoropolymer production in living cells. Angew. Chem. Int. Ed. 56, 13637–13640 (2017).

32. Musiol-Kroll, E. M., et al. Polyketide bioderivatization using the promiscuous acyltransferase KirCII. ACS Synth. Biol. 6, 421–427 (2017).

33. Carpenter, S. M. & Williams, G. J. Extender unit promiscuity and orthogonal protein interactions of an aminomalonyl-ACP utilizing trans-acyltransferase from zwittermicin biosynthesis. ACS Chem. Biol. 13, 3361–3373 (2018).

34. Murli, S., Kennedy, J., Dayem, L. C., Carney, J. R. & Kealey, J. T. Metabolic engineering of *Escherichia coli* for improved 6-deoxyerythronolide B production. J. Ind. Microbiol. Biotechnol. 30, 500–509 (2003).

35. Yang, J., Yan, R., Roy, A., Xu, D., Poisson, J. & Zhang, Y. The I-TASSER Suite: protein structure and function prediction. Nat. Methods 12, 7–8 (2015).

36. Roy, A., Kucukural, A. & Zhang, Y. I-TASSER: a unified platform for automated protein structure and function prediction. Nat. Protoc. 5, 725–738 (2010).

37. Zhang, Y. I-TASSER server for protein 3D structure prediction. BMC Bioinform. 9, 40 (2008).

38. W. Humphrey, A. D., and K. Schulten. VMD - Visual Molecular Dynamics. J. Mol. Graph. 14, 33–38 (1996).

39. Pettersen, E. F., et al. UCSF Chimera--a visualization system for exploratory research and analysis. J. Comput. Chem. 25, 1605–1612 (2004).

40. Sanner, M. F., Olson, A. J. & Spehner, J. C. Reduced surface: an efficient way to compute molecular surfaces. Biopolymers 38, 305–320 (1996).

41. Roe, D. R. & Cheatham, T. E., 3rd. PTRAJ and CPPTRAJ: Software for processing and analysis of molecular dynamics trajectory data. J. Chem. Theory Comput. 9, 3084–3095 (2013).

42. Persistence of Vision Pty Ltd. Persistence of Vision Raytracer (Version 3.6).) (2004).

43. E. Vanquelef, S. S., G. Marquant, E. Garcia, G. Klimerak, J.C. Delepine, P. Cieplak, and F.Y. Dupradeau. R.E.D. Server: a web service for deriving RESP and ESP charges and building force field libraries for new molecules and molecular fragments. Nucl. Acids Res. 39, W511 (2011).

44. Frisch, M. J., et al. Gaussian 09. (Gaussian, Inc., 2009).

45. Case, D. A., et al. AMBER 2015. (University of California, San Francisco, 2015).

46. Koryakina, I., et al. Poly specific trans-acyltransferase machinery revealed via engineered acyl-CoA synthetases. ACS Chem. Biol. 8, 200–208 (2013).

