## Supplemental Information for "Computationally-guided exchange of substrate selectivity motifs in a modular polyketide synthase acyltransferase"

**Table of contents**

***Supplementary Tables***

**Supplementary Table S1. Wild-type EryAT6 amino acid sequences for MD simulations.**

**Supplementary Table S2. High-Resolution LC-MS retention times, calculated masses, and observed masses for DEBS PKS-catalyzed reaction products.**

**Supplementary Table S3. High-resolution LC-MS peak areas for PKS-catalyzed reaction products with five extenders.**

**Supplementary Table S4. Mutations in Ery6TE that did not result in a shift from the wild-type product distribution.**

**Supplementary Table S5. High-resolution LC-MS peak areas for PKS-catalyzed reaction products with two extenders.**

**Supplementary Table S6. DEBS PKS DNA FASTA sequences.**

**Supplementary Table S7. Primers for DEBS PKS construction and mutagenesis.**

**Supplementary Table S8. High-resolution LC-MS parameters and gradient.**

***Supplementary Figures***

**Supplementary Figure S1. Conserved motifs of PKS acyltransferases.**

**Supplementary Figure S2. Evaluation of EryAT6 models.**
**Supplementary Figure S3. Structural convergence of EryAT6 MD simulations.**
**Supplementary Figure S4. Snapshots of EryAT6 individual mutations.**
**Supplementary Figure S5. RMSF plot for EryAT6 V742A/Y744R simulation.**
**Supplementary Figure S6. EryAT6 wild-type with methylmalonyl-CoA and propargylmalonyl-CoA.**
**Supplementary Figure S7. EryAT6 wild-type with propargylmalonated Ser644.**
**Supplementary Figure S8. Retention of Asp743 salt bridge.**
**Supplementary Figure S9. Representative HPLC traces for MatB-synthesized malonyl-CoAs.**
**Supplementary Figure S10. Motif sequences from natural acyltransferases.**
**Supplementary Figure S11. Surface models of motif-swapped EryAT6.**
**Supplementary Figure S12. Representative mass spectra of 10-deoxymethynolide B compounds from lysate module reactions.**
**Supplementary Figure S13. Representative mass spectra of keto-10-deoxymethynolide B compounds from lysate module reactions.**

***Supplemental Methods***
**Site-directed mutagenesis of Ery6TE**
**Construction of Ery6TE motif chimeras**
**Expression and purification of wild-type and mutant MatB**
**Synthesis of acyl-CoAs by MatB**

***Supplemental References***

**Supplementary Tables**

**Table S1. Wild-type EryAT6 amino acid sequences for MD simulations.** Linkers are in bold.

| EryAT6 Construct | Amino Acid FASTA Sequences for MD Simulations |
|---|---|
| **Wild-Type with Both Linkers** | **AEPPEPEPLPEPGPVGVLAAANSVPVLLSARTETALAAQARLLESAVDDSVPLTA**<br>**LASALATGRAHLPRRAALLAGDHEQLRGQLRAVAEGVAAPGATTGTASAGGVV**<br>**F**VFPGQGAQWEGMARGLLSVPVFAESIAECDAVLSEVAGFSASEVLEQRPDAPSL<br>ERVDVVQPVLFSVMVSLARLWGACGVSPSAVIGHSQGEIAAAVVAGVLSLEDGVR<br>VVALRAKALRALAGKGGMVSLAAPGERARALIAPWEDRISVAAVNSPSSVVVSGD<br>PEALAELVARCEDEGVRAKTLPVDYASHSRHVEEIRETILADLDGISARRAAIPLYST<br>LHGERRDGADMGPRYWYDNLRSQVRFDEAVSAAVADGHATFVEMSPHPVLTAA<br>VQE**IAADAVAIGSLHRDTAEEHLIAELARAHVHGVAVDWRNVFPAA** |
| **Wild-Type with N-Terminal Linker** | **AEPPEPEPLPEPGPVGVLAAANSVPVLLSARTETALAAQARLLESAVDDSVPLTA**<br>**LASALATGRAHLPRRAALLAGDHEQLRGQLRAVAEGVAAPGATTGTASAGGVV**<br>**F**VFPGQGAQWEGMARGLLSVPVFAESIAECDAVLSEVAGFSASEVLEQRPDAPSL<br>ERVDVVQPVLFSVMVSLARLWGACGVSPSAVIGHSQGEIAAAVVAGVLSLEDGVR<br>VVALRAKALRALAGKGGMVSLAAPGERARALIAPWEDRISVAAVNSPSSVVVSGD<br>PEALAELVARCEDEGVRAKTLPVDYASHSRHVEEIRETILADLDGISARRAAIPLYST<br>LHGERRDGADMGPRYWYDNLRSQVRFDEAVSAAVADGHATFVEMSPHPVLTAA<br>VQE |
| **Wild-Type with No Linkers** | VFPGQGAQWEGMARGLLSVPVFAESIAECDAVLSEVAGFSASEVLEQRPDAPSLE<br>RVDVVQPVLFSVMVSLARLWGACGVSPSAVIGHSQGEIAAAVVAGVLSLEDGVRV<br>VALRAKALRALAGKGGMVSLAAPGERARALIAPWEDRISVAAVNSPSSVVVSGDP<br>EALAELVARCEDEGVRAKTLPVDYASHSRHVEEIRETILADLDGISARRAAIPLYSTL<br>HGERRDGADMGPRYWYDNLRSQVRFDEAVSAAVADGHATFVEMSPHPVLTAAV<br>QE |

**Table S2. High-Resolution LC-MS retention times, calculated masses, and observed masses for DEBS PKS-catalyzed reaction products.**
N.D. = Not Detected.

| Compound & Retention Time (min) | | Calculated Mass ($[M-H_2O+H]^+$) | Observed Mass ($[M-H_2O+H]^+$) | Calculated Mass ($[M+H]^+$) | Observed Mass ($[M+H]^+$) | Calculated Mass ($[M+Na]^+$) | Observed Mass ($[M+Na]^+$) |
|---|---|---|---|---|---|---|---|
| **4** | 8.03 | 279.1955 | 279.1962 | 297.2060 | 297.2069 | 319.1880 | 319.1904 |
| **5a** | 9.68 | 293.2111 | 293.2119 | 311.2217 | 311.2227 | 333.2036 | 333.2044 |
| **5b** | 8.96 | 303.1955 | 303.1963 | 321.2060 | 321.2068 | 343.1880 | 343.1890 |
| **5c** | 10.63 | 307.2268 | 307.2274 | 325.2373 | 325.2378 | 347.2193 | 347.2198 |
| **5d** | N.D. | 321.2424 | N.D. | 339.2530 | N.D. | 361.2349 | N.D. |
| **5e** | 10.12 | 305.2111 | 305.2105 | 323.2217 | 323.2201 | 345.2036 | 345.2035 |
| **5f** | N.D. | 335.2581 | N.D. | 353.2686 | N.D. | 375.2506 | N.D. |
| **6a** | 12.39 | -- | -- | 309.2060 | 309.2067 | 331.1880 | 331.1886 |
| **6b** | 11.80 | -- | -- | 319.1904 | 319.1911 | 341.1723 | 341.1732 |
| **6c** | 11.79 | -- | -- | 323.2217 | 323.2224 | 345.2036 | 345.2044 |
| **6d** | 14.56 | -- | -- | 337.2373 | 337.2380 | 359.2193 | 359.2200 |
| **6e** | 12.67 | -- | -- | 321.2060 | 321.2055 | 343.1880 | 343.1876 |
| **6f** | 15.07 | -- | -- | 351.2530 | 351.2525 | 373.2349 | 373.2345 |

**Table S3. High-resolution LC-MS peak areas for PKS-catalyzed reaction products with five extenders.** Extracted ion count peak areas from one replicate are shown for each reaction condition, prior to correction with enzyme concentration. N.D. = Not Detected.

| **Ery6TE Variant** | **Representative EIC Peak Areas** | | | | |
|---|---|---|---|---|---|
| | **4** | **5a** | **6a** | **5b** | **6b** |
| WT | 211,060,257 | 8,350,707 | 3,201,023 | 60,428,975 | 33,336,885 |
| V742A | 1,074,530,536 | 74,181,724 | 8,289,132 | 621,019,227 | 62,604,070 |
| Y744R | 201,560,426 | 18,273,398 | 10,450,848 | 425,370,308 | 221,736,187 |
| L673G | 412,033,607 | 11,485,854 | 3,946,441 | 96,211,067 | 47,618,973 |
| A672S | 918,117,379 | 17,567,468 | 16,886,399 | 174,246,461 | 190,537,319 |
| L676V | 913,134,214 | 20,437,965 | 21,610,476 | 202,357,895 | 254,127,057 |
| G681A | 258,612,993 | 7,211,911 | 4,379,076 | 42,211,949 | 38,450,516 |
| D613E | 1,112,681,260 | 25,901,052 | 23,610,928 | 246,571,818 | 263,851,451 |
| R674A | 1,137,937,000 | 22,945,009 | 25,007,975 | 178,486,582 | 202,512,483 |
| H643S | 244,645,247 | 12,743,255 | 8,023,759 | 70,681,023 | 47,042,985 |
| H643F | 215,279,432 | 1,629,754 | 960,271 | 2,131,336 | 2,185,209 |
| I648G | 365,230,755 | 869,632 | 475,403 | 1,770,386 | 1,262,385 |
| L797E | 565,835,459 | 782,613 | 2,352,056 | 829,163 | 1,769,043 |
| Cin1 | 1,122,551 | 14,392 | 29,947 | 1,093,275 | 2,750,716 |
| Tha13 | 29,754,912 | 1,565,605 | 928,229 | 31,038,395 | 17,687,295 |

| **Ery6TE Variant** | **Representative EIC Peak Areas** | | | |
|---|---|---|---|---|
| | **5c** | **6c** | **5d** | **6d** |
| WT | 11,975,213 | 22,923,302 | N.D. | 3,401,711 |
| V742A | 41,645,025 | 4,688,419 | N.D. | 32,702,810 |
| Y744R | 87,834,327 | 60,330,553 | N.D. | 110,340,604 |
| L673G | 12,692,412 | 27,903,903 | N.D. | 15,313,652 |
| A672S | 7,799,155 | 195,172,467 | N.D. | 18,325,167 |
| L676V | 2,065,221 | 161,120,479 | N.D. | 28,957,619 |
| G681A | 13,320,019 | 63,892,775 | N.D. | 6,301,166 |
| D613E | 20,124,173 | 244,569,998 | N.D. | 37,219,146 |
| R674A | 16,724,087 | 256,290,037 | N.D. | 19,347,354 |
| H643S | 25,063,933 | 35,489,999 | N.D. | 18,784,167 |
| H643F | 74,197 | 3,727,611 | N.D. | 589,128 |
| I648G | 595,456 | 6,006,129 | N.D. | 1,376,049 |
| L797E | 128,196 | 1,623,863 | N.D. | 1,573,907 |
| Cin1 | 39,917 | 439,077 | N.D. | 1,402,533 |
| Tha13 | 22,492,383 | 39,231,371 | N.D. | 123,779,145 |

**Table S4. Mutations in Ery6TE that did not result in a shift from the wild-type product distribution.** All mutants that were constructed and tested in vitro with **1** and non-native extenders but showed no difference in product distribution (other than total activity) from wild-type Ery6TE were excluded from the main text but are included in the following table.

| **Ery6TE Mutants Indistinguishable from Wild-Type** | | |
|---|---|---|
| A596L | A672S | L740V |
| E598D | R674A | P741Δ |
| V612D | A675L | P741G |
| E647D | G689A | P790A |
| G662A | D701G | L797S |
| A670T | A707G | |
| A670V | S717A | |

**Table S5. High-resolution LC-MS peak areas for PKS-catalyzed reaction products with two extenders.** Extracted ion count peak areas from one replicate are shown for each reaction condition, prior to correction with enzyme concentration. N.D. = Not Detected.

| | | **Representative EIC Peak Areas** | | | | |
|---|---|---|---|---|---|---|
| Substrates | Product | WT | I648G | Y744R | Cin1 | Tha13 |
| **1 + 2b** | **4** | 532,578,041 | 799,908,718 | 78,684,251 | 29,848,810 | 26,068,751 |
| | **5b** | 31,090,929 | 2,583,102 | 320,918,411 | 20,904,574 | 45,737,083 |
| | **6b** | 75,186,842 | N.D. | 109,088,198 | 44,368,738 | 37,497,560 |
| **1 + 2d** | **4** | 1,241,364,749 | 650,111,454 | 230,399,192 | 478,245 | 29,616,734 |
| | **5d** | N.D. | N.D. | N.D. | N.D. | N.D. |
| | **6d** | 6,953,819 | N.D. | 88,178,801 | 4,152,337 | 193,602,896 |
| **1 + 2e** | **4** | 1,263,268,254 | 1,467,946,423 | 122,464,111 | 45,392,299 | 43,610,053 |
| | **5e** | 7,647,855 | N.D. | 28,872,748 | 964,634 | 10,197,042 |
| | **6e** | 21,784,302 | N.D. | 53,689,472 | 17,802,789 | 33,024,254 |
| **1 + 1f** | **4** | 1,144,743,869 | 1,115,568,669 | 622,633,725 | 1,042,802,675 | 30,377,170 |
| | **5f** | N.D. | N.D. | N.D. | N.D. | N.D. |
| | **6f** | 1,583,892 | N.D. | 217,518,820 | 4,458,627,491 | 473,073,518 |

**Table S6. DEBS PKS DNA FASTA sequences.** AT domains in modules are in bold. Pik docking domain is italicized.

| Construct | Nucleotide Sequences |
|---|---|
| **ThaAT13 Motifs** | GACCCAATACGCGCCGTCGCTGGAGCGTCTGGATGTGAATCAGCCGGTTCTGTTTAGT<br>GTTATGGTGAGTCTGGCCCGTCTGTGGGGTGCCTGTGGTGTGAGTCCGAGTGCCGTTA<br>TTGGCCATAGCCAGGGTGAAATTGCAGCAGCAGTTGTTGCAGGCGTGCTGAGTCTGGA<br>AGATGGTGTGCGCGTGGTTGCACTGCGTGCCAAAGCACTGCGTGCGCTGGCAGGCAAA<br>GGTGGCATGGTTAGCCTGGCCGCACCGGGCGAACGCGCTAGAGCTCTGATTGCCCCGT<br>GGGAAGATCGTATTAGCGTTGCCGCAGTTAATAGTCCGAGTAGCGTGGTGGTGAGTGG<br>TGACCCGGAAGCCCTGGCCGAACTGGTGGCACGTTGTGAAGATGAAGGTGTTCGCGCA<br>AAAGTTCTGCCGGGTGCCGATGCCGCAGGCCACTCCCGCCACGTCGAGACCCAAT |
| **CinAT1 Motifs** | ACGCGCCGTCGCTGGAGGAAGGCGATATTCAGCAGCCGGTTCTGTTTAGTGTGATGGT<br>TAGTCTGGCACGCCTGTGGGGCGCCTGCGGTGTTAGTCCGAGTGCAGTTATTGGCCAT<br>AGCCAGGGCGAAATTGCAGCCGCCGTTGTTGCAGGCGTTCTGAGCCTGGAAGATGGCG<br>TTCGTGTTGTGGCCCTGCGCGCCAAAGCACTGCGCGCACTGGCAGGCAAAGGTGGTAT<br>GGTGAGTCTGGCAGCACCGGGTGAACGCGCCCGTGCTCTGATTGCACCGTGGGAAGAT<br>CGCATTAGCGTGGCCGCAGTGAATAGCCCGAGCAGCGTTGTTGTTAGTGGCGATCCGG<br>AAGCCCTGGCAGAACTGGTGGCCCGTTGCGAAGATGAAGGCGTTCGTGCCAAAGCCCT<br>GCGCGTGGAACGCGCAGGCCACTCCCGCCACGTCGA |
| **Ery6 WT** | ATGGTCGGCGCAGCAGAGGCGGAGCAAGCCCCGGCGCTCGTGCGCGAGGTGCCGAAGG<br>ATGCCGACGACCCGATCGCGATCGTCGGCATGGCCTGCCGCTTCCCCGGCGGCGTGCA<br>CAACCCCGGTGAGCTGTGGGAGTTCATCGTCGGCGGCGGAGACGCCGTGACGGAGATG<br>CCCACCGACCGCGGCTGGGACCTCGACGCGCTGTTCGACCCCGACCCGCAGCGCCACG<br>GAACCAGCTACTCGCGACACGGCGCGTTCCTCGACGGGGCCGCCGACTTCGACGCGGC<br>GTTCTTCGGGATCTCGCCGCGCGAGGCGCTGGCGATGGACCCGCAGCAGCGCCAGGTC<br>CTGGAAACGACGTGGGAGCTGTTCGAGAACGCCGGCATCGACCCGCACTCGCTGCGGG<br>GCAGCGACACCGGCGTCTTCCTCGGCGCCGCGTACCAGGGCTACGGCCAGGACGCGGT<br>GGTGCCCGAGGACAGCGAGGGCTACCTGCTCACCGGCAACTCCTCCGCCGTGGTGTCC<br>GGCCGGGTCGCCTACGTGCTGGGGCTGGAAGGCCCCGCGGTCACGGTGGACACGGCGT<br>GTTCGTCGTCGTTGGTGGCCTTGCATTCGGCGTGTGGGTCGTTGCGTGACGGTGACTG<br>CGGTCTTGCGGTGGCCGGTGGTGTGTCGGTGATGGCGGGCCCGGAGGTGTTCACCGAG<br>TTCTCCCGCCAGGGCGGCTTGGCCGTGGACGGGCGCTGCAAGGCGTTCTCCGCGGAGG<br>CCGACGGCTTCGGTTTCGCCGAGGGCGTCGCGGTGGTCCTGCTCCAGCGGTTGTCCGA<br>CGCCCGCAGGGCGGGTCGCCAGGTGCTCGGCGTGGTCGCGGGCTCGGCGATCAACCAG<br>GACGGCGCGAGCAACGGTCTCGCGGCGCCGAGCGGCGTCGCCCAGCAGCGCGTGATCC<br>GCAAGGCGTGGGCGCGTGCGGGGATCACGGGCGCGGATGTGGCCGTGGTGGAGGCGCA<br>TGGGACCGGTACGCGGCTGGGCGATCCGGTGGAGGCGTCGGCGTTGCTGGCTACTTAC<br>GGCAAGTCGCGCGGGTCGTCGGGCCCGGTGCTGCTGGGTTCGGTGAAGTCGAACATCG<br>GTCACGCGCAGGCGGCCGCGGGTGTCGCGGGCGTGATCAAGGTGGTCCTGGGGTTGAA<br>CCGCGGCCTGGTGCCGCCGATGCTCTGCCGCGGCGAGCGGTCGCCGCTGATCGAATGG<br>TCCTCGGGTGGTGTGGAACTTGCCGAGGCCGTGAGCCCGTGGCCTCCGGCCGCGGACG<br>GGGTGCGCCGGGCCGGTGTGTCGGCGTTCGGGGTGAGC**GGGACGAACGCGCACGTGAT**<br>**CATCGCCGAGCCCCCGGAGCCCGAGCCGCTGCCGGAACCCGGACCGGTGGGCGTGCTG**<br>**GCCGCTGCGAACTCGGTGCCCGTACTGCTGTCGGCCAGGACCGAGACCGCGTTGGCAG**<br>**CGCAGGCGCGGCTCCTGGAGTCCGCAGTGGACGACTCGGTTCCGTTGACGGCATTGGC**<br>**TTCCGCGCTGGCCACCGGACGCGCCCACCTGCCGCGTCGTGCGGCGTTGCTGGCAGGC**<br>**GACCACGAACAGCTCCGCGGGCAGTTGCGAGCGGTCGCCGAGGGCGTTGCGGCTCCCG**<br>**GTGCCACCACCGGAACCGCCTCCGCCGGCGGCGTGGTTTTCGTCTTCCCAGGTCAGGG**<br>**TGCTCAGTGGGAGGGCATGGCCCGGGGCTTGCTCTCGGTCCCCGTCTTCGCCGAGTCG**<br>**ATCGCCGAGTGCGATGCGGTGTTGTCGGAGGTGGCCGGGTTCTCGGCCTCCGAAGTGC**<br>**TGGAGCAGCGTCCGGACGCGCCGTCGCTGGAGCGGGTCGACGTCGTACAGCCGGTGTT** |

| | |
|---|---|
| | **GTTCTCCGTGATGGTGTCGCTGGCGCGGCTGTGGGCGCTTGCGGAGTCAGCCCCTCG<br>GCCGTCATCGGCCATTCGCAGGGCGAGATCGCCGCCGCGGTGGTGGCCGGGGTGTTGT<br>CGCTGGAGGACGGCGTGCGCGTCGTGGCCCTGCGCGCGAAGGCGTTGCGTGCGCTGGC<br>GGGCAAGGGCGGCATGGTCTCGTTGGCGGCTCCCGGTGAACGCGCCCGCGCGCTGATC<br>GCACCGTGGGAGGACCGGATCTCCGTCGCGGCGGTCAACTCCCCGTCCTCGGTCGTGG<br>TCTCCGGCGATCCGGAGGCGCTGGCCGAACTCGTCGCACGTTGCGAGGACGAGGGCGT<br>GCGCGCCAAGACGCTCCCGGTGGACTACGCCTCGCACTCCCGCCACGTCGAGGAGATC<br>CGCGAGACGATCCTCGCCGACCTCGACGGCATCTCCGCGCGGCGTGCCGCCATCCCGC<br>TCTACTCCACGCTGCACGGCGAACGGCGCGACGGCGCCGACATGGGTCCGCGGTACTG<br>GTACGACAACCTGCGCTCCCAGGTGCGCTTCGACGAGGCGGTCTCGGCCGCCGTCGCC<br>GACGGTCACGCCACCTTCGTCGAGATGAGCCCGCACCCGGTGCTCACCGCGGCGGTGC<br>AGGAGATCGCCGCGGACGCCGTGGCCATCGGGTCGCTGCACCGCGACACCGCGGAGGA<br>GCACCTGATCGCCGAGCTCGCCCGGGCGCACGTGCACGGCGTGGCCGTGGACTGGCGG<br>AACGTCTTCCCGGCGGCACCTCCGGTGGCG**CTGCCCAACTACCCGTTCGAGCCCCAGC<br>GGTACTGGCTCGCGCCGGAGGTGTCCGACCAGCTCGCCGACAGCCGCTACCGCGTCGA<br>CTGGCGACCGCTGGCCACCACGCCGGTGGACCTGGAAGGCGGCTTCCTGGTCCACGGG<br>TCCGCACCGGAGTCGCTGACCAGCGCAGTCGAGAAGGCCGGAGGCCGCGTCGTGCCGG<br>TCGCCTCGGCCGACCGCGAAGCGCTCGCGGCGGCCCTGCGGGAGGTGCCGGGCGAGGT<br>CGCCGGCGTGCTCTCGGTCCACACCGGCGCCGCAACGCACCTCGCCCTGCACCAGTCG<br>CTGGGTGAGGCCGGCGTGCGGGCCCCGCTCTGGCTGGTCACCAGCCGAGCGGTCGCGC<br>TCGGGGAGTCCGAGCCGGTCGATCCCGAGCAGGCGATGGTGTGGGGTCTCGGGCGCGT<br>CATGGGCCTGGAGACCCCGGAACGGTGGGGCGGTCTGGTGGACCTGCCCGCCGAACCC<br>GCGCCGGGGGACGGCGAGGCGTTCGTCGCCTGCCTCGGCGCGGACGGCCACGAGGACC<br>AGGTCGCGATCCGTGACCACGCCCGCTACGGCCGCCGCCTCGTCCGCGCCCCGCTGGG<br>CACCCGCGAGTCGAGCTGGGAGCCGGCGGGCACGGCGCTGGTCACCGGCGGCACCGGT<br>GCGCTCGGCGGCCACGTCGCCCGCCACCTCGCCAGGTGCGGGGTGGAGGACCTGGTGC<br>TGGTCAGCAGGCGCGGCGTCGACGCTCCCGGCGCGGCCGAGCTGGAAGCCGAACTGGT<br>CGCCCTCGGCGCGAAGACGACCATCACCGCCTGCGACGTGGCCGACCGCGAGCAGCTC<br>TCCAAGCTGCTGGAAGAACTGCGCGGGCAGGGACGTCCGGTGCGGACCGTCGTGCACA<br>CCGCCGGGGTGCCCGAATCGAGGCCGCTGCACGAGATCGGCGAGCTGGAGTCGGTCTG<br>CGCGGCGAAGGTGACCGGGGCCCGGCTGCTCGACGAGCTGTGCCCGGACGCCGAGACC<br>TTCGTCCTGTTCTCGTCCGGAGCGGGGGTGTGGGGCAGTGCGAACCTCGGCGCCTACT<br>CCGCGGCCAACGCCTACCTCGACGCGCTGGCCCACCGCCGCCGTGCGGAAGGCCGTGC<br>GGCGACGTCCGTCGCGTGGGGCGCCTGGGCGGGCGAGGGCATGGCCACCGGCGACCTC<br>GAGGGGCTCACCCGGCGCGGCCTGCGCCCGATGGCGCCCGAGCGCGCGATCCGCGCGC<br>TGCACCAGGCGCTGGACAACGGCGACACGTGCGTTTCGATCGCCGACGTCGACTGGGA<br>GCGCTTCGCGGTCGGCTTCACCGCCGCCCGGCCGCGTCCGCTGCTGGACGAGCTCGTC<br>ACGCCGGCGGTGGGGGCCGTCCCCGCGGTGCAGGCGGCCCCGGCGCGGGAGATGACGT<br>CGCAGGAGTTGCTGGAGTTCACGCACTCGCACGTCGCGGCGATCCTCGGGCATTCCAG<br>CCCGGACGCGGTCGGGCAGGACCAGCCGTTCACCGAGCTCGGCTTCGACTCGCTGACC<br>GCGGTCGGGCTGCGCAACCAGCTCCAGCAGGCCACCGGGCTCGCGCTGCCCGCGACCC<br>TGGTGTTCGAGCACCCCACGGTCCGCAGGTTGGCCGACCACATAGGACAGCAGCTCTG<br>A |
| **PDDEry6TE WT** | *ATGACGAGTTCCAACGAACAGTTGGTGGACGCTCTGCGCGCCTCTCTCAAGGAGAACG<br>AAGAACTCCGGAAAGAGAGCCGTCGCCGGGCCGACCGTCGGCAGGAGGAG*ATCGCGAT<br>CGTCGGCATGGCCTGCCGCTTCCCCGGCGGCGTGCACAACCCCGGTGAGCTGTGGGAG<br>TTCATCGTCGGCGGCGGAGACGCCGTGACGGAGATGCCCACCGACCGCGGCTGGGACC<br>TCGACGCGCTGTTCGACCCCGACCCGCAGCGCCACGGAACCAGCTACTCGCGACACGG<br>CGCGTTCCTCGACGGGGCCGCCGACTTCGACGCGGCGTTCTTCGGGATCTCGCCGCGC<br>GAGGCGCTGGCGATGGACCCGCAGCAGCGCCAGGTCCTGGAAACGACGTGGGAGCTGT<br>TCGAGAACGCCGGCATCGACCCGCACTCGCTGCGGGGCAGCGACACCGGCGTCTTCCT<br>CGGCGCCGCGTACCAGGGCTACGGCCAGGACGCGGTGGTGCCCGAGGACAGCGAGGGC |

| | |
|---|---|
| | TACCTGCTCACCGGCAACTCCTCCGCCGTGGTGTCCGGCCGGGTCGCCTACGTGCTGG<br>GGCTGGAAGGCCCCGCGGTCACGGTGGACACGGCGTGTTCGTCGTCGTTGGTGGCCTT<br>GCATTCGGCGTGTGGGTCGTTGCGTGACGGTGACTGCGGTCTTGCGGTGGCCGGTGGT<br>GTGTCGGTGATGGCGGGCCCGGAGGTGTTCACCGAGTTCTCCCGCCAGGGCGGCTTGG<br>CCGTGGACGGGCGCTGCAAGGCGTTCTCCGCGGAGGCCGACGGCTTCGGTTTCGCCGA<br>GGGCGTCGCGGTGGTCCTGCTCCAGCGGTTGTCCGACGCCCGCAGGGCGGGTCGCCAG<br>GTGCTCGGCGTGGTCGCGGGCTCGGCGATCAACCAGGACGGCGCGAGCAACGGTCTCG<br>CGGCGCCGAGCGGCGTCGCCCAGCAGCGCGTGATCCGCAAGGCGTGGGCGCGTGCGGG<br>GATCACGGGCGCGGATGTGGCCGTGGTGGAGGCGCATGGGACCGGTACGCGGCTGGGC<br>GATCCGGTGGAGGCGTCGGCGTTGCTGGCTACTTACGGCAAGTCGCGCGGGTCGTCGG<br>GCCCGGTGCTGCTGGGTTCGGTGAAGTCGAACATCGGTCACGCGCAGGCGGCCGCGGG<br>TGTCGCGGGCGTGATCAAGGTGGTCCTGGGGTTGAACCGCGGCCTGGTGCCGCCGATG<br>CTCTGCCGCGGCGAGCGGTCGCCGCTGATCGAATGGTCCTCGGGTGGTGTGGAACTTG<br>CCGAGGCCGTGAGCCCGTGGCCTCCGGCCGCGGACGGGGTGCGCCGGGCCGGTGTGTC<br>GGCGTTCGGGGTGAGC**GGGACGAACGCGCACGTGATCATCGCCGAGCCCCCGGAGCCC**<br>**GAGCCGCTGCCGGAACCCGGACCGGTGGGCGTGCTGGCCGCTGCGAACTCGGTGCCCG**<br>**TACTGCTGTCGGCCAGGACCGAGACCGCGTTGGCAGCGCAGGCGCGGCTCCTGGAGTC**<br>**CGCAGTGGACGACTCGGTTCCGTTGACGGCATTGGCTTCCGCGCTGGCCACCGGACGC**<br>**GCCCACCTGCCGCGTCGTGCGGCGTTGCTGGCAGGCGACCACGAACAGCTCCGCGGGC**<br>**AGTTGCGAGCGGTCGCCGAGGGCGTTGCGGCTCCCGGTGCCACCACCGGAACCGCCTC**<br>**CGCCGGCGGCGTGGTTTTCGTCTTCCCAGGTCAGGGTGCTCAGTGGGAGGGCATGGCC**<br>**CGGGGCTTGCTCTCGGTCCCCGTCTTCGCCGAGTCGATCGCCGAGTGCGATGCGGTGT**<br>**TGTCGGAGGTGGCCGGGTTCTCGGCCTCCGAAGTGCTGGAGCAGCGTCCGGACGCGCC**<br>**GTCGCTGGAGCGGGTCGACGTCGTACAGCCGGTGTTGTTCTCCGTGATGGTGTCGCTG**<br>**GCGCGGCTGTGGGGCGCTTGCGGAGTCAGCCCCTCGGCCGTCATCGGCCATTCGCAGG**<br>**GCGAGATCGCCGCCGCGGTGGTGGCCGGGGTGTTGTCGCTGGAGGACGGCGTGCGCGT**<br>**CGTGGCCCTGCGCGCGAAGGCGTTGCGTGCGCTGGCGGGCAAGGGCGGCATGGTCTCG**<br>**TTGGCGGCTCCCGGTGAACGCGCCCGCGCGCTGATCGCACCGTGGGAGGACCGGATCT**<br>**CCGTCGCGGCGGTCAACTCCCCGTCCTCGGTCGTGGTCTCCGGCGATCCGGAGGCGCT**<br>**GGCCGAACTCGTCGCACGTTGCGAGGACGAGGGCGTGCGCGCCAAGACGCTCCCGGTG**<br>**GACTACGCCTCGCACTCCCGCCACGTCGAGGAGATCCGCGAGACGATCCTCGCCGACC**<br>**TCGACGGCATCTCCGCGCGGCGTGCCGCCATCCCGCTCTACTCCACGCTGCACGGCGA**<br>**ACGGCGCGACGGCGCCGACATGGGTCCGCGGTACTGGTACGACAACCTGCGCTCCCAG**<br>**GTGCGCTTCGACGAGGCGGTCTCGGCCGCCGTCGCCGACGGTCACGCCACCTTCGTCG**<br>**AGATGAGCCCGCACCCGGTGCTCACCGCGGCGGTGCAGGAGATCGCCGCGGACGCCGT**<br>**GGCCATCGGGTCGCTGCACCGCGACACCGCGGAGGAGCACCTGATCGCCGAGCTCGCC**<br>**CGGGCGCACGTGCACGGCGTGGCCGTGGACTGGCGGAACGTCTTCCCGGCGGCACCTC**<br>**CGGTGGCG**CTGCCCAACTACCCGTTCGAGCCCCAGCGGTACTGGCTCGCGCCGGAGGT<br>GTCCGACCAGCTCGCCGACAGCCGCTACCGCGTCGACTGGCGACCGCTGGCCACCACG<br>CCGGTGGACCTGGAAGGCGGCTTCCTGGTCCACGGGTCCGCACCGGAGTCGCTGACCA<br>GCGCAGTCGAGAAGGCCGGAGGCCGCGTCGTGCCGGTCGCCTCGGCCGACCGCGAAGC<br>GCTCGCGGCGGCCCTGCGGGAGGTGCCGGGCGAGGTCGCCGGCGTGCTCTCGGTCCAC<br>ACCGGCGCCGCAACGCACCTCGCCCTGCACCAGTCGCTGGGTGAGGCCGGCGTGCGGG<br>CCCCGCTCTGGCTGGTCACCAGCCGAGCGGTCGCGCTCGGGGAGTCCGAGCCGGTCGA<br>TCCCGAGCAGGCGATGGTGTGGGGTCTCGGGCGCGTCATGGGCCTGGAGACCCCGGAA<br>CGGTGGGGCGGTCTGGTGGACCTGCCCGCCGAACCCGCGCCGGGGGACGGCGAGGCGT<br>TCGTCGCCTGCCTCGGCGCGGACGGCCACGAGGACCAGGTCGCGATCCGTGACCACGC<br>CCGCTACGGCCGCCGCCTCGTCCGCGCCCCGCTGGGCACCCGCGAGTCGAGCTGGGAG<br>CCGGCGGGCACGGCGCTGGTCACCGGCGGCACCGGTGCGCTCGGCGGCCACGTCGCCC<br>GCCACCTCGCCAGGTGCGGGGTGGAGGACCTGGTGCTGGTCAGCAGGCGCGGCGTCGA<br>CGCTCCCGGCGCGGCCGAGCTGGAAGCCGAACTGGTCGCCCTCGGCGCGAAGACGACC<br>ATCACCGCCTGCGACGTGGCCGACCGCGAGCAGCTCTCCAAGCTGCTGGAAGAACTGC |

| | |
|---|---|
| | GCGGGCAGGGACGTCCGGTGCGGACCGTCGTGCACACCGCCGGGGTGCCCGAATCGAG<br>GCCGCTGCACGAGATCGGCGAGCTGGAGTCGGTCTGCGCGGCGAAGGTGACCGGGGCC<br>CGGCTGCTCGACGAGCTGTGCCCGGACGCCGAGACCTTCGTCCTGTTCTCGTCCGGAG<br>CGGGGGTGTGGGGCAGTGCGAACCTCGGCGCCTACTCCGCGGCCAACGCCTACCTCGA<br>CGCGCTGGCCCACCGCCGCCGTGCGGAAGGCCGTGCGGCGACGTCCGTCGCGTGGGGC<br>GCCTGGGCGGGCGAGGGCATGGCCACCGGCGACCTCGAGGGGCTCACCCGGCGCGGCC<br>TGCGCCCGATGGCGCCCGAGCGCGCGATCCGCGCGCTGCACCAGGCGCTGGACAACGG<br>CGACACGTGCGTTTCGATCGCCGACGTCGACTGGGAGCGCTTCGCGGTCGGCTTCACC<br>GCCGCCCGGCCGCGTCCGCTGCTGGACGAGCTCGTCACGCCGGCGGTGGGGGCCGTCC<br>CCGCGGTGCAGGCGGCCCCGGCGCGGGAGATGACGTCGCAGGAGTTGCTGGAGTTCAC<br>GCACTCGCACGTCGCGGCGATCCTCGGGCATTCCAGCCCGGACGCGGTCGGGCAGGAC<br>CAGCCGTTCACCGAGCTCGGCTTCGACTCGCTGACCGCGGTCGGGCTGCGCAACCAGC<br>TCCAGCAGGCCACCGGGCTCGCGCTGCCCGCGACCCTGGTGTTCGAGCACCCCACGGT<br>CCGCAGGTTGGCCGACCACATAGGACAGCAGCTCGACAGCGGGACTCCCGCCCGGGAA<br>GCGAGCAGCGCTCTTCGCGACGGCTACCGGCAGGCGGGCGTGTCGGGCAGGGTCCGGT<br>CCTACCTCGACCTGCTGGCGGGGCTGTCGGACTTCCGCGAGCACTTCGACGGCTCCGA<br>CGGGTTCTCCCTCGATCTCGTGGACATGGCCGACGGTCCCGGAGAGGTCACGGTGATC<br>TGCTGCGCGGGAACGGCGGCGATCTCCGGTCCGCACGAGTTCACCCGGCTCGCCGGGG<br>CGCTGCGCGGAATCGCTCCGGTTCGGGCCGTGCCCCAGCCCGGCTACGAGGAGGGCGA<br>ACCTCTGCCGTCGTCGATGGCGGCGGTGGCGGCGGTGCAGGCCGATGCGGTCATCAGG<br>ACACAGGGGGACAAGCCGTTCGTGGTGGCCGGTCACTCCGCGGGGGCACTGATGGCCT<br>ACGCGCTGGCGACCGAACTGCTCGATCGCGGGCACCCGCCACGCGGTGTCGTCCTGAT<br>CGACGTCTACCCGCCCGGTCACCAGGACGCGATGAACGCCTGGCTGGAGGAGCTGACC<br>GCCACGCTGTTCGACCGCGAGACGGTGCGGATGGACGACACCAGGCTCACCGCCCTGG<br>GCGCCTACGACCGCCTCACCGGTCAGTGGCGACCCCGGGAAACCGGGCTGCCGACGCT<br>GCTGGTCAGCGCCGGCGAGCCGATGGGTCCGTGGCCCGACGACAGCTGGAAGCCGACG<br>TGGCCCTTCGAGCACGACACCGTCGCCGTCCCCGGCGACCACTTCACGATGGTGCAGG<br>AACACGCCGACGCGATCGCGCGGCACATCGACGCCTGGCTGGGCGGAGGGAATTCAAG<br>A |

**Table S7. Primers for DEBS PKS construction and mutagenesis.** GA = Gibson Assembly. RTH = 'Round-the-horn mutagenesis.[1] RTH primers have 5' phosphates.

| Primer Name | 5'→3' Primer Sequence | Primer Function |
|---|---|---|
| AT_Motif.Gib1<br>AT_Motif.Gib2 | CACTCCCGCCACGTCGAG<br>CTCCAGCGACGGCGCG | Amplifies Ery6TE without AT motifs for GA |
| AT_Motif.FOR<br>AT_Motif.REV | ACGCGCCGTCGCTGG<br>TCGACGTGGCGGGAGTG | Amplifies motif fragments for GA |
| Ery6V742A.FOR<br>Ery6V742A.REV | GCGGACTACGCCTCGCACTCCC<br>CGGGAGCGTCTTGGCGC | Ery6 V742A RTH |
| Ery6Y744RGV.RTH1<br>Ery6Y744RGV.RTH2 | SKGGCCTCGCACTCCCGC<br>GTCCACCGGGAGCGTCTTG | Ery6TE Y744R/G RTH |
| Ery6L676V.FOR<br>Ery6L676V.REV | GTGGCGGGCAAGGGCG<br>CGCACGCAACGCCTTCG | Ery6 L676V RTH |
| Ery6L673G.FOR<br>Ery6L673G.REV | GGACGTGCGCTGGCGGG<br>CGCCTTCGCGCGCAG | Ery6 L673G RTH |
| Ery6E647D.FOR<br>Ery6E647D.REV | GATATCGCCGCCGCGG<br>GCCCTGCGAATGGCCGAT | Ery6 E647D RTH |
| Ery6L797X.FOR<br>Ery6L797X.REV | NNKCGCTCCCAGGTGCGC<br>GTTGTCGTACCAGTACCGCGGA | Ery6 L797X RTH |
| Ery6I648A.FOR<br>Ery6I648X.REV | GCCGCCGCGGTGGTGG<br>GNSCTCGCCCTGCGAATGGC | Ery6 I648G/A/V RTH |
| Ery6H643F.FOR<br>Ery6H643F.REV | TTTTCGCAGGGCGAGATCGC<br>GCCGATGACGGCCGAGG | Ery6 H643F RTH |
| Ery6H643S.FOR<br>Ery6H643F.REV | TCTTCGCAGGGCGAGATCGC<br>GCCGATGACGGCCGAGG | Ery6 H643S RTH |
| Ery6L740V.FOR<br>Ery6L740V.REV | GTCCCGGTGGACTACGCCTC<br>CGTCTTGGCGCGCACG | Ery6 L740V RTH |
| Ery6A707G.FOR<br>Ery6A707G.REV | GGGGTCAACTCCCCGTCCTCG<br>CGCGACGGAGATCCGGTC | Ery6 A707G RTH |
| Ery6G681A.FOR<br>Ery6G681A.REV | GCAATGGTCTCGTTGGCGGC<br>GCCCTTGCCCGCCAGC | Ery6 G681A RTH |
| Ery6V751A.FOR<br>Ery6V751A.REV | GCCGAGGAGATCCGCGAGAC<br>GTGGCGGGAGTGCGAGG | Ery6 V751A RTH |
| Ery6S644A.FOR<br>Ery6S644A.REV | GTCAGCCGTTATCGGTCATGCT<br>CCGCAATTTCGCCCTG | Ery6 S644A RTH |
| Ery6D613E.FOR<br>Ery6D613E.REV | GAAGTCGTACAGCCGGTGTTG<br>GACCCGCTCCAGCGACG | Ery6 D613E RTH |
| Ery6V751A.FOR<br>Ery6V751A.REV | GCCGAGGAGATCCGCGAGAC<br>GTGGCGGGAGTGCGAGG | Ery6 V751A RTH |

**Table S8. High-resolution LC-MS parameters and gradient.**

| HESI Source Parameters | |
|---|---|
| Spray Voltage | 3.5 kV |
| Capillary Temperature | 350 °C |
| Heater Temperature | 300 °C |
| S Lens RF Level | 70 V |
| Sheath Gas Flow Rate | 60 au |
| Resolution | 70,000 FWHM |
| Scan Range | 100-1000 m/z |

**LC Gradient**

| Time (min) | % B |
|---|---|
| 0.0 | 25.0 |
| 1.0 | 25.0 |
| 10.0 | 59.0 |
| 11.0 | 85.0 |
| 12.0 | 85.0 |
| 12.5 | 25.0 |
| 16.5 | 25.0 |

### Supplementary Figures

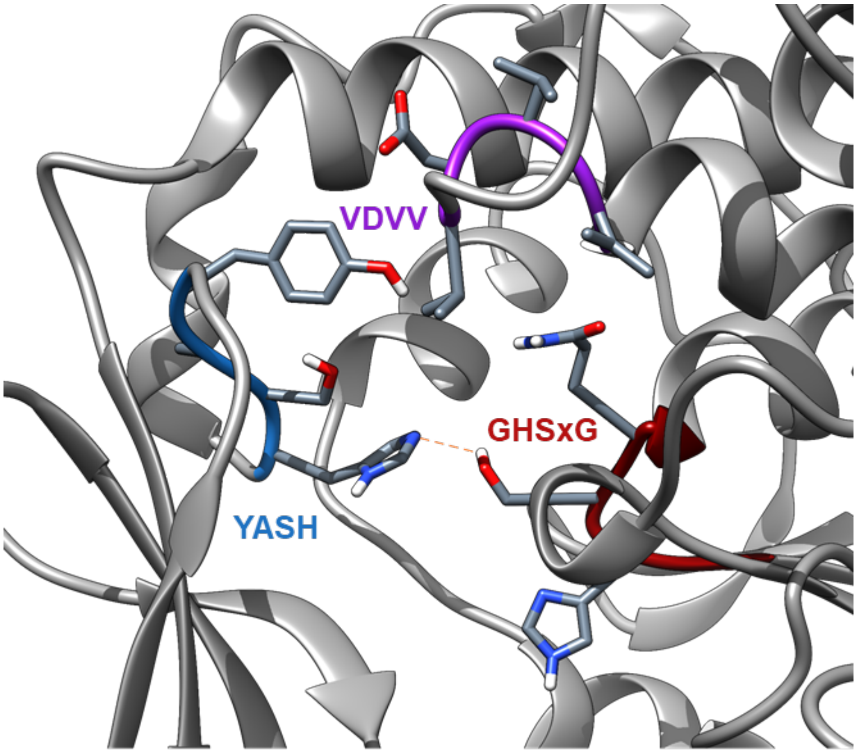


**Figure S1. Conserved motifs of PKS acyltransferases.** Active site of EryAT6 with well-conserved motifs from methylmalonyl-CoA-specific ATs depicted as sticks (SeqLogos in **Figure S10**).

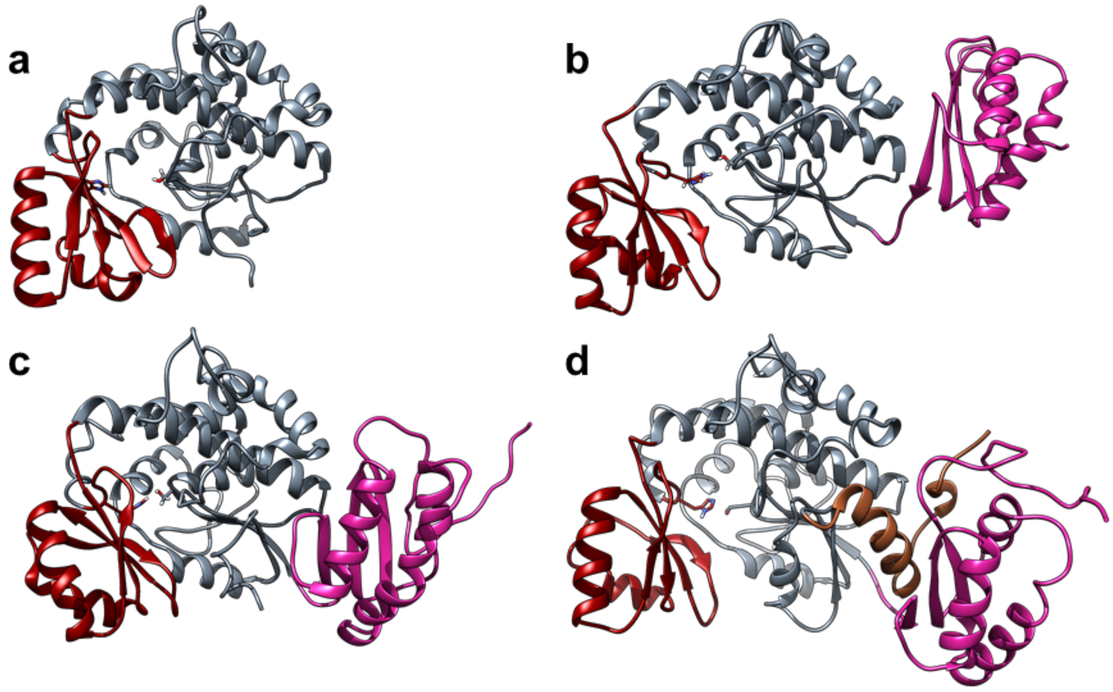


**Figure S2. Evaluation of EryAT6 models.** EryAT6 homology model simulated with (**a**) no linkers and Nε-protonated His747 (278 residues; average Ser644-His747 distance of 8.24 Å), (**b**) an N-terminal linker (magenta) and Nε-protonated His747 (387 residues; average Ser644-His747 distance of 4.13 Å), (**c**) an N-terminal linker (magenta) and Nδ-protonated His747 (387 residues; average Ser644-His747 distance of 3.03 Å), and (**d**) an N-terminal linker (magenta), C-terminal linker (brown), and Nδ-protonated His747 (430 residues; average Ser644-His747 distance of 3.03 Å). This final model also showed improved longevity of secondary structures. The small (red) and large (grey) subunits are included in all models, with the catalytic dyad depicted as sticks.

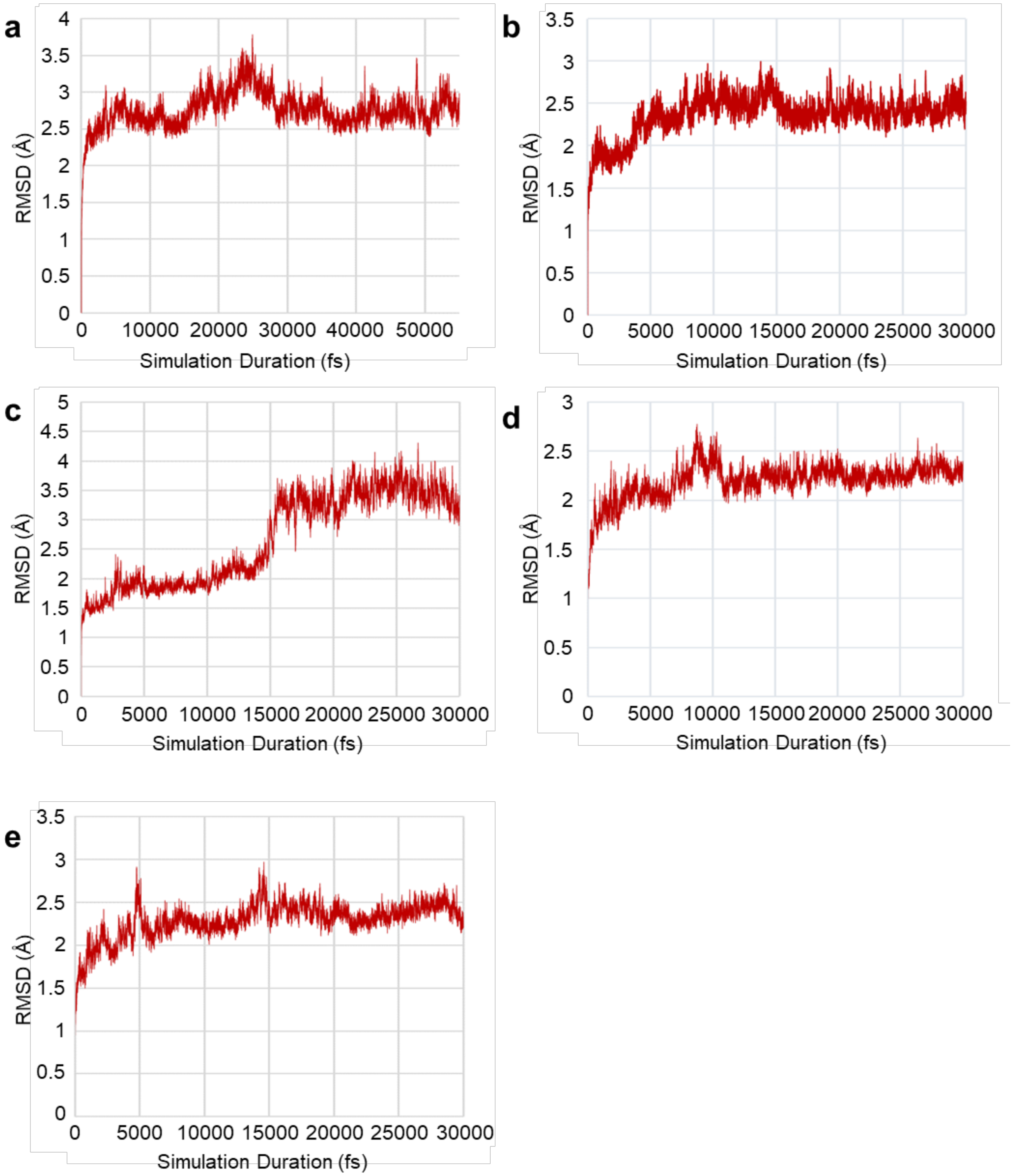


**Figure S3. Structural convergence of EryAT6 MD simulations.** RMSD of the Cα from all residues within the AT (excluding linkers) for each of the EryAT6 models: (**a**) wild-type. (**b**) V742A. (**c**) Y744R. (**d**) L673H. (**e**) V742A/Y744R.

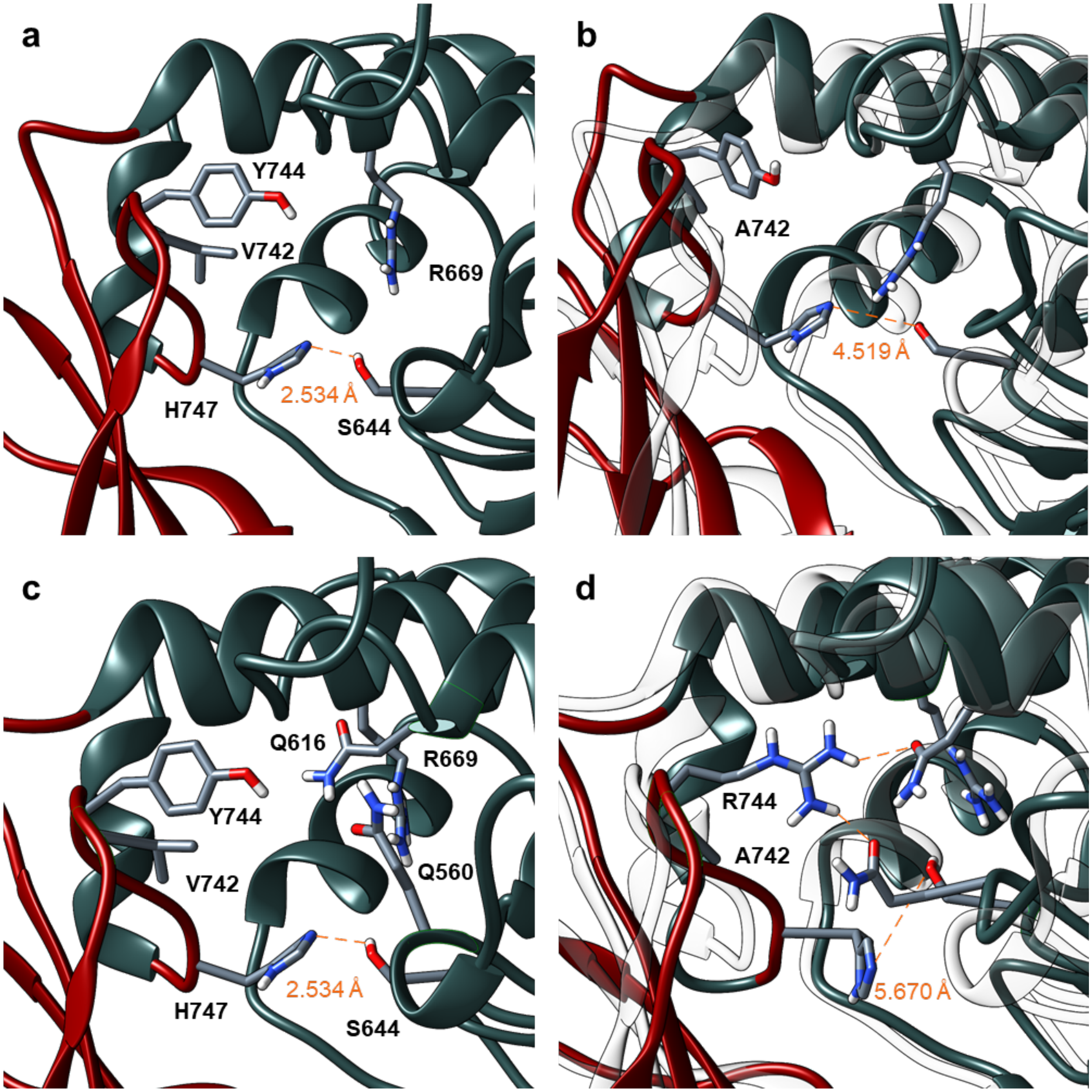


**Figure S4. Snapshots of EryAT6 individual mutations.** (**a**) Wild-type EryAT6 typically maintains a catalytic dyad distance under 4 Å, in part because of the hydrophobic interaction of Val742 with the large subunit. (**b**) In the V742A MD simulation, significant flexibility has been introduced into the small subunit loop, with a concomitant decrease in a hydrophobic interaction with the large subunit, allowing access to longer catalytic dyad distances. (**c**) In wild-type EryAT6, Gln560 and Gln616 help to form the oxyanion hole required for catalysis; however, in the double V742A/Y744R mutant, Arg744 interacts with both glutamines (**d**). The entirety of the double mutant simulation was highly unstable, consistent with its poor *in vitro* activity.

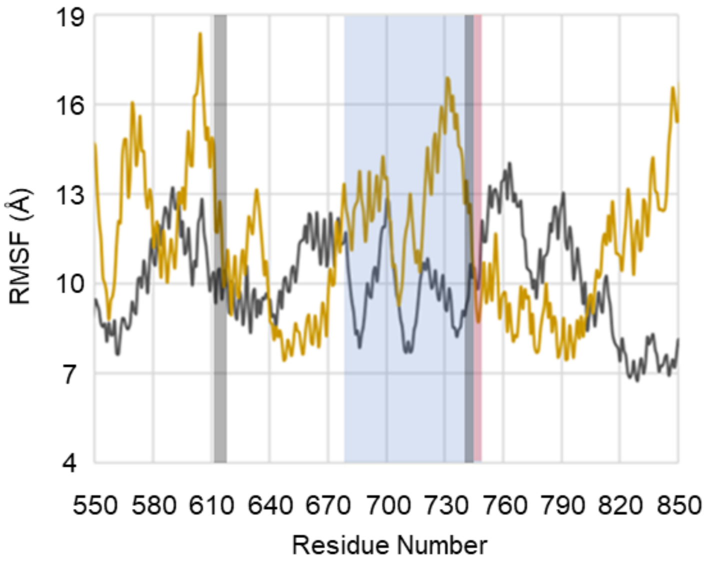


**Figure S5. RMSF plot for EryAT6 V742A/Y744R simulation.** Root-mean-square fluctuation (RMSF) of the 60 ns wild-type (grey) and the 30 ns V742A/Y744R (gold) MD simulation. Higher values correspond to increased movement of a residue over the time frame. The blue box represents the residues found in the small subunit of the AT. The red box highlights the location of the YASH motif, and the grey boxes describe the locations of the SSM, including the YASH motif, and the LSM. Linkers were excluded from the analyses.

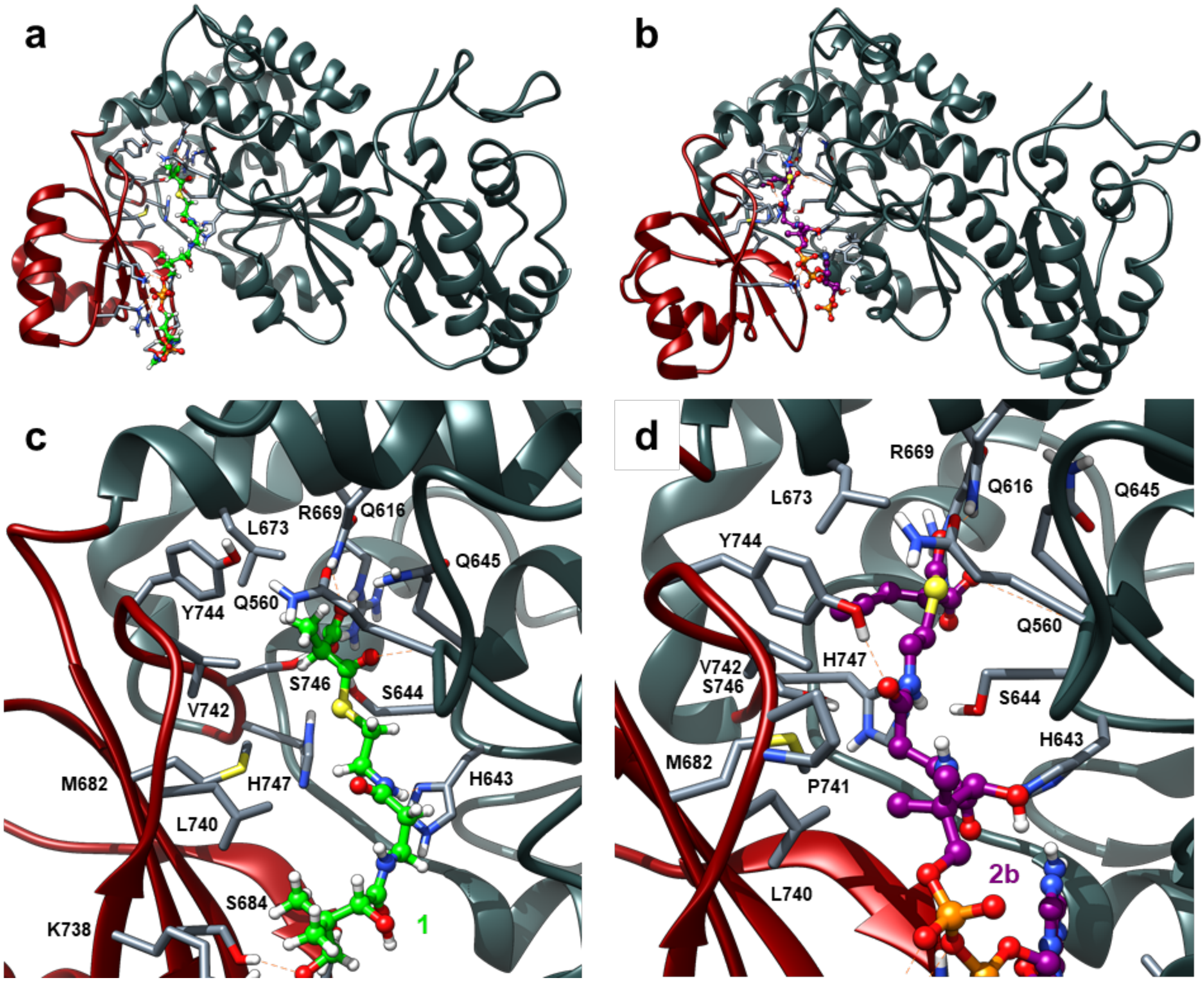


**Figure S6. EryAT6 wild-type with methylmalonyl-CoA and propargylmalonyl-CoA.** (**a**) Overview of the whole EryAT6 wild-type model with mmCoA (**1**, green) in the active site after a 10 ns MD simulation. The small subunit is shown in red, and the large subunit is depicted in grey. (**b**) Overview of the whole EryAT6 wild-type model with pgmCoA (**2b**, purple) in the active site after a 10 ns MD simulation. (**c**) A close-up view of the active site of EryAT6 wild-type with mmCoA (**1**, green) bound. Most residues predicted to directly interact with the substrate are indicated. (**d**) A close-up view of the active site of EryAT6 wild-type with pgmCoA (**2b**, purple) bound. Most residues predicted to directly interact with the substrate are indicated. Relative to the mmCoA-bound model, the pgmCoA takes on a less ideal conformation for attack by the catalytic Ser644.

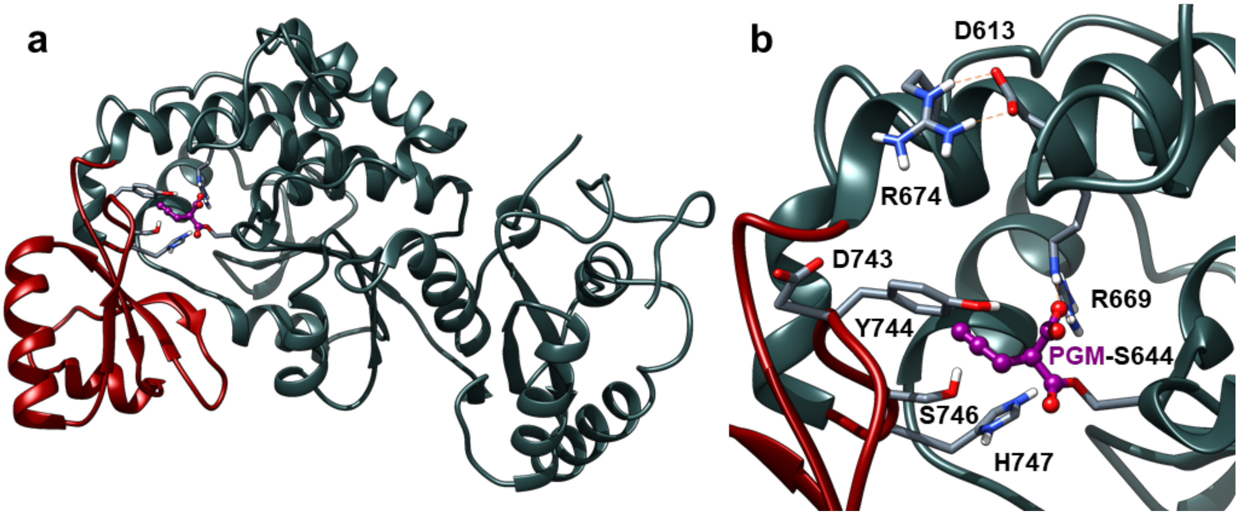


**Figure S7. EryAT6 wild-type with propargylmalonated Ser644.** (**a**) Overall architecture of EryAT6 did not change over 10 ns MD simulation with a covalently-linked propargylmalonate (PGM) group on the catalytic Ser644 (purple). (**b**) The larger size of the propargylmalonate (purple) results in an active site that cannot retain the narrower active site required for the salt bridge network between Asp743, Arg674, and Asp613 that is found in the unbound form.

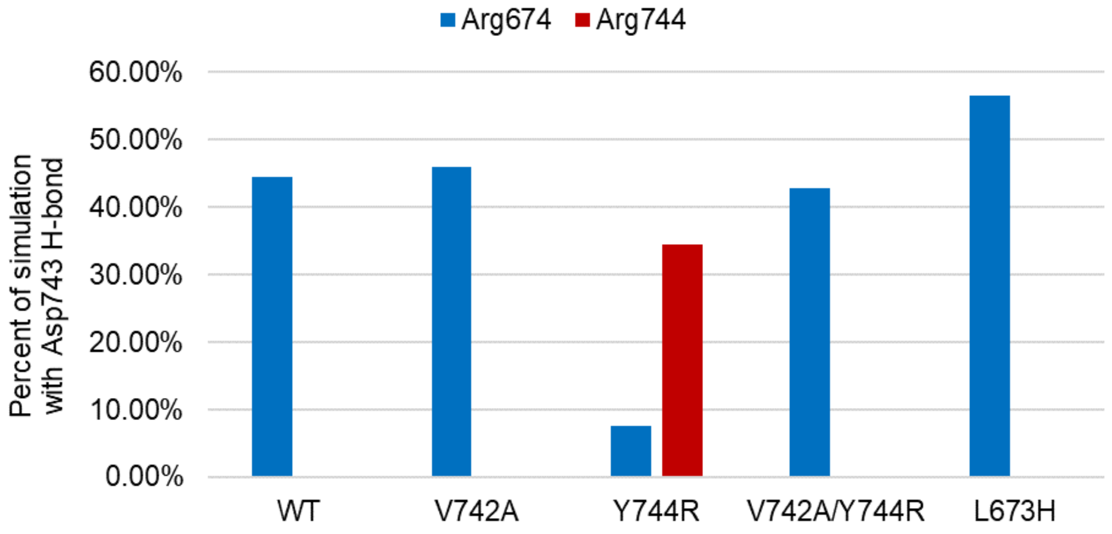


**Figure S8. Retention of Asp743 salt bridge.** The percentage of frames from each simulation in which a salt bridge is observed with Asp743 and either Arg674 or Arg744. In the EryAT6 Y744R simulation, the intra-subunit salt bridge between Asp743 and Arg674 is nearly abolished. In the EryAT6 L673H simulation, a significant increase in presence of the salt bridge is observed, likely due to the narrower active site of this mutant. An interaction between Asp743 and Arg744 is seen in less than 1% of the V742A/Y744R simulation.

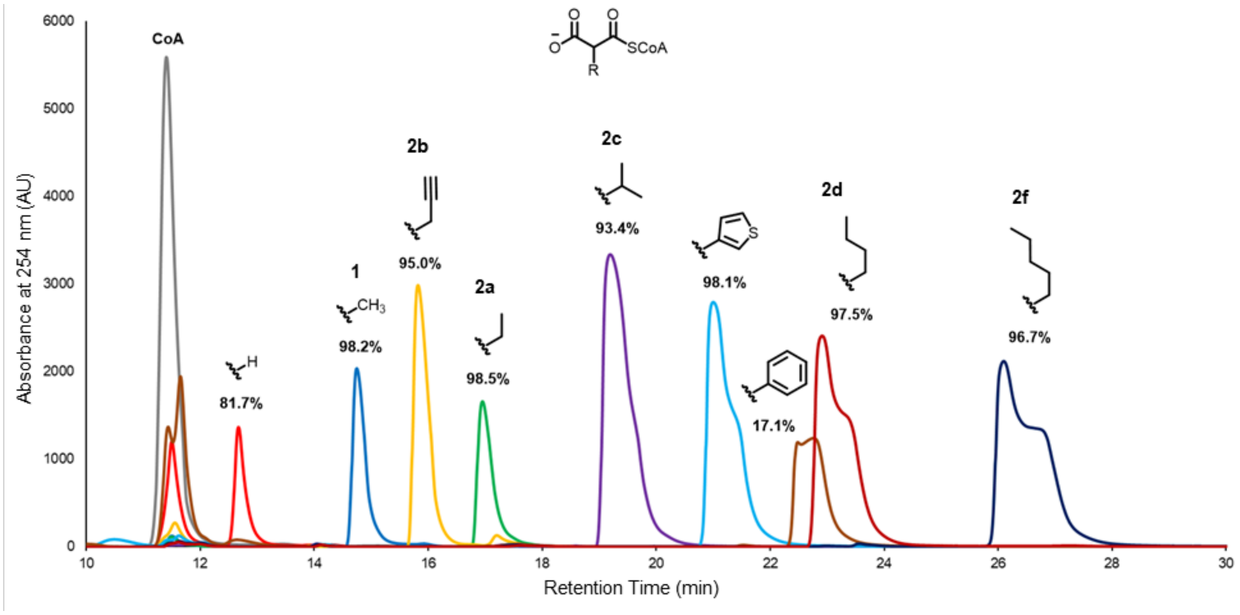


**Figure S9. Representative HPLC traces for MatB-synthesized malonyl-CoAs.** Representative HPLC traces for MatB-synthesized malonyl-CoAs. Percent conversions were determined by remaining CoA peak.

**a**

| (2S)-X-malonyl-CoA | Acyltransferase | Small Subunit Motif | Large Subunit Motif |
|---|---|---|---|
| H | Avermectin AT5 | TLP-TNHAFH | QTPYAQP |
| Methyl | Erythromycin AT6 | TLP-VDYASH | RVDVVQP |
| Methyl | Rifamycin AT3 | RVA-VDYASH | RVDVVQP |
| Ethyl/Methyl | Monensin AT5 | AVA-SDVAGH | RIDVVQP |
| Ethyl | Niddamycin AT5 | PIPGVDTAGH | RVDVVQP |
| Allyl/Ethyl | FK506 AT4 | RIA-VDCPTH | RVDVVHP |
| Chloroethyl | Salinosporamide AT1 | AVR-VDRPGH | ATDIQQP |
| Butyl | Thailandin AT13 | VLPGADAAGH | RLDVNQP |
| Hexyl | Cinnabaramide AT1 | ALR-VERAGH | EGDIQQP |
| Benzyl | Splenocin AT1 | RLR-MPAAAH | --TAAFM |

**b**

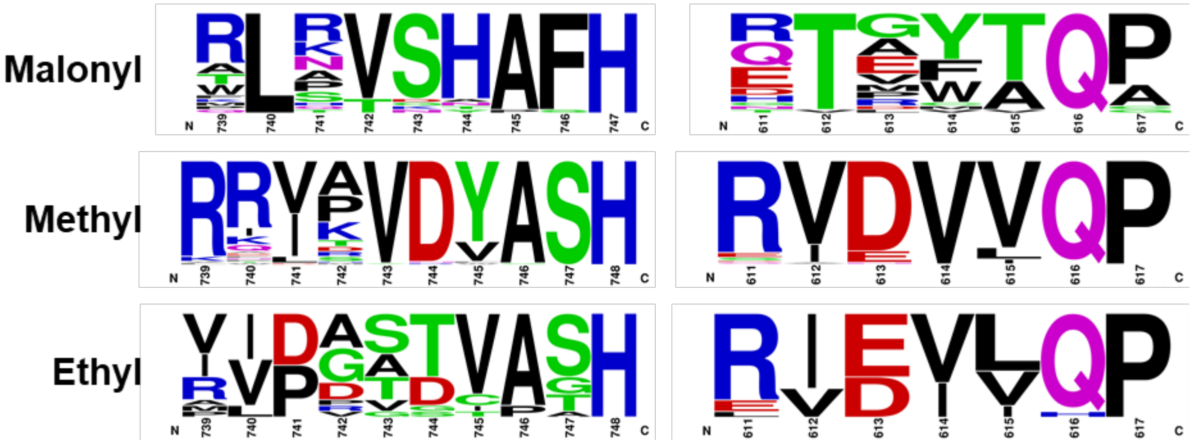


**Figure S10. Motif sequences from natural acyltransferases.** (**a**) Selected examples of SSM and LSM sequences for ATs with different CoA-linked substrates. (**b**) Consensus sequences of AT domains with substrate specificities for malonyl-CoA (100 sequences), methylmalonyl-CoA (100 sequences), and ethylmalonyl-CoA (19 sequences) are shown. Sequence logos were created using WebLogo.[6]

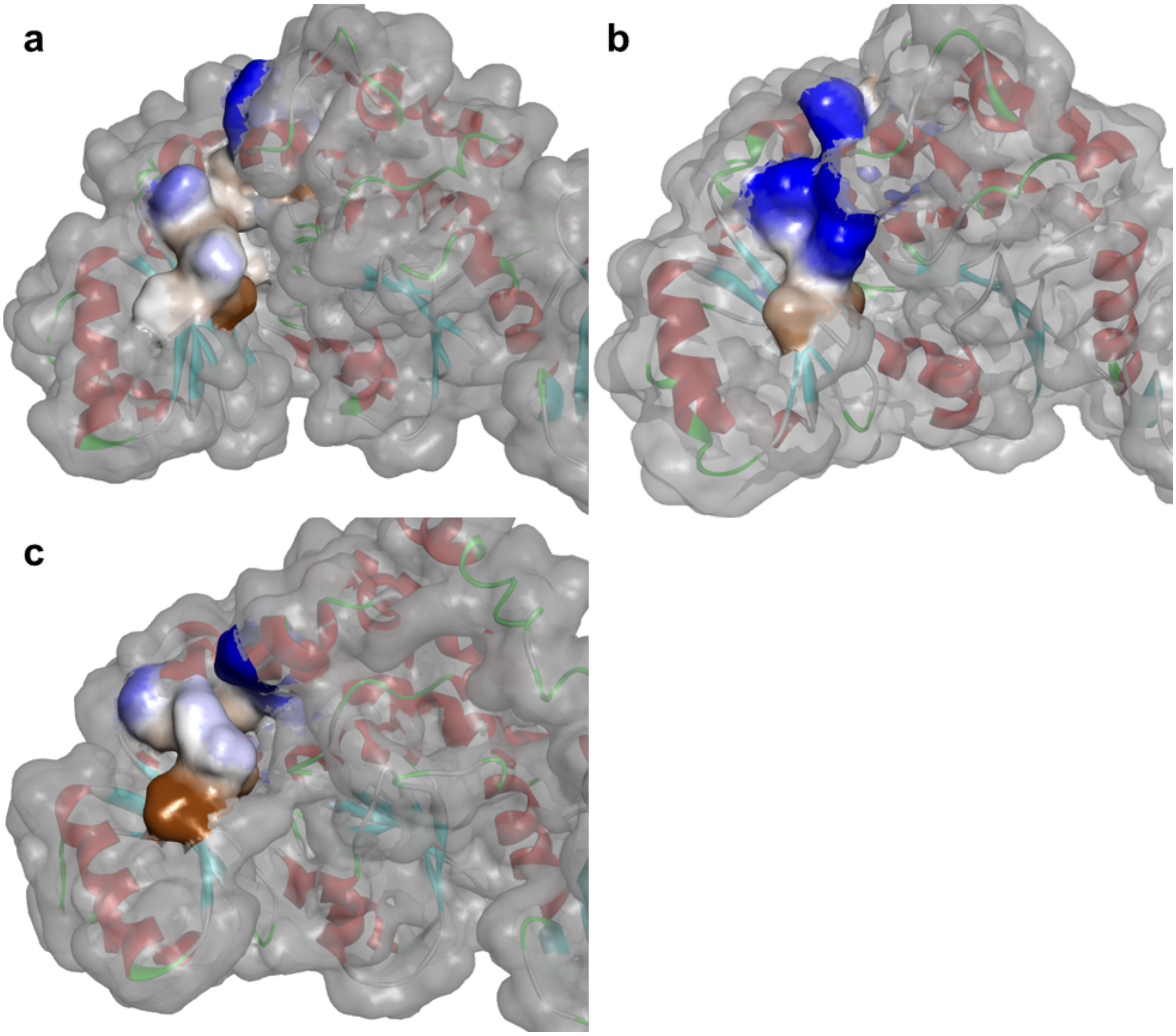


**Figure S11. Surface models of wild-type and motif-swapped EryAT6.** Surface depiction of EryAT6 models with the two motifs colored by hydrophobicity (blue for hydrophilic; brown for hydrophobic). (**a**) EryAT6 wild-type shows mostly hydrophobic interactions between the two motifs. (**b**) EryAT6 with CinAT1 motifs replaces those interactions with mostly charged residues. (**c**) EryAT6 with ThaAT13 motifs utilize similar hydrophobic interactions as those found in the wild-type enzyme. In each panel, the small subunit is on the left.

**Figure S12. Representative mass spectra of 10-deoxymethynolide B compounds from lysate module reactions.** Top panel for each compound shows the extracted ion chromatogram. Bottom panel(s) show the total ion spectra.

**a**

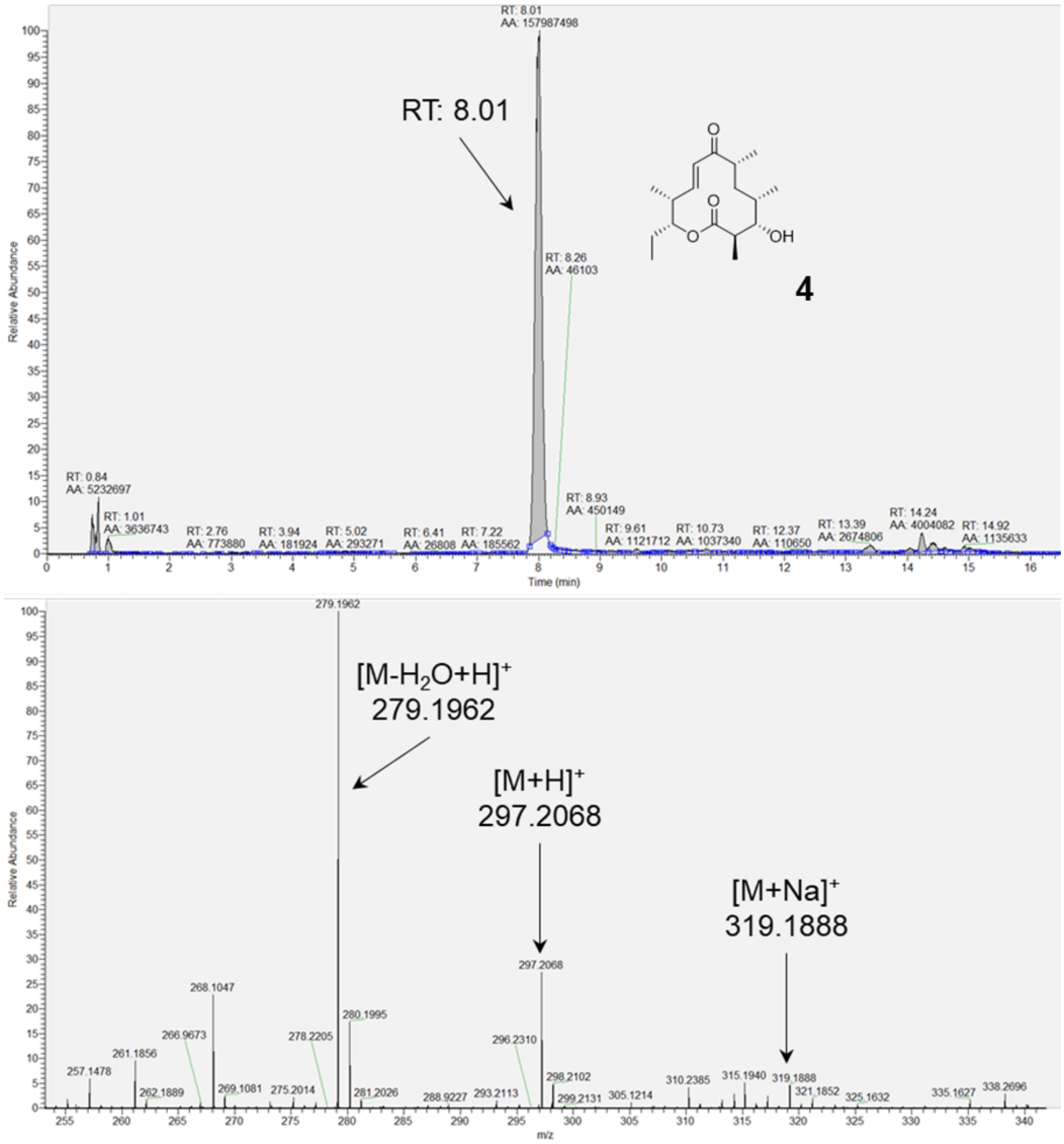

**b**

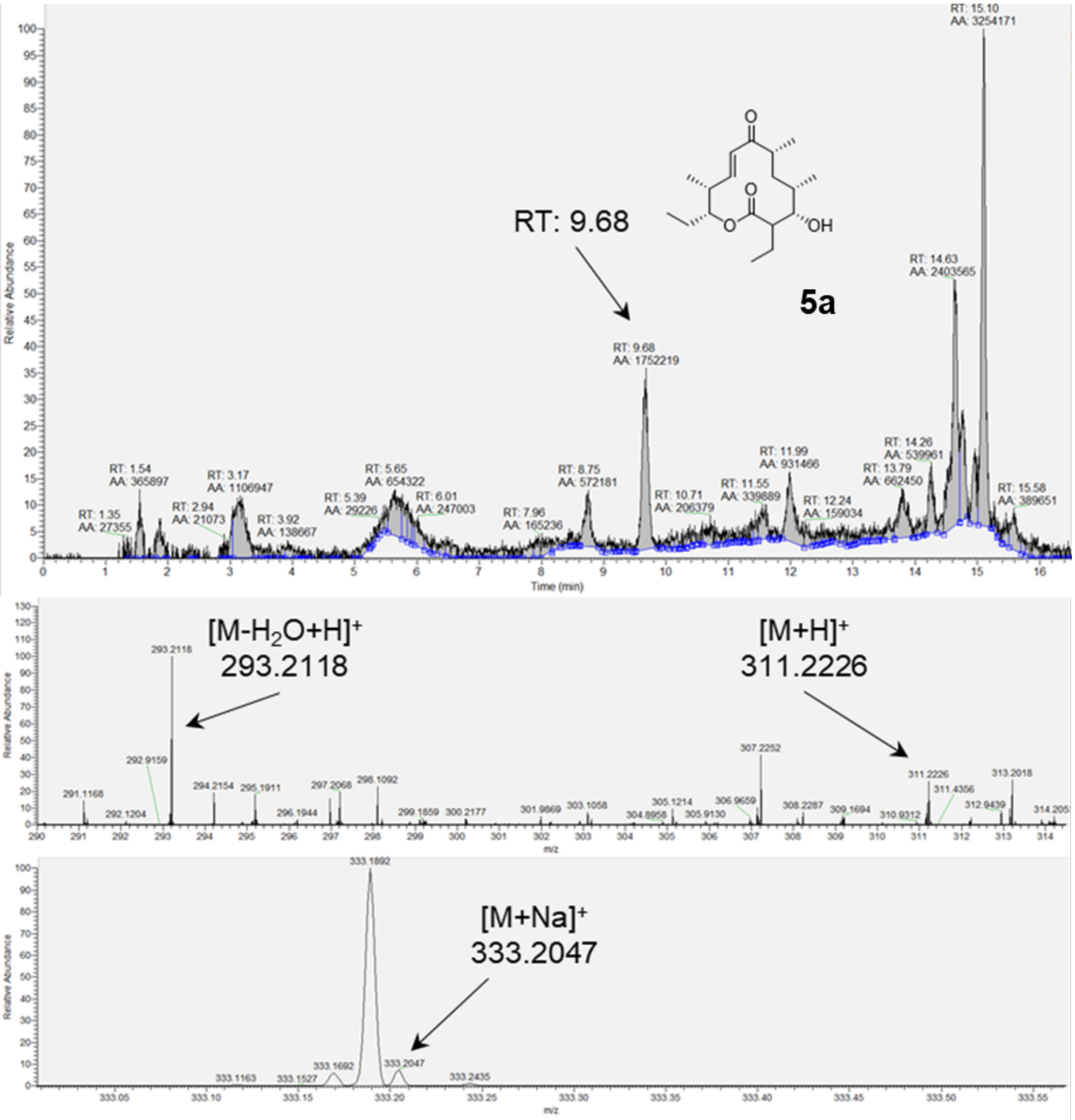
RT: 9.68
5a
OH
Relative Abundance
Time (min)
RT: 1.35 AA: 27355
RT: 1.54 AA: 365897
RT: 2.94 AA: 21073
RT: 3.17 AA: 1106947
RT: 3.92 AA: 138667
RT: 5.39 AA: 29226
RT: 5.65 AA: 654322
RT: 6.01 AA: 247003
RT: 7.96 AA: 165236
RT: 8.75 AA: 572181
RT: 9.68 AA: 1752219
RT: 10.71 AA: 206379
RT: 11.55 AA: 339889
RT: 11.99 AA: 931466
RT: 12.24 AA: 159034
RT: 13.79 AA: 662450
RT: 14.26 AA: 539961
RT: 14.63 AA: 2403565
RT: 15.10 AA: 3254171
RT: 15.58 AA: 389651
[M-H$_2$O+H]$^+$
293.2118
[M+H]$^+$
311.2226
m/z
[M+Na]$^+$
333.2047
333.1892
333.1692
333.2047
m/z

**c**

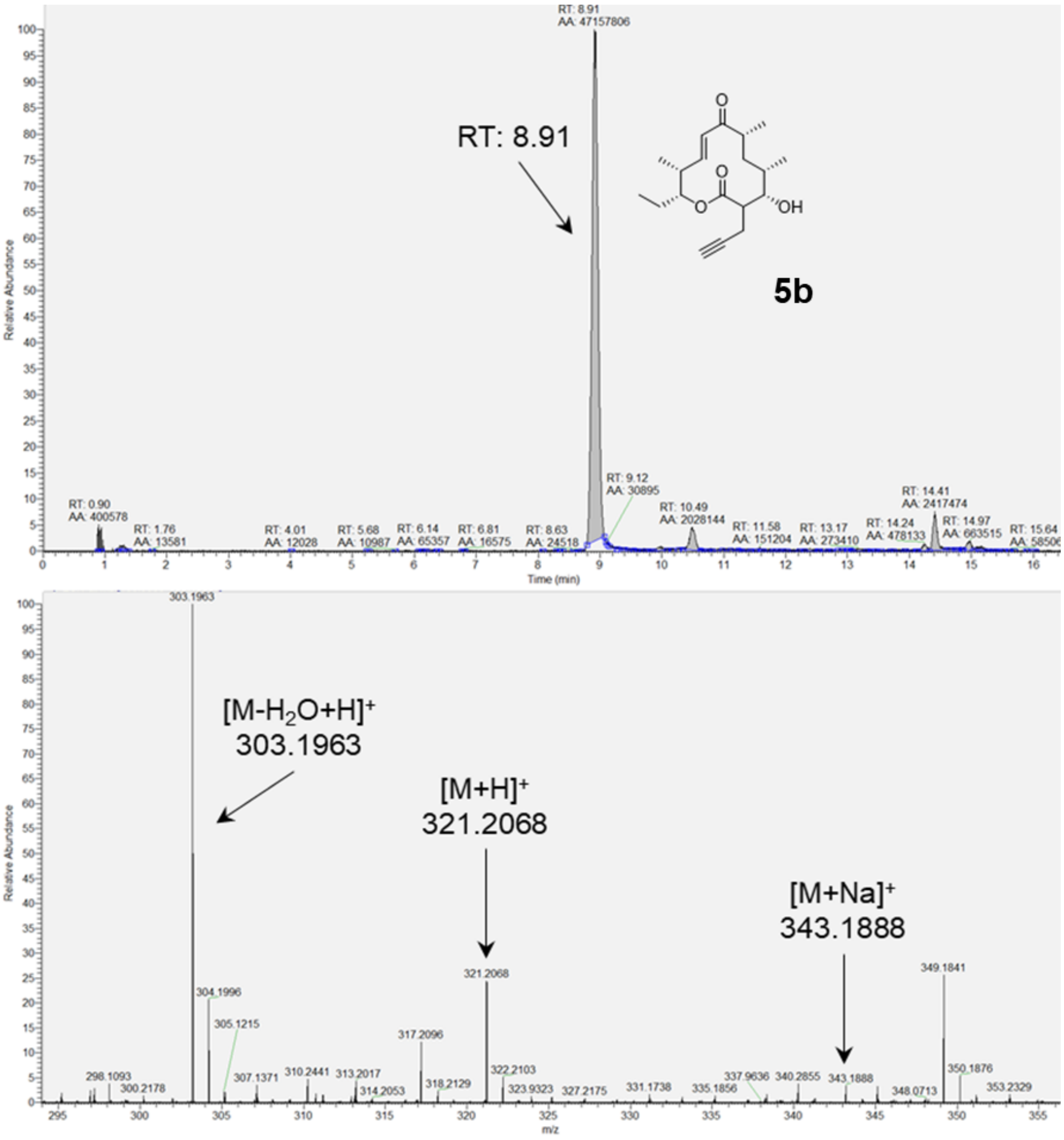
RT: 8.91
AA: 47157806
RT: 8.91
5b
OH
O
Relative Abundance
RT: 0.90
AA: 400578
RT: 1.76
AA: 13581
RT: 4.01
AA: 12028
RT: 5.68
AA: 10987
RT: 6.14
AA: 65357
RT: 6.81
AA: 16575
RT: 8.63
AA: 24518
RT: 9.12
AA: 30895
RT: 10.49
AA: 2028144
RT: 11.58
AA: 151204
RT: 13.17
AA: 273410
RT: 14.24
AA: 478133
RT: 14.41
AA: 2417474
RT: 14.97
AA: 663515
RT: 15.64
Time (min)
303.1963
[M-H$_2$O+H]$^+$
303.1963
[M+H]$^+$
321.2068
[M+Na]$^+$
343.1888
304.1996
305.1215
298.1093
300.2178
307.1371
310.2441
313.2017
314.2053
317.2096
318.2129
321.2068
322.2103
323.9323
327.2175
331.1738
335.1856
337.9636
340.2855
343.1888
348.0713
349.1841
350.1876
353.2329
m/z

d

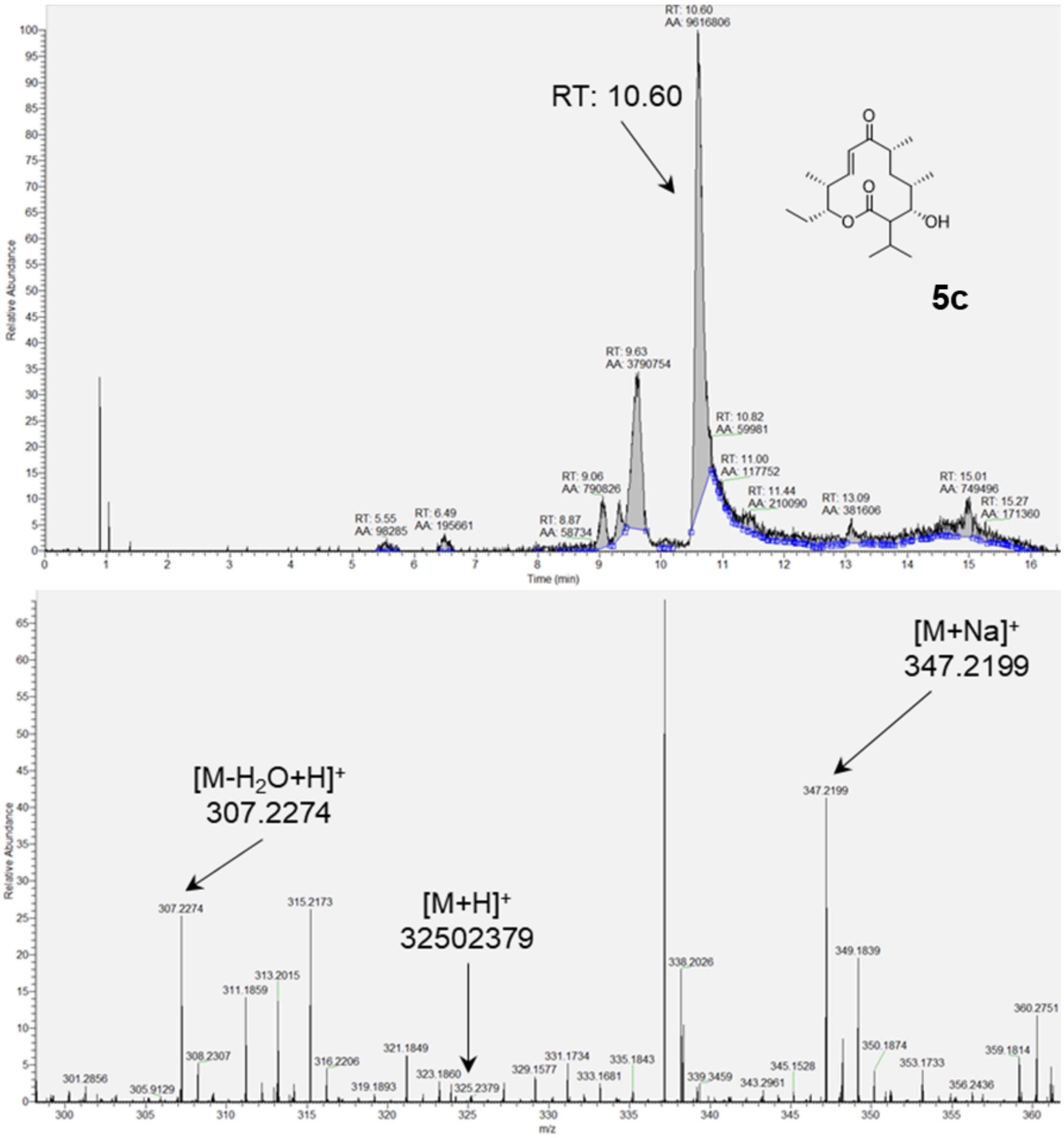

RT: 10.60
AA: 9616806
RT: 10.60
5c
OH
RT: 9.63
AA: 3790754
RT: 10.82
AA: 59981
RT: 11.00
AA: 117752
RT: 11.44
AA: 210090
RT: 9.06
AA: 790826
RT: 8.87
AA: 58734
RT: 5.55
AA: 98285
RT: 6.49
AA: 195661
RT: 13.09
AA: 381606
RT: 15.01
AA: 749496
RT: 15.27
AA: 171360
Relative Abundance
Time (min)
[M+Na]+
347.2199
[M-H2O+H]+
307.2274
[M+H]+
32502379
307.2274
315.2173
313.2015
311.1859
308.2307
316.2206
301.2856
305.9129
319.1893
321.1849
323.1860
325.2379
329.1577
331.1734
333.1681
335.1843
338.2026
339.3459
343.2961
345.1528
347.2199
349.1839
350.1874
353.1733
356.2436
359.1814
360.2751
Relative Abundance
m/z

e

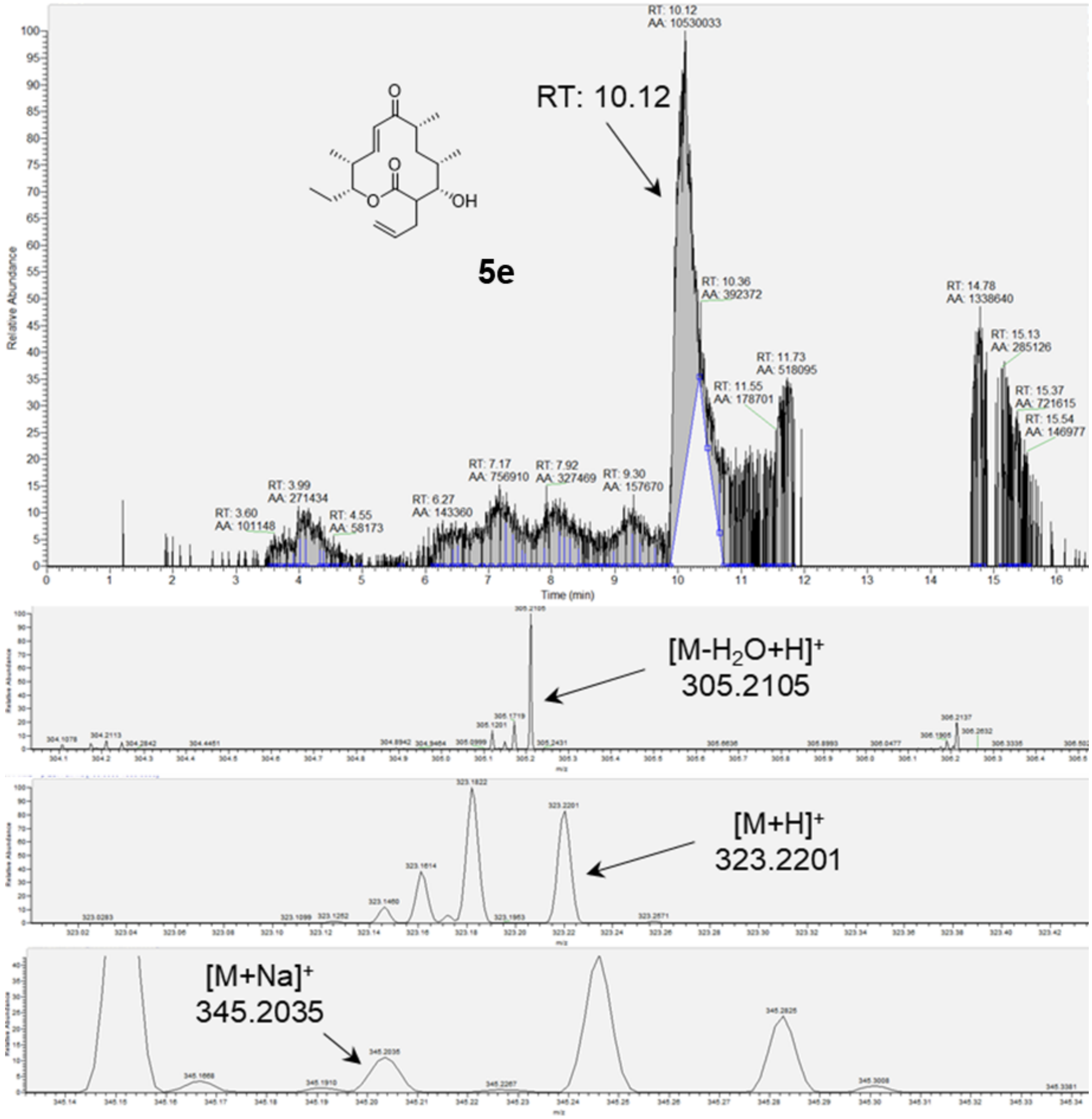

RT: 10.12
5e
[M-H2O+H]+
305.2105
[M+H]+
323.2201
[M+Na]+
345.2035
Relative Abundance
Time (min)

**Figure S13. Representative mass spectra of keto-10-deoxymethynolide B compounds from lysate module reactions.** Top panel for each compound shows the extracted ion chromatogram. Bottom panel(s) show the total ion spectra.

a

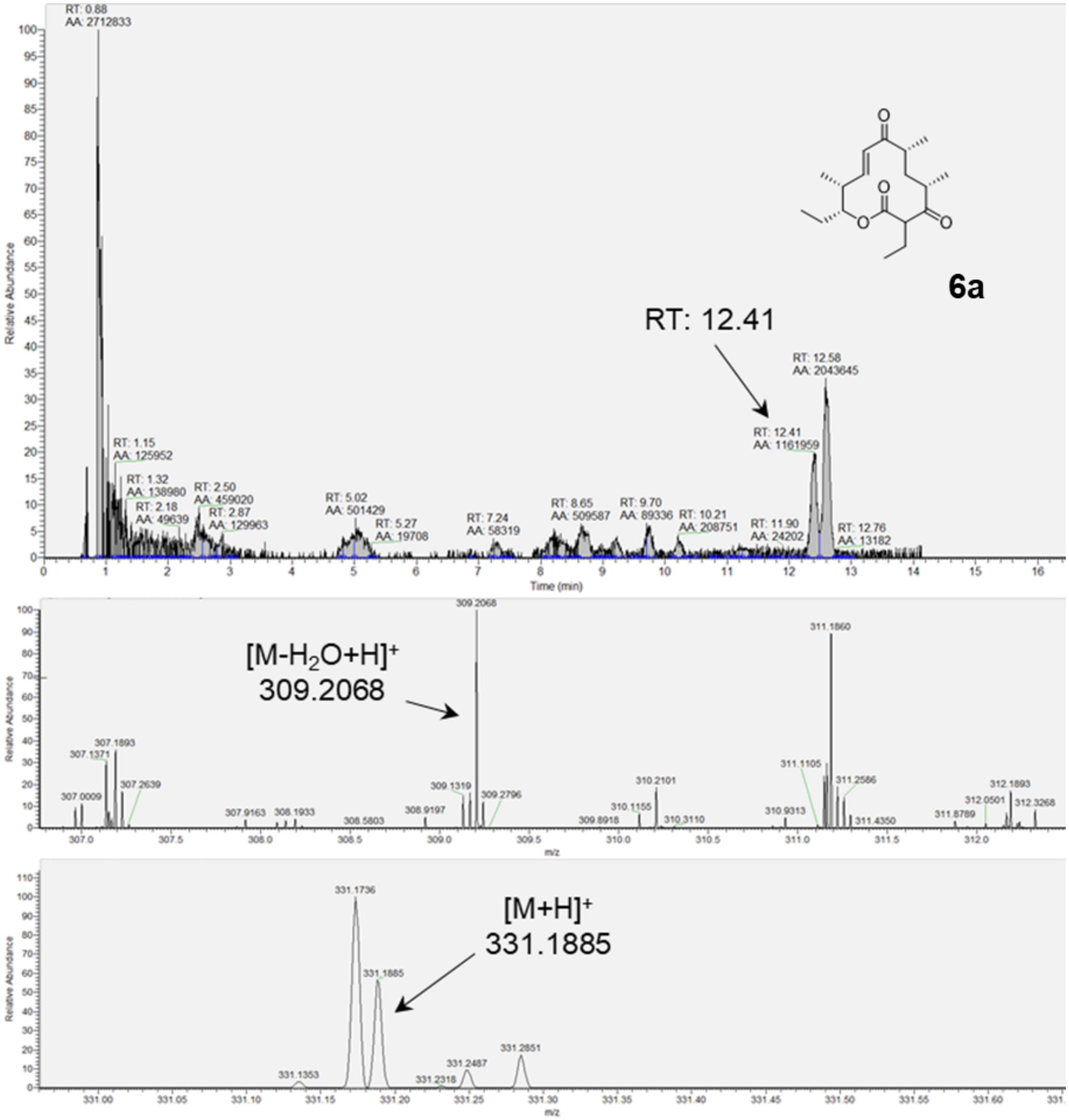

**b**

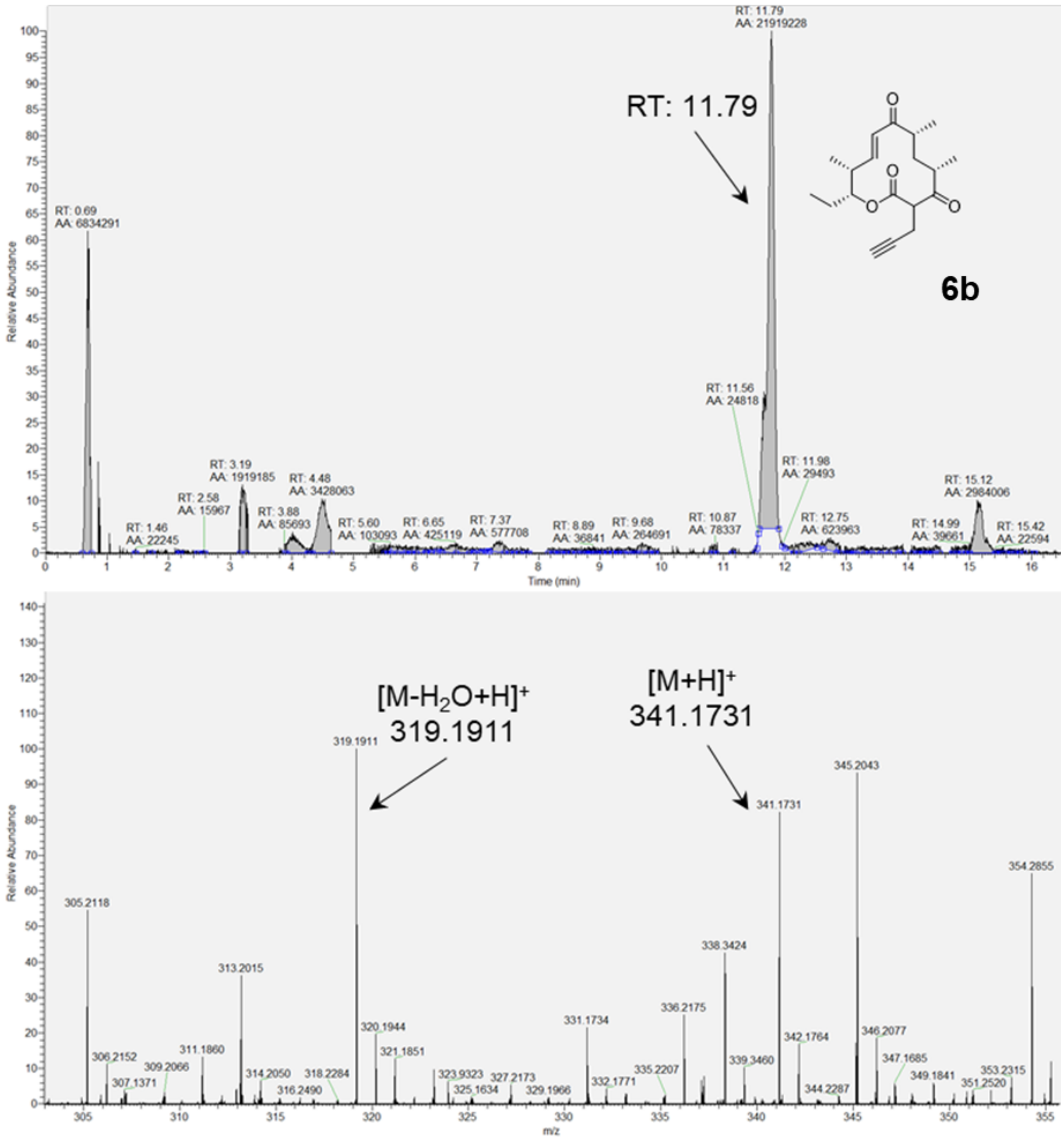
RT: 11.79
6b
Relative Abundance
Time (min)
[M-H2O+H]+
319.1911
[M+H]+
341.1731
345.2043
354.2855
305.2118
338.3424
313.2015
m/z

c

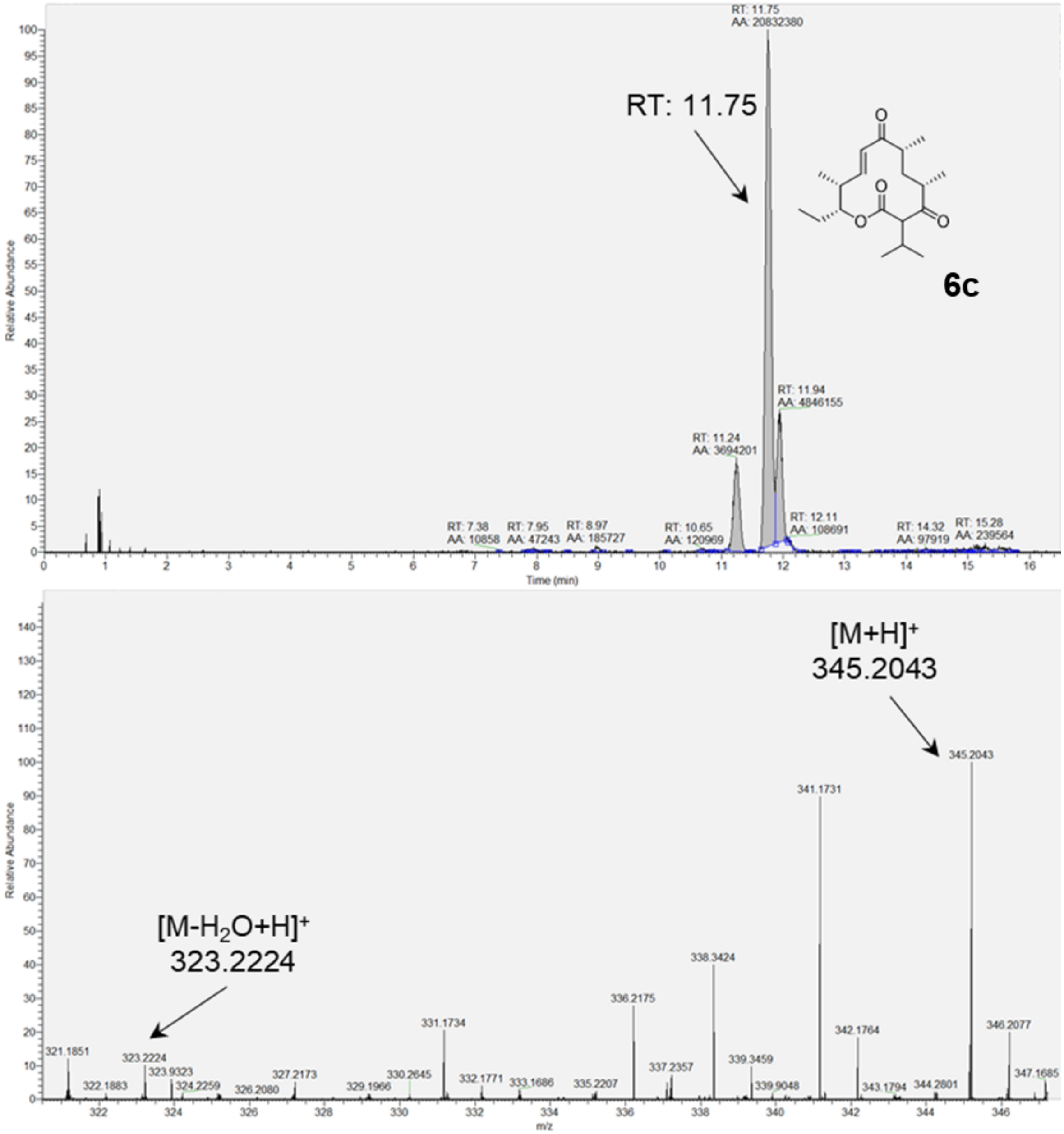

RT: 11.75
AA: 20832380
RT: 11.75
6c
RT: 11.94
AA: 4846155
RT: 11.24
AA: 3694201
RT: 7.38
AA: 10858
RT: 7.95
AA: 47243
RT: 8.97
AA: 185727
RT: 10.65
AA: 120969
RT: 12.11
AA: 108691
RT: 14.32
AA: 97919
RT: 15.28
AA: 239564
Relative Abundance
Time (min)
[M+H]+
345.2043
[M-H2O+H]+
323.2224
321.1851
322.1883
323.2224
323.9323
324.2259
326.2080
327.2173
329.1966
330.2645
331.1734
332.1771
333.1686
335.2207
336.2175
337.2357
338.3424
339.3459
339.9048
341.1731
342.1764
343.1794
344.2801
345.2043
346.2077
347.1685
Relative Abundance
m/z

d

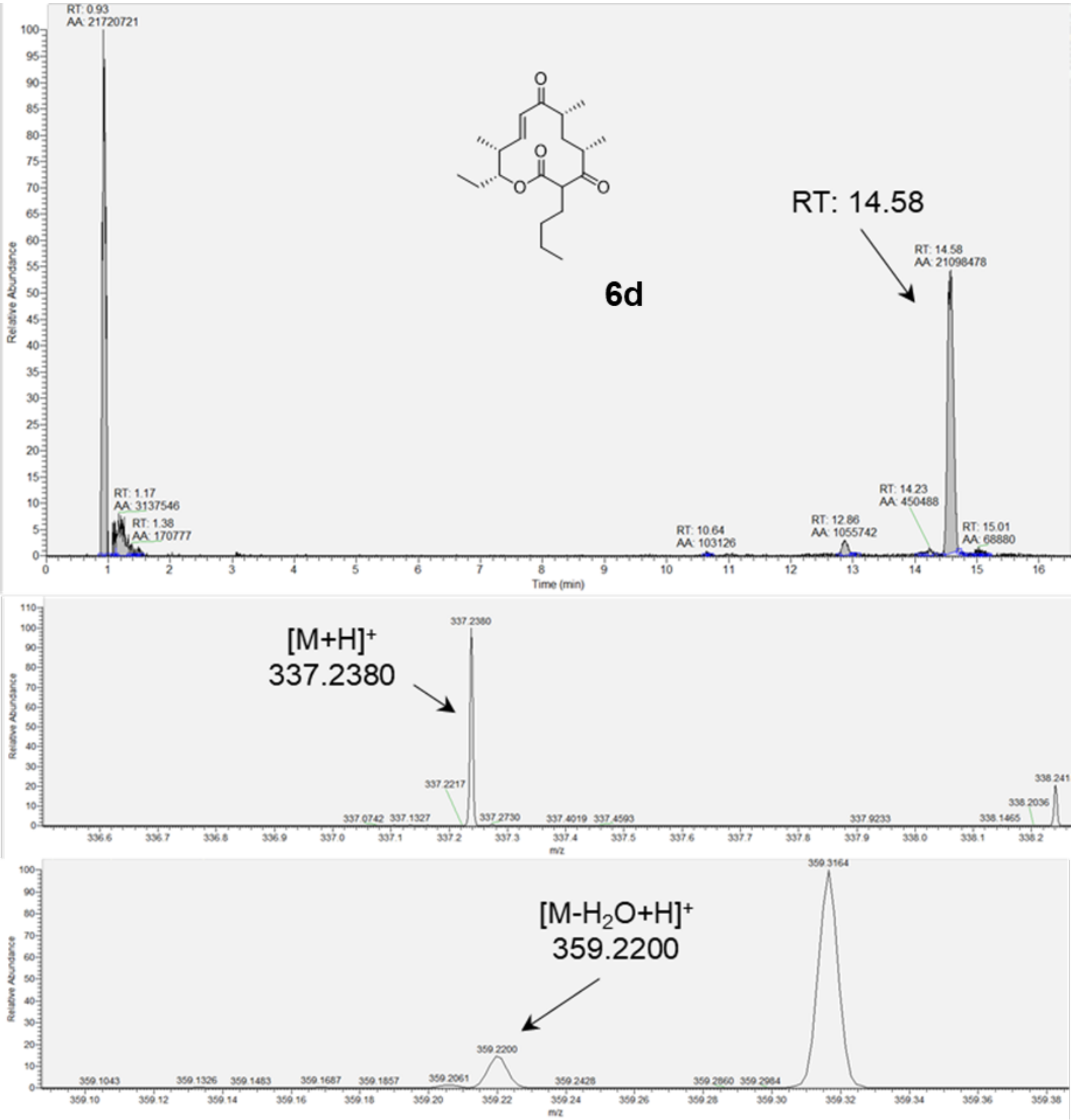

6d
RT: 14.58
RT: 0.93
AA: 21720721
RT: 1.17
AA: 3137546
RT: 1.38
AA: 170777
RT: 10.64
AA: 103126
RT: 12.86
AA: 1055742
RT: 14.23
AA: 450488
RT: 14.58
AA: 21098478
RT: 15.01
AA: 68880
Relative Abundance
Time (min)
[M+H]+
337.2380
m/z
[M-H2O+H]+
359.2200

**e**

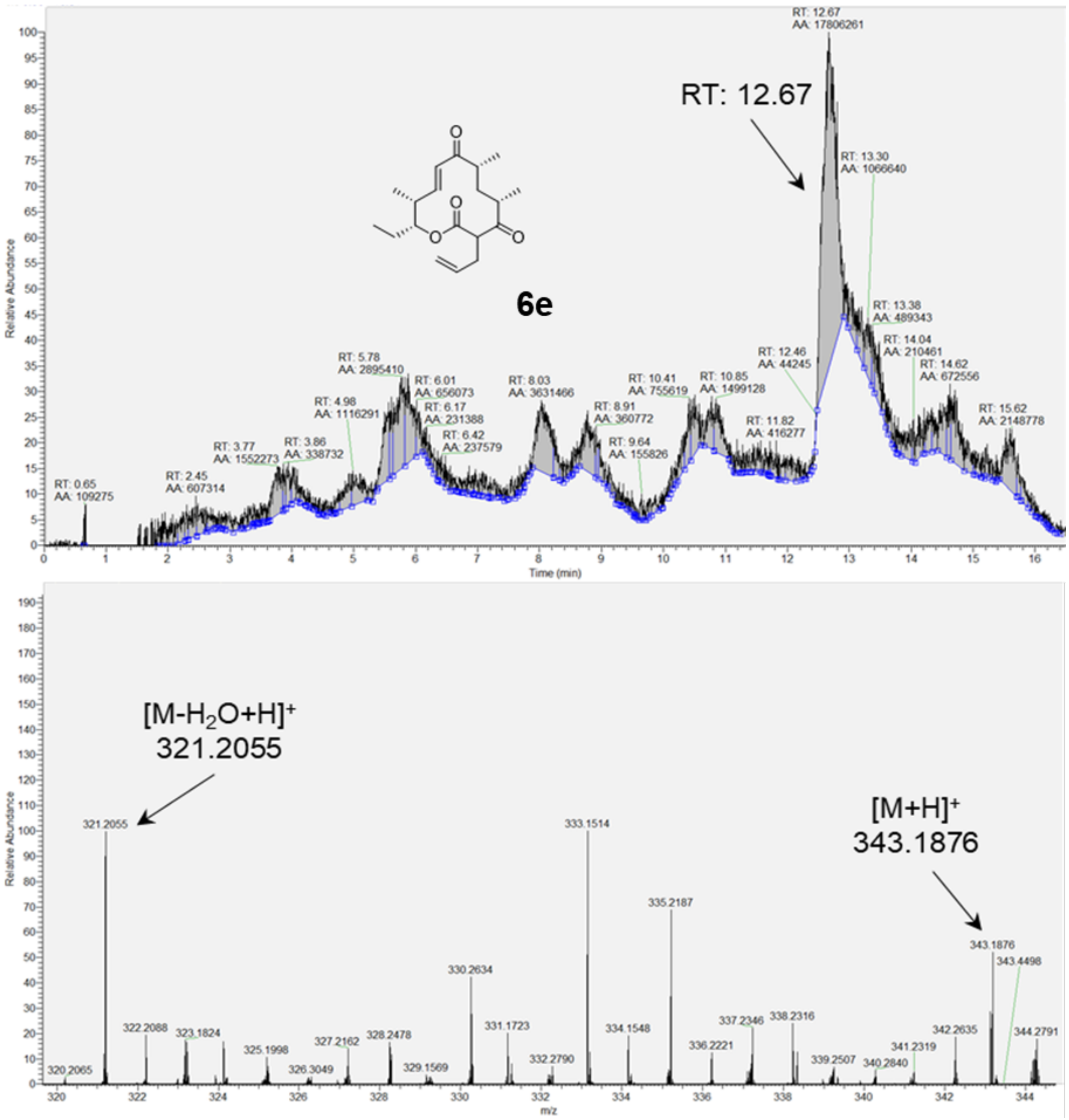

RT: 12.67
AA: 17806261
RT: 12.67
6e
Relative Abundance
Time (min)
RT: 0.65
AA: 109275
RT: 2.45
AA: 607314
RT: 3.77
AA: 1552273
RT: 3.86
AA: 338732
RT: 4.98
AA: 1116291
RT: 5.78
AA: 2895410
RT: 6.01
AA: 656073
RT: 6.17
AA: 231388
RT: 6.42
AA: 237579
RT: 8.03
AA: 3631466
RT: 8.91
AA: 360772
RT: 9.64
AA: 155826
RT: 10.41
AA: 755619
RT: 10.85
AA: 1499128
RT: 11.82
AA: 416277
RT: 12.46
AA: 44245
RT: 13.30
AA: 1066640
RT: 13.38
AA: 489343
RT: 14.04
AA: 210461
RT: 14.62
AA: 672556
RT: 15.62
AA: 2148778
[M-H2O+H]+
321.2055
[M+H]+
343.1876
Relative Abundance
m/z
320.2065
321.2055
322.2088
323.1824
325.1998
326.3049
327.2162
328.2478
329.1569
330.2634
331.1723
332.2790
333.1514
334.1548
335.2187
336.2221
337.2346
338.2316
339.2507
340.2840
341.2319
342.2635
343.1876
343.4498
344.2791

f

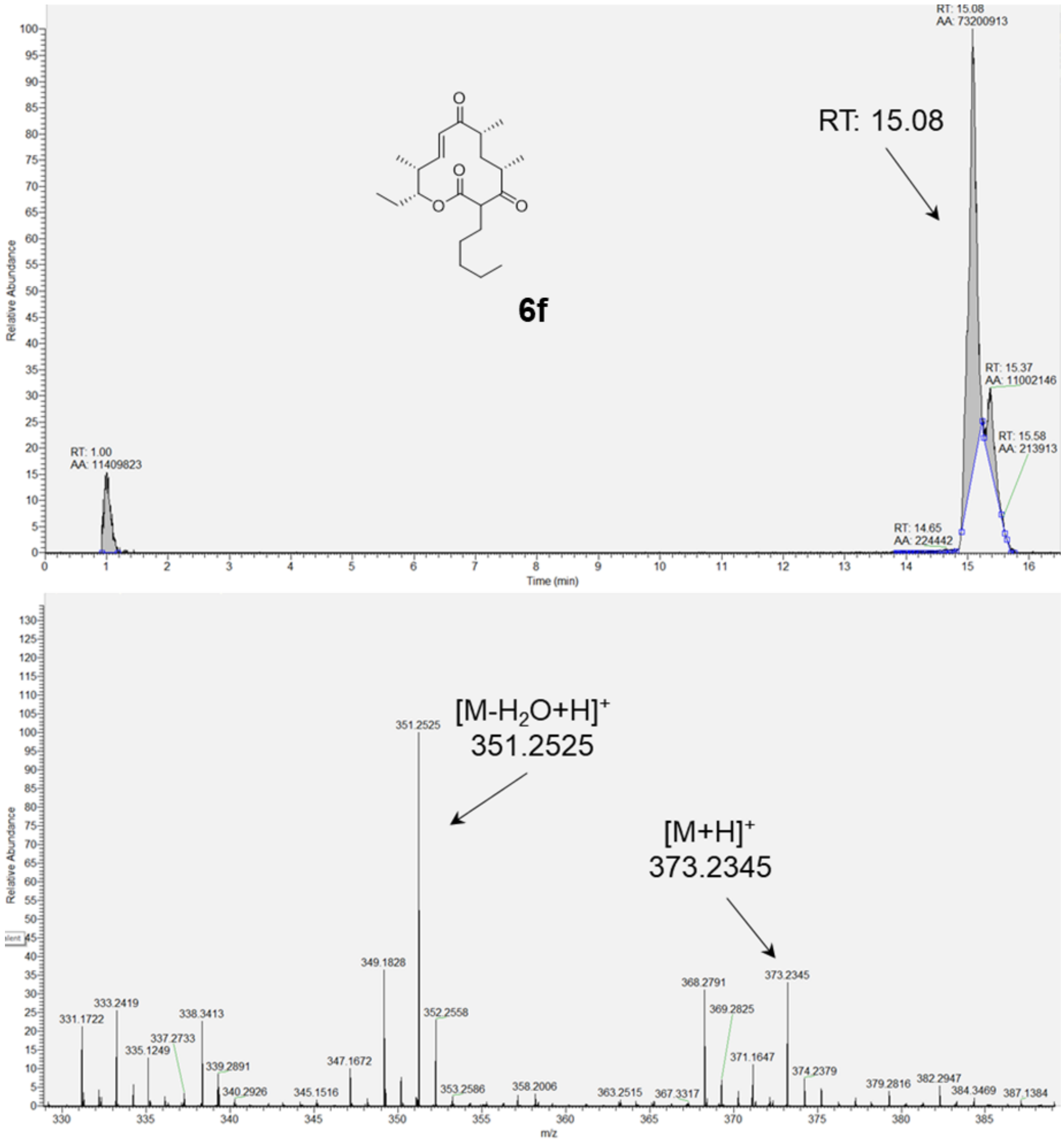

RT: 15.08
AA: 73200913
RT: 15.08
6f
RT: 15.37
AA: 11002146
RT: 15.58
AA: 213913
RT: 1.00
AA: 11409823
RT: 14.65
AA: 224442
Relative Abundance
Time (min)
[M-H2O+H]+
351.2525
[M+H]+
373.2345
351.2525
349.1828
373.2345
368.2791
369.2825
352.2558
333.2419
331.1722
338.3413
337.2733
335.1249
339.2891
340.2926
345.1516
347.1672
353.2586
358.2006
363.2515
367.3317
371.1647
374.2379
379.2816
382.2947
384.3469
387.1384
Relative Abundance
m/z

### Supplemental Methods

#### *Site-directed mutagenesis of Ery6TE*

All Ery6 and Er6TE site-directed mutants were constructed by 'round-the-horn' mutagenesis using the templates from **Supplementary Table S6** and the oligonucleotide sequences described **Supplementary Table S7**.[1] Each PCR contained: 5X GC Phusion DNA polymerase buffer (4 µL), DNAase free water (11.8 µL), forward/reverse primer mix (2 µL, 10 µM), template DNA (1 µL, 50 ng/µL), dNTPs (0.4 µL, 2.5 mM each dNTP), DMSO (0.6 µL), and Phusion High Fidelity DNA Polymerase (0.2 µL) (New England Biolabs, NEB). PCR cycling parameters: step 1) 98 °C, 60 s; step 2) 29x [a) 98 °C, 10 s; b) 72 °C, 5 min]; step 3) 72 °C, 5 min. Next, the PCR mixture was subjected to restriction enzyme digest with *Dpn*I to remove any remaining template DNA. The *Dpn*I reaction mixture contained: 10X CutSmart Buffer (2 µL), the PCR mixture (17 µL), and *Dpn*I (1 µL). The mixture was vortexed, centrifuged, and incubated for 3 h at 37 °C. The digested reaction mixture was then purified by agarose gel electrophoresis, with a total of 8 µl DNA eluted at the final step. The PCR product was then ligated using T4 DNA ligase in a reaction containing: 1 µl of the 10X T4 DNA ligase buffer, 8 µl of the *Dpn*I-treated and purified PCR product, and 1 µl of the T4 DNA ligase (overnight incubation at 16 °C). Each subsequent ligation reaction was transformed directly into *E. cloni* 10G electrocompetent cells (Lucigen). Individual transformants were sequenced to identify incorporation of mutant codons.

#### *Construction of Ery6TE motif chimeras*

Motif swaps of Ery6TE AT were constructed using Gibson assembly (NEB) and boundaries defined by the Keasling group.[2] The Gibson assembly mixtures were used to transform *E. coli* DH5α competent cells. Successful incorporation of mutations was confirmed by DNA sequencing of purified plasmids from single transformants. Each mutant plasmid was transformed into *E. coli* K207-3 competent cells and plated onto LB agar plates (50 µg $mL^{-1}$ kanamycin and 100 µg $mL^{-1}$ spectinomycin).

#### *Expression and purification of wild-type and mutant MatB*

The expression and purification of MatB T207G/M306I has been previously described.[3-5] Briefly, *E. coli* BL21(DE3) pLysS competent cells were transformed with plasmid. and positive transformants were selected on LB agar supplemented with 50 µg/mL kanamycin. A single colony was transferred to LB (3 mL) supplemented with kanamycin (50 µg/mL) and grown at 37 °C and 250 rpm overnight. The culture was used to inoculate LB media (1 L) supplemented with kanamycin (50 µg/mL). One liter culture was incubated at 37 °C and 250 rpm to an $OD_{600}$ of 0.6, at which time protein synthesis was induced by the addition of IPTG to a final concentration of 1 mM. After incubation at 18 °C and 200 rpm for 18 h, cells were collected by centrifugation at 5,000 *g* for 20 min and resuspended in 100 mM Tris-HCl pH 8.0 (20 mL) containing NaCl (300 mM) and then lysed by sonication. Following centrifugation at 10,000 *g*, the soluble extract was loaded onto a 1 mL HisTrap HP column (GE Healthcare, Piscataway, NJ) and purified by fast protein liquid chromatography using the following buffers: wash buffer [20 mM phosphate (pH 7.4) containing 0.5 M NaCl and 20 mM imidazole] and elution buffer [20 mM phosphate (pH 7.4) containing 0.5 M NaCl and 200 mM imidazole]. The purified protein was concentrated using an Amicon Ultra 30 kDa MWCO centrifugal filter (Millipore Corp., Billerica, MA) and stored as 10% glycerol stocks at -80 °C. Protein purity was verified by SDS-PAGE. Protein quantification was carried out using the Bradford Protein Assay Kit from Bio-Rad.

***Synthesis of acyl-CoAs by MatB***

The MatB-catalyzed synthesis of extender units **1** and **2a-f** has been previously described.[3-5] Briefly, reactions were performed in a 50 µL reaction mixture containing 100 mM sodium phosphate (pH 7), $MgCl_2$ (2 mM), ATP (12 mM), coenzyme A (8 mM), malonate or corresponding analog (16 mM) and wild-type or mutant MatB (10 µg) at 25 °C. Aliquots were removed after 3 h incubation, and quenched with an equal volume of ice-cold methanol, centrifuged at 10,000 *g* for 10 min, and cleared supernatants used for HPLC analysis on a Varian ProStar HPLC system. A series of linear gradients was developed from 0.1% TFA (A) in water to methanol (HPLC grade, B) using the following protocol: 0-32 min, 80% B; 32-35 min, 100% A. The flow rate was 1 mL/min, and the absorbance was monitored at 254 nm using Pursuit XRs C18 column (250 x 4.6 mm, Varian Inc.). To ensure complete conversion, the malonate analogue and the acyl-CoA product HPLC peak areas were integrated, and the conversion (%) calculated as a percent of the total peak area. Product elution times and LC-MS data were in complete agreement with that previous described.[3,4]

**Supplemental References**


1 Moore, S. *'Round-the-horn site-directed mutagenesis*, <https://openwetware.org/wiki/'Round-the-horn_site-directed_mutagenesis>

2 Yuzawa, S. *et al.* Comprehensive in vitro analysis of acyltransferase domain exchanges in modular polyketide synthases and its application for short-chain ketone production. *ACS Synth. Biol.* **6**, 139-147 (2017).

3 Koryakina, I. *et al.* Poly specific trans-acyltransferase machinery revealed via engineered acyl-CoA synthetases. *ACS Chem. Biol.* **8**, 200-208 (2013).

4 Koryakina, I., McArthur, J. B., Draelos, M. M. & Williams, G. J. Promiscuity of a modular polyketide synthase towards natural and non-natural extender units. *Org. Biomol. Chem.* **11**, 4449-4458 (2013).

5 Koryakina, I. & Williams, G. J. Mutant malonyl-CoA synthetases with altered specificity for polyketide synthase extender unit generation. *ChemBioChem* **12**, 2289-2293 (2011).

6 Crooks, G. E., Hon, G., Chandonia, J. M. & Brenner, S. E. WebLogo: A sequence logo generator. *Genome Res.* **14**, 1188-1190 (2004).
